# A Multiscale, Systems-level, Neuropharmacological Model of Cortico-Basal Ganglia System for Arm Reaching under Normal, Parkinsonian and Levodopa Medication Conditions

**DOI:** 10.1101/2021.02.10.430544

**Authors:** Sandeep Sathyanandan Nair, Vignayanandam Ravindernath Muddapu, V. Srinivasa Chakravarthy

## Abstract

The root cause of Parkinson’s disease (PD) is the death of dopaminergic neurons in Substantia Nigra pars compacta (SNc). The exact cause of this cell death is still not known. Loss of SNc cells manifest as the cardinal symptoms of PD, including tremor, rigidity, bradykinesia, and postural imbalance. To investigate the PD condition in detail and understand the link between loss of cells in SNc and PD symptoms, it is important to have an integrated multiscale computational model that can replicate the symptoms at the behavioural level by evoking the key cellular and molecular underlying mechanisms that contribute to the pathology. In line with this objective, we present a multiscale integrated model of cortico-basal ganglia motor circuitry for arm reaching task, incorporating a detailed biophysical model of SNc dopaminergic neuron. Earlier researchers have shown that fluctuations in dopamine (DA) signals are analogous to reward/punishment signals, thereby prompting application of concepts from reinforcement learning (RL) to modelling the basal ganglia system. In our model, we replace the abstract representations of reward with the realistic variable of extracellular DA released by a network of SNc cells and incorporate it with the RL-based behavioural model, which simulates the arm reaching task. Our results showed that as SNc cell loss increases, the percentage success rate to reach the target decreases, and average time to reach the target increases. With levodopa (L-DOPA) medication, both the success rate and the average time to reach the target improved significantly. The proposed model also exhibits how differential dopaminergic axonal degeneration in basal ganglia results in various cardinal symptoms of PD as manifest in reaching movements. From the model results, we were able to show the side effects of L-DOPA mediation, such as wearing off and peak dosage dyskinesias. Moreover, from the results, we were able to predict the optimum dosage for varying degrees of cell loss and L-DOPA medication. The proposed model has a potential clinical application where drug dosage can be optimized as per patient characteristics. We conclude that our model presents a realistic and efficient way of simulating the PD pathology conditions and the effect of levodopa medication, thereby giving a reliable indicator towards the optimization of the drug dosage.

## 1. INTRODUCTION

Parkinson’s Disease (PD) is the second most prominent neurodegenerative disease after Alzheimer’s (Gonzalez-Rodriguez et al., 2020; Marino et al., 2020; Muddapu and Chakravarthy, 2021). The onset of the disease is characterized by shaky movements, rigidity of joints, unregulated movements, and even loss of smell (Morley and Duda, 2010; Fullard et al., 2017; Armstrong and Okun, 2020; Balestrino and Schapira, 2020; Goldman and Guerra, 2020; Marino et al., 2020). The major cause of this disease is the death of dopaminergic neurons in Substantia Nigra pars compacta (SNc) (Michel et al., 2016; Surmeier, 2018; Muddapu et al., 2020a). Dopamine (DA) deficiency due to SNc cell loss manifest as the cardinal PD symptoms that include tremor, rigidity, bradykinesia and postural imbalance (Bereczki, 2010; Poewe et al., 2017; Balestrino and Schapira, 2020). Epidemiological data from the United States alone indicates that there has been an exponential growth of people suffering from PD over the last few decades (Dorsey et al., 2018; Marras et al., 2018). However, the exact cause of this cell death is still not known. Various lines of investigation, experimental and computational, are in progress and hopefully we will be able to narrow down the roots of this disease (Pissadaki and Bolam, 2013; Pacelli et al., 2015; Fu et al., 2018; Giguère et al., 2019; Muddapu et al., 2019, 2020a, 2020b; Anilkumar et al., 2020; Gonzalez-Rodriguez et al., 2020; Muddapu and Chakravarthy, 2021). Understanding the cause and effect relationship between the underlying pathology and symptoms of any neurological disease has fundamental challenges since the roots of the disease are at molecular and cellular level while the symptoms are seen at the behavioural level (Bakshi et al., 2019). Hence it is important to have a multi-scale model that spans molecular mechanisms to behavioral outputs. With this motivation in mind, we present a computational model that relates DA deficiency in PD to motor symptoms in ON and OFF conditions of medication. As an example of drug action, we simulate the effect of levodopa (L-DOPA) drug administration in our model.

### Basal Ganglia & Dopamine

Dopaminergic input from the SNc neurons modulates the DA receptors present in the striatal neurons, the input nuclei of the basal ganglia (BG) differentially. The striatum (STR) consists of the D1 and D2-type expressing medium spiny neurons (MSN) that project via two different pathways. D1-MSN neurons project along the direct pathway, D2-MSNs project along the indirect pathway. The direct pathway projects directly to the output nuclei, globus pallidus interna (GPi) and Substantia Nigra pars reticulata (SNr), whereas the indirect pathway projects to the output nucleus, GPi, via globus pallidus externa (GPe) and subthalamic nucleus (STN). DA release from SNc neurons maintains the balance between activation of direct and indirect pathways. In order to understand the effect of DA deficiency as in PD conditions, or the mechanism of DA replenishment by administration of L-DOPA, we need to understand DA synthesis, uptake and release (Chakravarthy and Moustafa, 2018; Muddapu and Chakravarthy, 2021).

In this paper, we present a multiscale model of the cortical basal ganglia system to simulate arm reaching movements under conditions of PD pathology and L-DOPA medication. At the lowest level, the intracellular molecular pathways of SNc cells are modelled so as to capture dopamine synthesis, uptake and release. At the next level, the BG circuitry is modelled using rate coded neurons which is cast within the reinforcement learning framework with striatum acting as the neural correlate for critic and the direct and indirect pathways facilitating exploitation and exploration, respectively. At the highest level, arm reaching movements are modelled by a two-link arm model driven by a sensory-motor cortical loop.

This article is organized into multiple sections. Section 2 describes the model architecture, equations and methods. Here we discuss various functional loops that constitute in the model and how they are interconnected. This section also covers the integration of the pharmacological intervention. In section 3 we showcase the results from the model starting with training the model, simulating the behavior of a control subject, replicating the PD ON condition and some of the cardinal symptoms, assessing the performance in terms of reaching time and verifying the effect of L-DOPA therapeutic intervention. The model results also gave an indicator of how to optimize the drug dosage. Section 4 discusses the simulation results in detail and based on that the conclusion derived is updated in section 5. Section 6 presents the potential future scope.

## 2. MATERIALS AND METHODS

The proposed multiscale cortico-basal ganglia (MCBG) model was able to simulate the arm reaching in normal and Parkinsonian conditions which includes some of the cardinal symptoms of PD (Figure 1). In addition, the effect of L-DOPA medication on arm reaching in PD condition was simulated (Figure S1).

**Figure 1:**
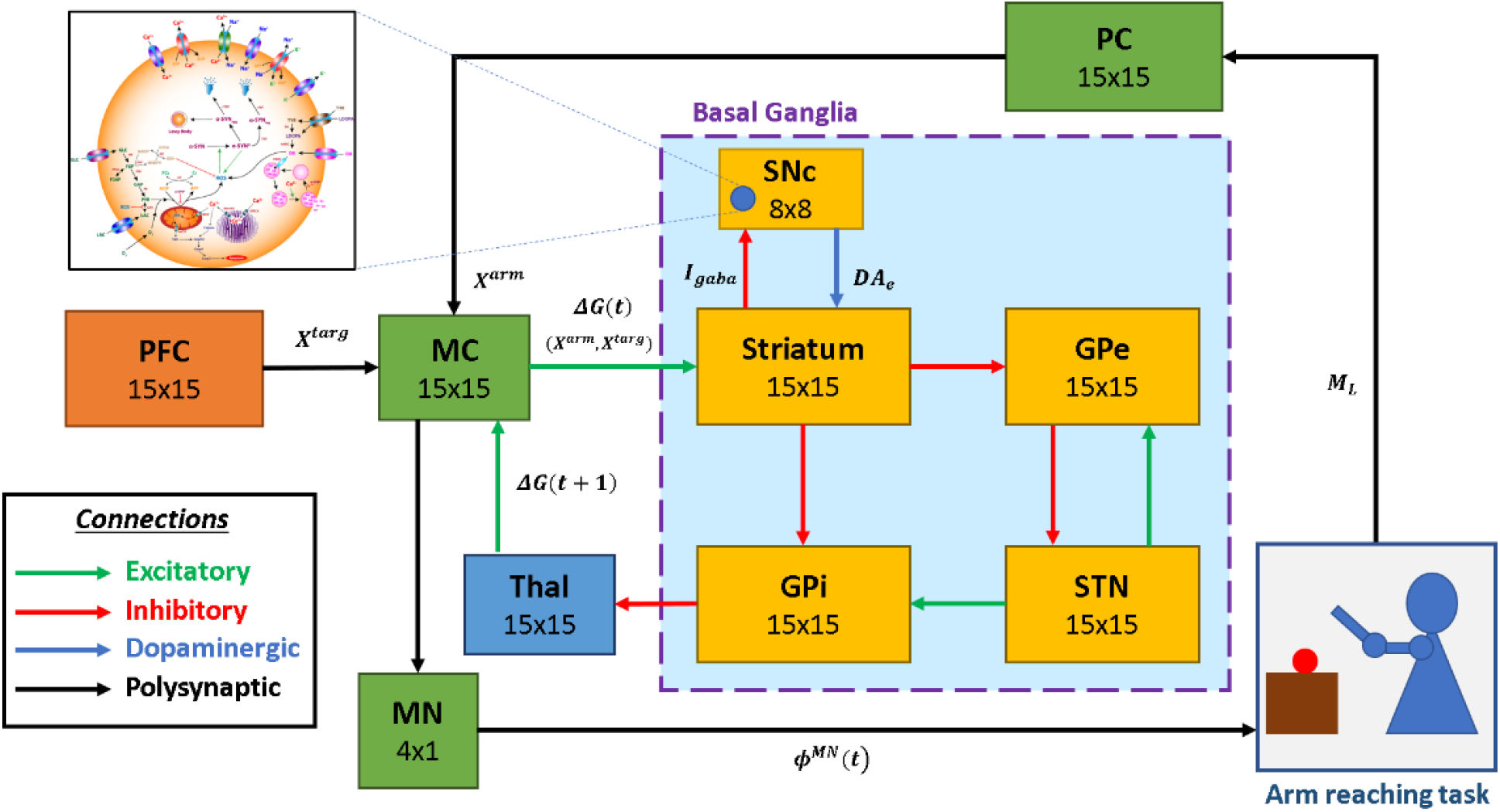
Model architecture of multiscale cortico-basal ganglia model for arm reaching. SNc, substantia nigra pars compacta; GPe, globus pallidus externa; GPi, globus pallidus interna; STN, subthalamic nucleus; Thal, thalamus; MC, motor cortex; MN, motor neuron; PC, proprioceptive cortex; PFC, prefrontal cortex. X^targ^, the target position; X^arm^, the current arm position; ϕ^MN^, the motor neuron activations; M_L_, muscle lengths; I_gaba_, inhibitory GABAergic current; DA_e_, extracellular dopamine; ΔG(t), the MC output; ΔG(t + 1), the BG-derived activity of thalamus.

The proposed model can be broadly described in three parts. i) Outer loop – motor-sensory loop, ii) Inner loop – cortico-basal ganglia loop and iii) Central loop – nigrostriatal loop (Figure 2). The outer loop consists of motor cortex (MC), motor neurons (MNs), arm, proprioceptive cortex (PC) and prefrontal cortex (PFC). The inner loop consists of MC, thalamus, and BG nuclei comprised of striatum, GPi, GPe, and STN. The central loop consists of striatum and SNc, which plays an important role in simulating PD conditions, where nigrostriatal and nigrosubthalamic pathways affected by SNc cell loss. For L-DOPA medication, a pharmacokinetic module was formulated where input will be L-DOPA dosage given to the PD patient and output will be the amount of DA released in striatum during the medication. The subsequent sections describe the dynamics involved in each of these three loops.

**Figure 2:**
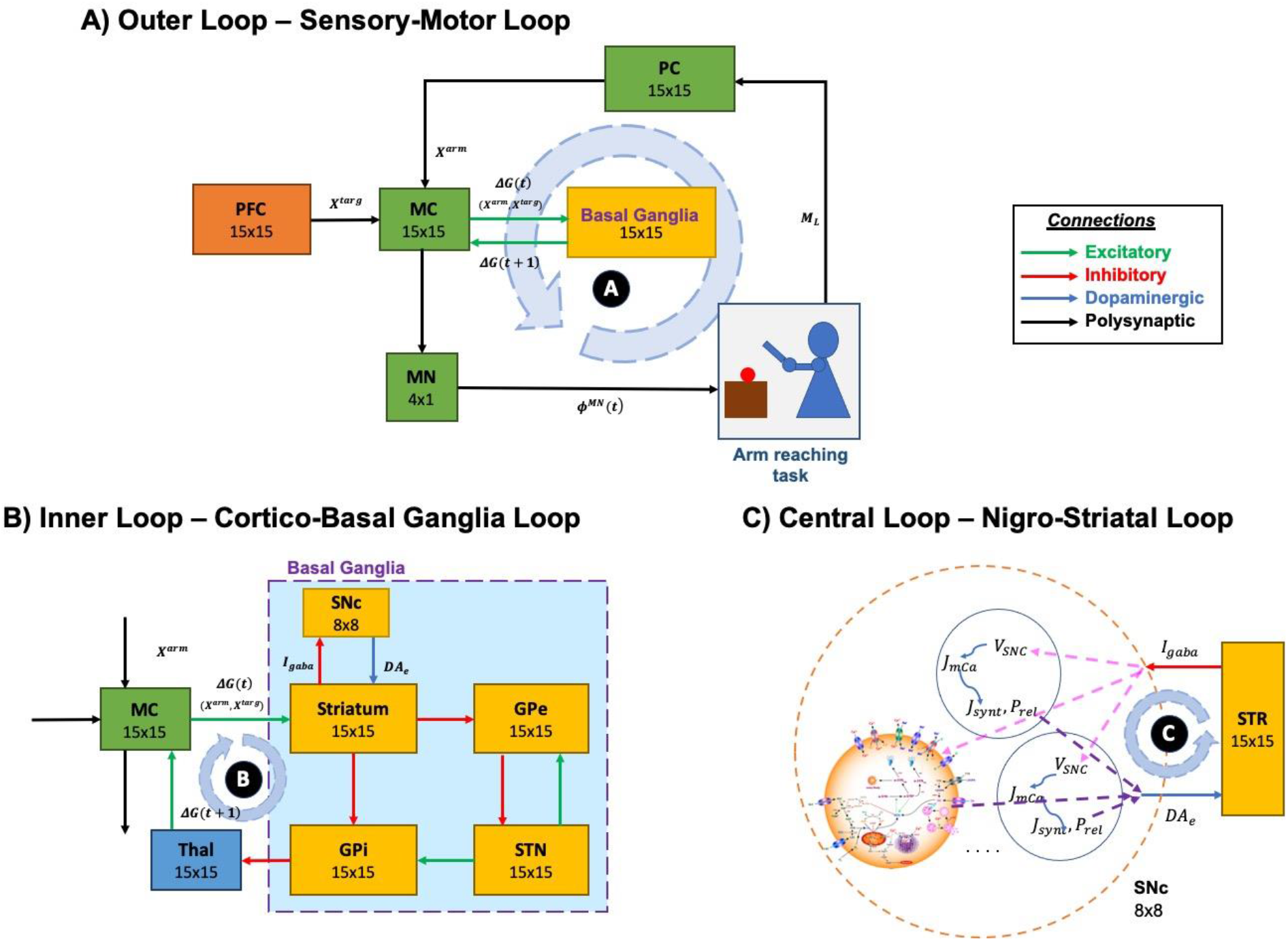
Different structural and functional loops of the proposed multiscale cortico-basal ganglia model. A) Outer loop – sensory-motor loop B) Inner loop – cortico-basal ganglia loop C) Central loop – nigrostriatal loop. SNc, substantia nigra pars compacta; GPe, globus pallidus externa; GPi, globus pallidus interna; STN, subthalamic nucleus; STR, striatum; Thal, thalamus; MC, motor cortex; MN, motor neuron; PC, proprioceptive cortex; PFC, prefrontal cortex. X^targ^, the target position; X^arm^, the current arm position; ϕ^MN^, the motor neuron activations; M_L_, muscle lengths; I_gaba_, inhibitory GABAergic current; DA_e_, extracellular dopamine; ΔG(t), the MC output; ΔG(t 1), the BG-derived activity of thalamus; V_sNc_, the voltage membrane of SNc neuron; J_m,Ca_, the calcium flux of SNc neuron as a function of V_sNc_; J_synt_, the dopamine synthesis flux as function of calcium; P_rel_, the probability release of dopamine extracellularly as a function of calcium.

### 2.1. Outer Loop – Sensory-Motor Loop

The functional pathway of the outer loop is shown in Figure 2A. The outer loop consists of a two-link arm model driven by MNs. MNs receive motor commands from MC. The end effector position of the arm is sensed by PC and it forwards the signal to MC, which receives signals from PFC and BG. MC issues the motor commands based on the integration of incoming signals.

#### 2.1.1. Arm Model

The kinetic model of the two-joint arm simulates the movement of the arm in two-dimensional space (Izawa et al., 2004; Zadravec and Matjačić, 2013) (Figure S2). Each joint (shoulder and elbow) is controlled by an agonist (Ag) and antagonist (An) muscle pair where the shoulder joint is controlled by anterior deltoid (shoulder flexor, *M*_1_) and posterior deltoid (shoulder extensor, *M*_2_) and elbow joint is controlled by brachialis (elbow flexor, *M*_3_) and triceps brachii (elbow extensor, *M*_4_) (Jagodnik and van den Bogert, 2010). The activations to these muscle groups (*ϕ*^*MN*^) are transformed into joint angles for both shoulder and the elbow as follows,

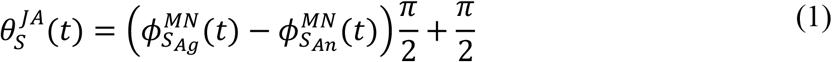

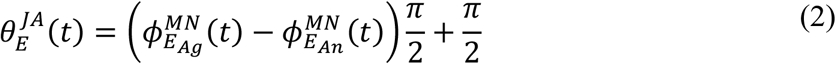

where, 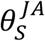 and 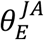 are the joint angles of shoulder and elbow with respect to the x-axis (Figure S1) and shoulder length (*l*_*s*_), respectively in two-dimensional space, 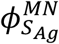 is the muscle activation of shoulder agonist muscle, 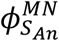 is the muscle activation of shoulder antagonist muscle, 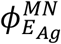 is the muscle activation of elbow agonist muscle, and 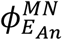 is the muscle activation of elbow antagonist muscle.

The coverage of the arm in two-dimensional space is controlled by these joint angles. The joint angles are used to calculate the muscle lengths for both shoulder and elbow as given below.

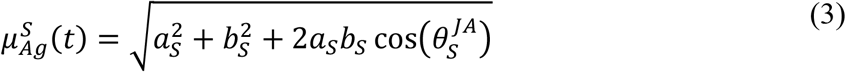

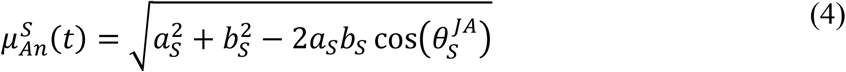

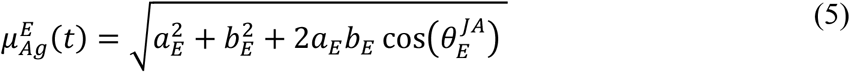

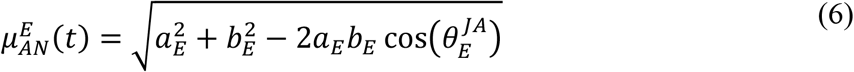

where, 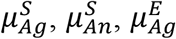, and 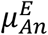 are the agonist and antagonist muscle lengths of shoulder and elbow, respectively, *a*_*s*_ is the distance between shoulder joint center and *M*_1_ or *M*_2_ moment lever, *b*_*s*_ is the distance between shoulder joint center and *M*_1_ or *M*_2_ moment lever, *a*_*E*_ is the distance between elbow joint center and *M*_3_ or *M*_4_ moment lever, and *b*_*E*_ is the distance between elbow joint center and *M*_3_ or *M*_4_ moment lever.

Using these muscle lengths in the form of a four-dimensional vector 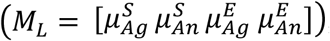 a sensory (proprioceptive) map of the arm was generated. The end effector position of the arm 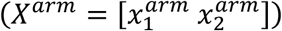 in the two-dimensional space is calculated as,

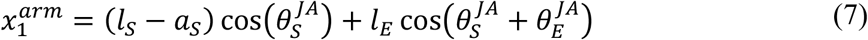

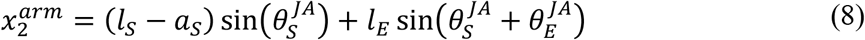

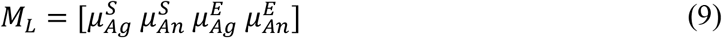

where, 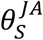 and 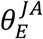 are the joint angles of shoulder and elbow with respect to the x-axis (Figure S1) and shoulder length (*l*_*s*_), respectively in two-dimensional space, *l*_*s*_ is the distance between the shoulder joint center (*s*) and elbow joint center (*E*), *l*_*E*_ is the distance between the elbow joint center (*E*) and end effector (*H*), *a*_*s*_ is the distance between shoulder joint center and *M*_1_ or *M*_2_ moment lever, 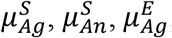, and 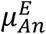 are the agonist (*M*_1_ or *M*_3_) and antagonist (*M*_2_ or *M*_4_) muscle lengths of shoulder and elbow, respectively.

#### 2.1.2. Proprioceptive Cortex

PC is modeled as self-organizing map (SOM) (Kohonen, 2001) of size *N*_*PC*_ *x N*_*PC*_ where sensory map of the arm was generated. Using muscle length vector (*M*_*L*_(*t*)) from the arm model (Eq. 9) as a feature vector, PC SOM was trained. The activation of a single node (*i, j*) in the PC SOM is given as,

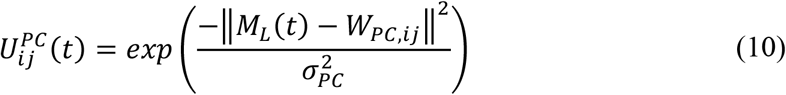

where, *W*_*PC,i*_ is the weight of the connection between the muscle length vector and *i*^*th*^ neuron of the two-dimensional PC network, *M*_*L*_ is the muscle length vector and *σ*_*PC*_ is the width of the Gaussian response of PC SOM.

#### 2.1.3. Prefrontal Cortex (PFC)

PFC encodes the goal position where, in real time, the goal information is formed using the visual sensory feedback, which is passed on to the frontal areas. In our current model, we fix the goal or target position and denote it by *X*^*targ*^. The motor command initially is driven by the PFC as the PFC specifies the goal to be reached. Similar to the PC, the PFC SOM is trained using target position vector as a feature vector. The input features of the PFC are the spatial locations where the arm can possibly reach in the two-dimensional space. The target locations produce the activation in the PFC network is given as.

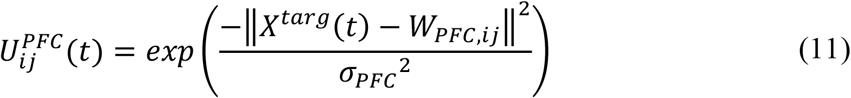

where, *W*_*PFC,ij*_ is the weight of the connection between the target position vector and (*i, j*)^*th*^ neuron of the two-dimensional PFC network, *X*^*targ*^ is the target position and *σ*_*PFC*_ is the width of the Gaussian response of PFC SOM.

#### 2.1.4. Motor Cortex

MC is modeled as a combination of SOM and continuous attractor neural network (CANN) (Trappenberg, 2011) of size *N*_*MC*_ *x N*_*MC*_. This type of architecture of MC is used to account for two distinct characteristics of cortical areas *viz*., low dimensional representation of input space and dynamics based on the connectivity in these cortical regions. A dynamic model like CANN is employed to facilitate the integration of multiple afferent inputs received from the PC, the BG and PFC. The output activity of the MC CANN (*G*_*MC*_) is defined by,

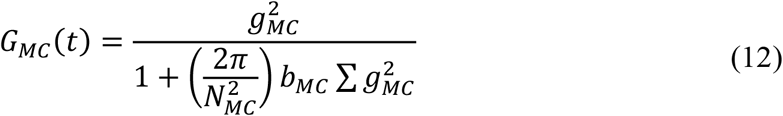

where, *g*_*MC*_ is the internal state of MC CANN, *N*_*MC*_ is the size of MC network, *b*_*MC*_ is the constant term.

The internal state of the MC CANN (*g*_*MC*_) is given by,

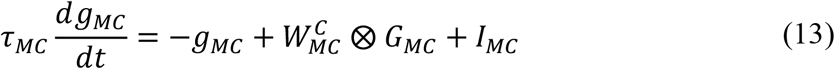

where, 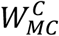 is the weight kernel representing lateral connectivity in MC CANN, which determines the local excitation/global inhibition dynamics, *G*_*MC*_ is the output activity of MC CANN, *I*_*MC*_ is the total input coming into MC CANN from PC, PFC and BG and ⨂ represents the convolutional operation.

##### 2.1.4.1. Lateral Connections in MC

The lateral connectivity in the MC CANN is characterized by short range (local) excitation and long range (global) inhibition whose dynamics are defined by the weight kernel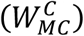 is given by,

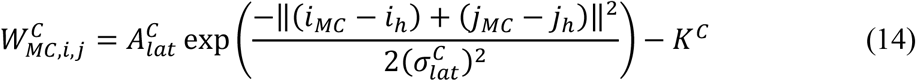

where, [*i*_*MC*_, *j*_*MC*_] are the location of the nodes in MC, [*i*_*h*_, *j*_*h*_] corresponds to the central node, 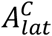 is the strength of the excitatory connections, *K*^*C*^ is the global inhibition constant and 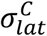 is the radius of the excitatory connections.

##### 2.1.4.2. Total Inputs into MC

The total input (*I*_*MC*_) coming into MC CANN from PC (information about current position of the arm), PFC (information about target position) and BG (error feedback signal) is given by,

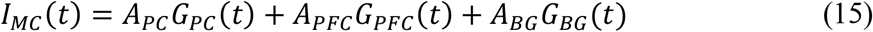

where, *A*_*PC*_, *A*_*PFC*_, *A*_*BG*_ are the respective gains of PC, PFC and BG, *G*_*PC*_, *G*_*PFC*_, *G*_*BG*_ are the output activities of PC-derived SOM part of MC, PFC-derived activation part of MC and BG-derived network activity of thalamus.

PC activity is used to generate low level feature maps in MC using SOM algorithm. The activation of the (*i, j*)^*th*^node in the SOM part of the MC (*G*_*PC,ij*_) is given as,

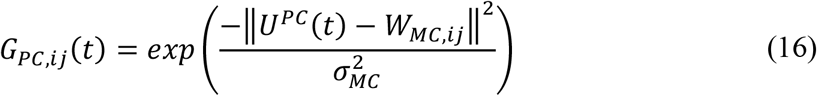

where, *U*^*PC*^ is the output activity of PC SOM network, *W*_*MC,i*_ is the weight of the connection between the PC SOM network and *i*^*th*^ neuron of the two-dimensional MC SOM network, and *σ*_*MC*_ is the width of the Gaussian response of MC SOM.

The input from PFC to MC (*G*_*PFC*_) is the product of weight matrix (*W*_*PFC*→*MC*_) and the output activity of PFC SOM is given by,

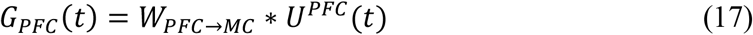

where, *U*^*PFC*^ is the output activity of PFC SOM network, *W*_*PFC*→*MC*_ is the weight matrix between PFC and MC.

In an earlier line of modelling studies, we have shown that the classical Go-NoGo interpretation of the functional anatomy of the BG must be expanded to Go-Explore-NoGo, in view of the putative role of the Indirect Pathway in exploration (Sridharan et al., 2006; Chakravarthy and Balasubramani, 2014). This series of models had resulted in the so-called Go-Explore-NoGo policy, that refers to a stochastic hill climbing performed on the value function computed inside the BG (Magdoom et al., 2011). When the arm reaches the target (ε < 0.1), the connections between the PFC and MC are trained by,

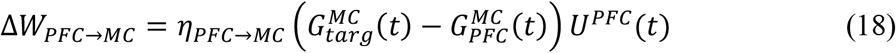

where, *η*_*PFC*→*MC*_ is the learning rate between PFC and MC, 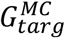 is the MC activation required for the arm to reach the target, and 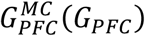 is the MC activation due to PFC.

#### 2.1.5. Motor Neurons

The output activity of MC CANN projects to the MN layer which consists of four MNs that drives four muscles of the arm whose activation is given by,

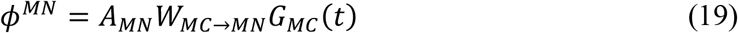

where, *A*_*MN*_ is the gain of MN, *W*_*MC*→*MN*_ is the weight matrix between MC CANN and MN layer, and *G*_*MC*_ is the output activity of MC CANN.

To close the sensory-motor loop, we perform a comparison with the initial activation to the MN layer that was used to produce desired activation 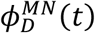. The weights between the MN and MC are trained in a supervised manner by comparing the network derived MN activation *ϕ*^*MN*^ (*t*) to the desired activation 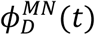. This gives a loop which is consistent in mapping the external arm space to the neuronal space and vice. The connection between MC and MN is trained by,

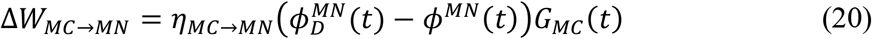

where, *η*_*MC*→*MN*_ is the learning rate between MC and MN, 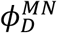 is the desired MN activation required for the arm to reach the target and *ϕ*^*MN*^ is the network-derived MN activation due to MC, and *G*_*MC*_ is the output activity of MC CANN. The training schema for the outer loop (sensory-motor loop) is described in section S3 of the supplementary information.

### 2.2. Inner Loop – Cortico-Basal Ganglia Loop

The functional pathway of the inner loop is shown in Figure 2B. The inner loop consists of MC, BG and thalamus. MC receives information from BG via thalamus. MC sends information to BG based on the integration of incoming signals received from PFC (target goal position, *X*^*targ*^), PC (current end effector position of the arm, *X*^*arm*^) and BG (via thalamus, error feedback signal, *G*_*BG*_).

#### 2.2.1 Basal Ganglia

BG consists of the striatum, GPe, GPi, STN and SNc. The output signal from BG provides the necessary control for the arm to reach the target by modulating the MC activity. The output of the MC is as given in Eq. 12.

##### 2.2.1.1 Value Computation and Stochastic Hill Climbing

The signal from the PC contains information about the current end effector position of the arm (*X*^*arm*^) whereas the signal from PFC contains the target goal position (*X*^*targ*^). These two signals are combined in the BG to form a value function, *V*^*arm*^(*t*), that represents the error between the desired and the actual positions of the hand as,

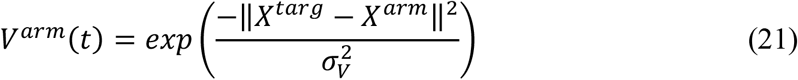

where, *X*^*targ*^ is the target goal position, *X*^*arm*^ is the current end effector position of the arm, *σ*_*V*_ is the spatial range over which the value function is sensitive for that particular target.

The output of the BG performs a stochastic hill climbing over the value function (Chakravarthy and Moustafa, 2018; Narayanamurthy et al., 2019) and drives the MC to facilitate the arm in reaching the target. The value difference (*δ*_*V*_) which is computed by comparing the current and previous values is given as,

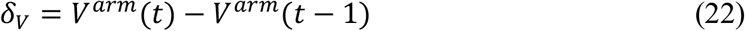

where, *V*^*arm*^(*t*) is the current value and *V*^*arm*^(*t* − 1) is the previous value.

Based on this value difference signal (*δ*_*V*_), the striatum will send the inhibitory GABAergic current (*I*_*gaba*_) to the SNc neurons while the SNc neurons will in turn release dopamine into the extracellular space (*DA*_*e*_), which is absorbed by the striatum. *DA*_*e*_ is transformed into 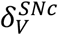. 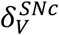 modulates the selection of direct and indirect pathways in the BG.

The dynamics between the striatum and the SNc is described in greater detail in the subsequent section, ‘The Central Loop’.

##### 2.2.1.2 Action Selection

###### Striatum

The resultant 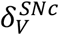 acts as a modulatory signal on D1R-MSNs and D2R-MSNs of the striatum, which process the input signal, Δ*G*_*MC*_(*t*), and send outputs *y*_*D*1_ & *y*_*D*2_ via direct and indirect pathways, respectively.

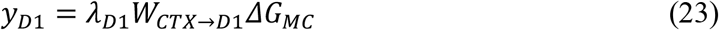

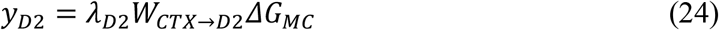

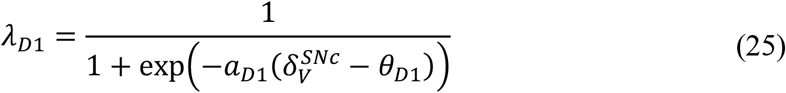

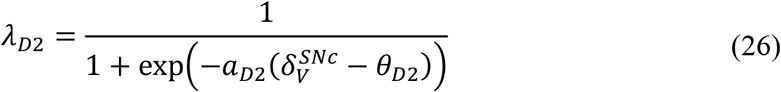

where, *λ*_*D*1_ and *λ*_*D*2_ represent the effect of dopamine (value difference) on the D1 and D2 MSNs, respectively, *W*_*CTX*→*D*1_ and *W*_*CTX*→*D*2_ represent connections between cortex and D1 MSNs and cortex and D2 MSNs, respectively, *ΔG*_*MC*_ is the output activity of MC, 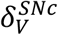 is the SNc-derived value difference, *θ*_*D*1_ and *θ*_*D*2_ are the thresholds of the D1 and D2 MSNs, respectively, *a*_*D*1_ and *a*_*D*2_ are the sigmoidal gains of the D1 and D2 MSNs, respectively. Since *a*_*D*1_ *=* −*a*_*D*2_, the activation of direct and indirect pathways depends on the 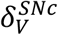 such that when 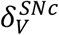 is positive (negative) the direct (indirect) pathway is selected.

###### STN-GPe Subsystem

In the indirect pathway, D2 MSNs of the striatum project to the GPe, where *y*_*D*2_ influences GPe neural dynamics, which in turn influences STN neural dynamics. STN-GPe forms a loop with inhibitory projections from GPe to STN and excitatory projections from STN to GPe. Such excitatory-inhibitory pairs of neuronal pools have been shown to exhibit limit cycle oscillations (Gillies et al., 2002) which was modeled as coupled Van der Pol oscillator (Kawahara, 1980). The dynamics of STN-GPe system is defined as,

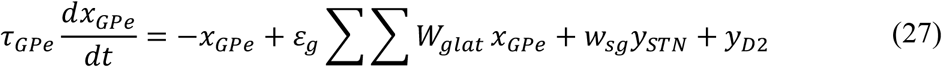

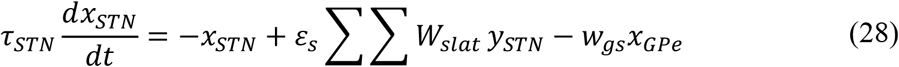

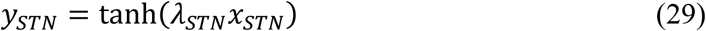

where, *x*_*GPe*_ and *x*_*sTN*_ are the internal states of GPe and STN neurons, respectively, *y*_*sTN*_ is the output of STN neuron, *ε*_*g*_ and *ε*_*s*_ are the strengths of lateral connections in GPe and STN networks, respectively, *W*_*glat*_ and *W*_*slat*_ are weight kernels representing lateral connectivity in GPe and STN networks, respectively, *y*_*D*2_ is the output of D2 MSN, *τ*_*GPe*_ and *τ*_*sTN*_ are the time constants of GPe and STN, respectively, *w*_*sg*_ is the connection strength from STN to GPe, *w*_*gs*_ is the connection strength from GPe to STN, and *λ*_*sTN*_ is the parameter which controls the slope of the sigmoid in STN.

###### Lateral Connections in STN-GPe

The lateral connectivity in STN or GPe network is modeled as Gaussian neighbourhood (Muddapu et al., 2019) which is defined by the weight kernel (*W*_*glat*/*slat*_) as,

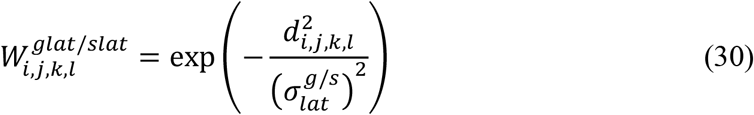

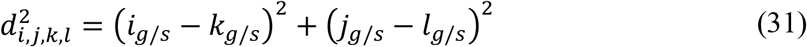

where, 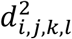is the distance of neuron (*i, j*) from center neuron (*k, l*), 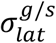is the spread of the lateral connections for GPe or STN network. The detailed analysis of STN-GPe subsystem is described in section S9 of the supplementary information.

###### GPi

The signals arriving from D1 MSN (*y*_*D*1_) and STN (*y*_*sTN*_) via direct and indirect pathways, respectively combines in GPi which is defined as,

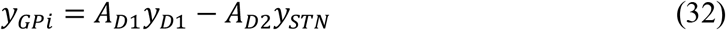

where, *y*_*D*1_ is the output of D1 MSN via direct pathway, *y*_*sTN*_ is the output of STN via indirect pathway, *A*_*D*1_ and *A*_*D*2_ are the gains of direct and indirect pathways, respectively.

#### 2.2.2 Thalamus

The combined inputs (*y*_*GPi*_) at GPi from direct (*y*_*D*1_) and indirect (*y*_*sTN*_) pathways are then passed on to thalamus. To integrate and filter the information from the GPi output, thalamus was modeled as a CANN which is defined as,

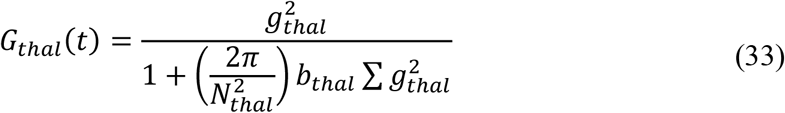

where, *g*_*thal*_ is the internal state of thalamus CANN, *N*_*thal*_ is the size of thalamus network, *b*_*thal*_ is the constant term.

The internal state of the thalamus CANN (*g*_*thal*_) is given by,

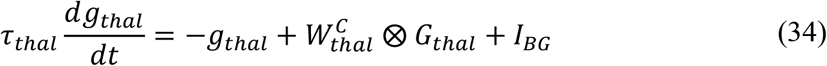

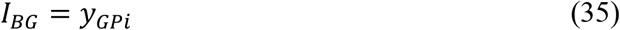

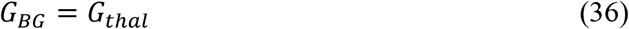

where, 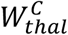 is the weight kernel representing lateral connectivity in thalamus CANN, which determines the local excitation/global inhibition dynamics, *G*_*thal*_ is the output activity of thalamus CANN, *I*_*BG*_ is the total input coming into thalamus CANN from BG, *y*_*GPi*_ is the output of GPi, *G*_*BG*_ is the BG-derived network activity of thalamus, and ⨂ represents the convolution operation.

### 2.3 Central Loop – Nigro-Striatal Loop

The functional pathway of the central loop is as represented in Figure 2C. The central loop consists of the striatum (the input nucleus of BG) and SNc. SNc projects to the striatum via dopaminergic axons (*DA*_*e*_) and striatum in turn projects to SNc via inhibitory GABAergic axons (*I*_*gaba*_). Based on the sensory feedback signal received from the PC (*X*^*arm*^) and the target information from the PFC (*X*^*targ*^), the striatum performs value computation (*V*^*arm*^). Based on the values from current (*V*^*arm*^(*t*)) and previous (*V*^*arm*^(*t* − 1)) instants, the value difference (error, *δ*_*V*_) is computed. Based on the value difference (*δ*_*V*_), appropriate amount of GABAergic current (*I*_*gaba*_) is sent to SNc, which in turn releases dopamine into the striatum (*DA*_*e*_) accordingly.

#### 2.3.1 SNc

##### 2.3.1.1 SNc Neuron (soma)

The detailed single-compartmental biophysical model of the SNc neuron is adopted from (Muddapu and Chakravarthy, 2021). The model incorporates all the essential molecular level mechanisms such as ion channels, active pumps, ion exchangers, dopamine turnover processes etc.

Based on the value difference signal (*δ*_*V*_), the inhibitory GABAergic current (*I*_*gaba*_), flows from striatum to SNc. *I*_*gaba*_along with excitatory glutamatergic current (*I*_*nmda*/*ampa*_) contributes to the overall synaptic input current flux (*J*_*syn*_) to the SNc neurons. *J*_*syn*_ regulates the membrane voltage of the SNc along with the sodium, calcium and potassium fluxes as given by,

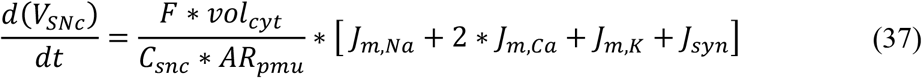

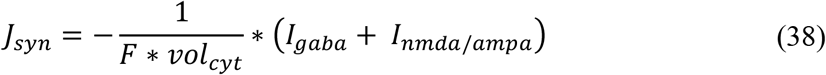

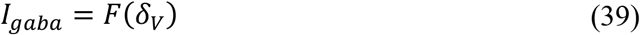

where, *F* is the Faraday’s constant, *C*_*snc*_ is the SNc membrane capacitance, *vol*_*cyt*_ is the cytosolic volume, *AR*_*pmu*_ is the cytosolic area, *J*_*m,Na*_ is the sodium membrane ion flux, *J*_*m,Ca*_ is the calcium membrane ion flux, *J*_*m,K*_ is the potassium membrane ion flux, *J*_*syn*_ is the overall input current flux, *δ*_*V*_ is the value difference, *I*_*gaba*_ is the inhibitory GABAergic current flux, and *I*_*nmda*/*ampa*_ is the excitatory glutamatergic (NMDA/AMPA) current flux.

The membrane voltage of SNc (*V*_*sNc*_) regulates the calcium membrane ionic flux which results in calcium oscillations inside SNc neuron. The calcium membrane ionic flux (*J*_*m,Ca*_) is given by,

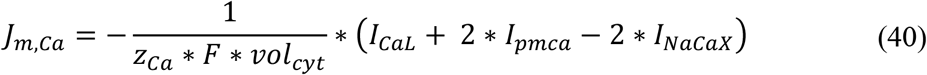

where, *F* is the Faraday’s constant, *z*_*Ca*_ is the valence of calcium ion, *vol*_*cyt*_ is the cytosolic volume, *I*_*CaL*_ is the L-type calcium channel current, *I*_*pmca*_ is the ATP-dependent calcium pump current, and *I*_*NaCaX*_ is the sodium-potassium exchanger current.

The intracellular calcium oscillation or dynamics ([*Ca*_*i*_]) is defined as,

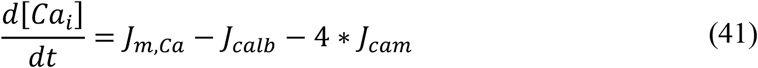

where, *J*_*m,Ca*_ is the flux of calcium ion channels, *J*_*calb*_ is the calcium buffering flux by calbindin, and *J*_*cam*_ is the calcium buffering flux by calmodulin. A detailed description of the SNc neuron is provided in section S4 of the supplementary information.

##### 2.3.1.2 SNc Terminal

The three-compartmental biochemical model of the SNc terminal is adopted from (Muddapu and Chakravarthy, 2021). The SNc terminal model incorporates all the necessary molecular level mechanisms of dopamine turnover process such as synthesis, packing, release and reuptake.

###### Calcium-Dependent Dopamine Release

Dopamine synthesis and release by SNc terminal depends on calcium oscillations. The flux of dopamine release (*J*_*rel*_) from the SNc terminal is given by,

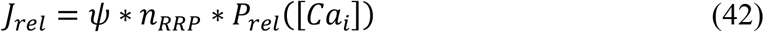

where, [*Ca*_*i*_] is the intracellular calcium concentration in the SNc terminal, *P*_*rel*_ is the release probability as a function of intracellular calcium concentration, *n*_*RRP*_ is the average number of readily releasable vesicles, and *ψ* is the average release flux per vesicle within a single synapse.

###### Calcium-Dependent Dopamine Synthesis

The flux of calcium-dependent dopamine synthesis is defined as,

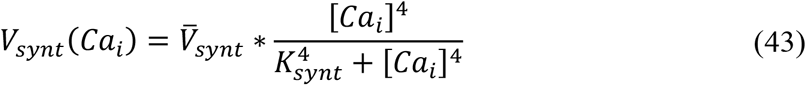

where, *K*_*synt*_ is the calcium sensitivity,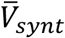 is the maximal velocity for L-DOPA synthesis, and [*Ca*_*i*_] is the intracellular calcium concentration.

The flux of synthesized L-DOPA, *J*_*synt*_, whose velocity is the function of intracellular calcium concentration and L-DOPA synthesis is regulated by the substrate (TYR) itself, extracellular DA (via autoreceptors) and intracellular DA concentrations, is given by,

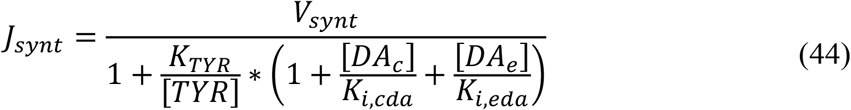

where, *V*_*synt*_ is the velocity of synthesizing L-DOPA, [*TYR*] is the tyrosine concentration in terminal bouton, *K*_*TYR*_ is the tyrosine concentration at which half-maximal velocity was attained, *K*_*i,cda*_ is the inhibition constant on *K*_*TYR*_ due to cytosolic DA concentration, *K*_*i,eda*_ is the inhibition constant on *K*_*TYR*_ due to extracellular DA concentration, [*DA*_*c*_] is the cytoplasmic DA concentration, and [*DA*_*e*_] is the extracellular DA concentration. A detailed description of the SNc terminal is provided in section S5 of the supplementary information.

###### Extracellular Dopamine

The three major mechanisms that determine the dynamics of extracellular DA ([*DA*_*e*_]) in the extracellular compartment (ECS) given by,

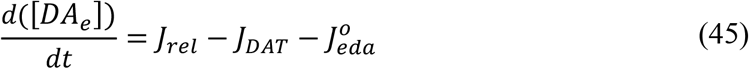

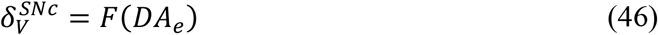

where, *J*_*rel*_ represents the flux of calcium-dependent DA release from the DA terminal, *J*_*DAT*_ represents the unidirectional flux of DA translocated from the ECS into the intracellular compartment (cytosol) via DA plasma membrane transporter (DAT), 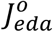 represents the outward flux of DA degradation, which clears DA from ECS, and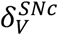 is the SNc-derived value difference. A detailed description of the SNc terminal is provided in section S4 of the supplementary information.

The cortical input to the striatum is modulated by the 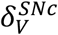 derived from the network of SNc neurons. When 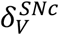is high, the direct pathway will be selected, else the indirect pathway is selected.

### 2.4 Simulating Parkinsonian Conditions

To simulate the Parkinsonian condition in the present model, the number of neurons in SNc population (network) was reduced. In order to kill the SNc neuron, we clamped their membrane voltage (*V*_*sNc*_) to resting membrane voltage (−80 *mV*). As the number of SNc neurons die the total amount of dopamine (*DA*_*e*_) that is made available to the striatum decreases. This influences the selection of the indirect pathway in BG system over the direct pathway resulting in pathological condition. In the present model, two types of PD conditions were simulated: in the first type, SNc cell loss affects striatum alone (PD1) and in the second type, SNc cell loss affects both striatum and STN (PD2).

In normal condition, the SNc-derived value difference 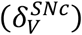 will be similar to actual value difference computed (*δ*_*V*_). In case of PD1, the SNc-derived value difference 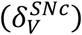 will be lesser than actual value difference computed (*δ*_*V*_). In case of PD2, along with 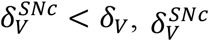 impacts the STN lateral connections, thereby influencing the complexity of STN-GPe subsystem. The STN-GPe subsystem is an integral component of the indirect pathway and is believed to play a major role in exploratory behaviour (Sridharan et al., 2006; Chakravarthy and Balasubramani, 2014).

In normal condition:

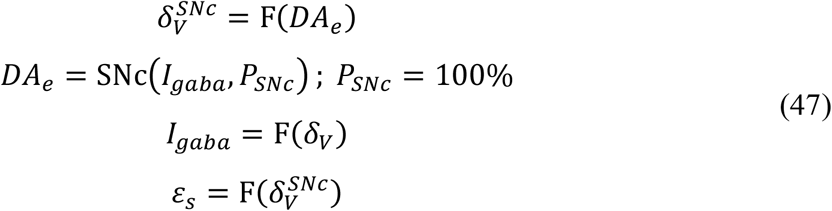

In PD1 condition:

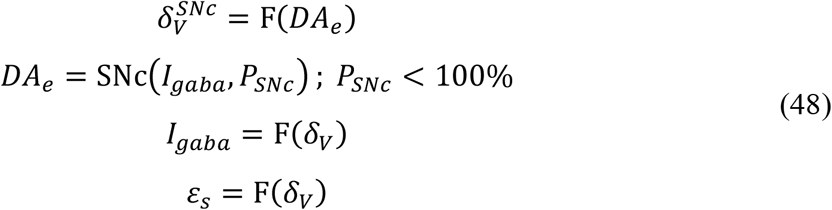

In PD2 condition:

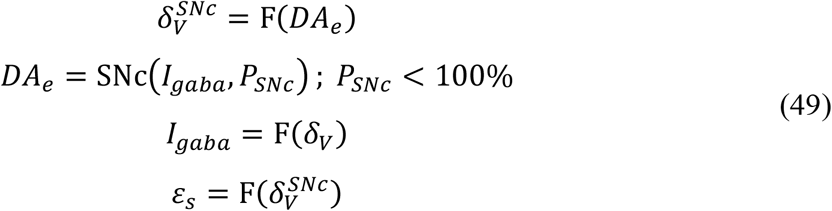

where, 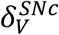 is the SNc-derived value difference, *δ*_*V*_ is the value difference computed, *DA*_*e*_ is the extracellular dopamine, *I*_*gaba*_ is the inhibitory GABAergic current from striatum, *P*_*sNc*_ is the percentage of SNc neurons, and *ε*_*s*_ is the lateral connection strength in STN network.

### 2.5 Levodopa Medication

When a drug is administered to a patient, the medication action is broadly classified into two major branches: pharmacokinetics (what the body does to the drug) and pharmacodynamics (what the drug does to the body) (Shanbhag and Shenoy, 2020).

#### 2.5.1 Pharmacokinetics

Pharmacokinetics deals with absorption, distribution, metabolism and excretion of drugs. In the present study, we have adapted a two-compartment pharmacokinetic model of levodopa (L-DOPA) (Baston et al., 2016), which consists of central and peripheral compartments (Figure 3). Orally consumed L-DOPA is absorbed in the intestine and reaches the blood stream. The blood stream carries the drug all over the body. Proteins break down L-DOPA and around three-fourth of the drug is deactivated before it even reaches the brain. The central compartment where L-DOPA is administered and plasma L-DOPA concentration was measured which is defined as,

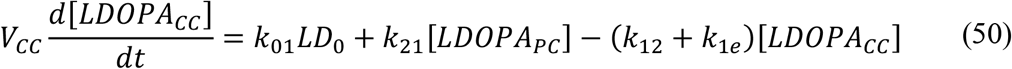

where, *V*_*CC*_ is the volume of central compartment, [*LDOPA*_*CC*_] is the L-DOPA concentration in central compartment, *LD*_0_ is the L-DOPA dose (in milligram), [*LDOPA*_*PC*_] is the L-DOPA concentration in peripheral compartment, *k*_01_ is the infusion rate of *LD*_0_ into central compartment, *k*_21_ is the rate constant from peripheral to central compartments, *k*_12_ is the rate constant from central to peripheral compartments, and *k*_1*e*_ is the total clearance rate constant from central compartment.

**Figure 3:**
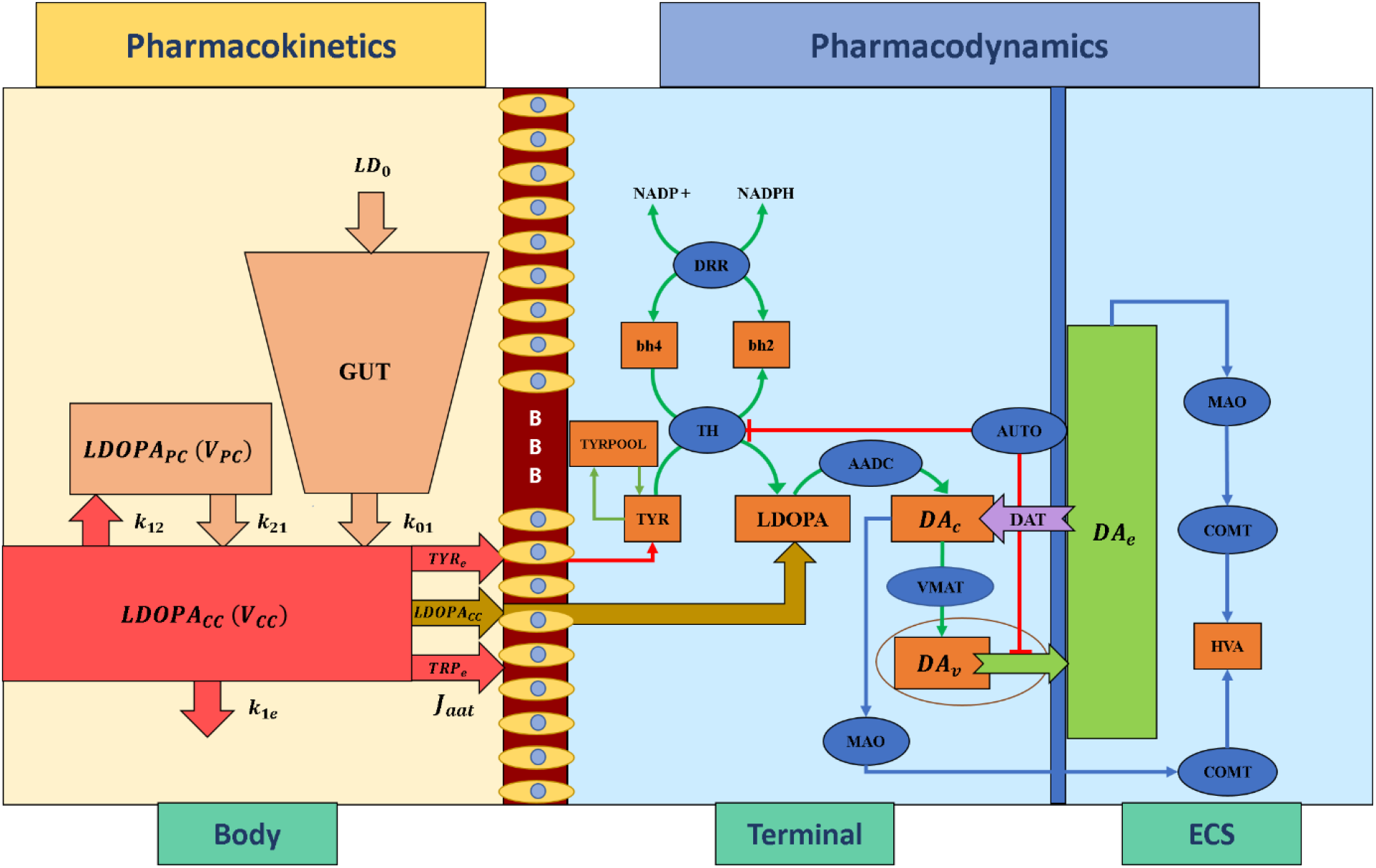
Schematic diagram of pharmacokinetics and pharmacodynamics of levodopa medication. BBB, blood-brain barrier; LDOPA, intracellular levodopa; LDOPA_CC_, levodopa in central compartment; LDOPA_PC_, levodopa in peripheral compartment; V_CC_, volume of central compartment; V_PC_, volume of peripheral compartment; TYR_e_, extracellular tyrosine; TRP_e_, extracellular tryptophan; k_21_, rate constant from peripheral to central compartments, k_12_, rate constant from central to peripheral compartments, k_1e_, total clearance rate constant from central compartment, k_01_, infusion rate of LD_0_ into central compartment, LD_0_, levodopa dose; J_aat_, flux of exogenous L-DOPA transported into the terminal through aromatic L-amino acid transporter; ECS, extracellular space; DA_c_, cytosolic dopamine; DA_v_, vesicular dopamine; DA_e_, extracellular dopamine; TYR, tyrosine; TRYPOOL, tyrosine pool; HVA, homovanillic acid; bh2, dihydrobiopterin; bh4, tetrahydrobiopterin; NADP+, nicotinamide adenine dinucleotide phosphate; NADPH, nicotinamide adenine dinucleotide phosphate hydrogen; TH, tyrosine hydroxylase; DDR, dihydropteridine reductase; AADC, aromatic amino acid decarboxylase; VMAT, vesicular monoamine transporter; DAT: dopamine transporter; AUTO, dopamine autoreceptors; MAO, monoamine oxidase; COMT, catecholamine methyltransferase;

The interaction between plasma L-DOPA and other body fluids, which occurs in the peripheral compartment, is defined as,

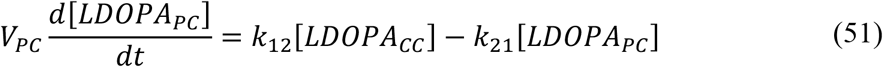

where, *V*_*PC*_ is the volume of peripheral compartment, [*LDOPA*_*CC*_] is the L-DOPA concentration in central compartment, [*LDOPA*_*PC*_] is the L-DOPA concentration in peripheral compartment, *k*_21_ is the rate constant from peripheral to central compartments, and *k*_12_ is the rate constant from central to peripheral compartments.

#### 2.5.2 Pharmacodynamics

Pharmacodynamics deals with molecular, biochemical, and physiological effects of drugs, including drug mechanism of action, receptor binding (including receptor sensitivity), postsynaptic receptor effects, and chemical interactions. In the present study, we have adapted three-compartment dopaminergic terminal model (Reed et al., 2012) which consists of extracellular, vesicular and cytoplasmic compartments.

When L-DOPA medication is administered, the flux of exogenous L-DOPA ([*LDOPA*_*CC*_]) transported into the terminal through aromatic L-amino acid transporter (AAT) while competing with other aromatic amino acids (such as tyrosine (TYR) and tryptophan (TRP)) (Reed et al., 2012) is given by,

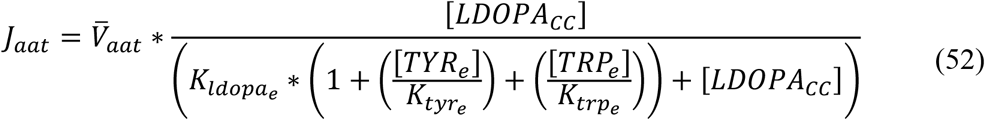

where, 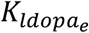 is the extracellular L-DOPA concentration at which half-maximal velocity was attained, 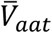 is the maximal velocity with which extracellular L-DOPA was transported into the cytosol, [*LDOPA*_*CC*_] is the extracellular (central compartment) L-DOPA concentration, [*TYR*_*e*_] is the extracellular TYR concentration, [*TRP*_*e*_] is the extracellular TRP concentration, 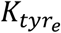 is the affinity constant for [*TYR*_*e*_], 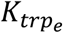 is the affinity constant for [*TRP*_*e*_].

The L-DOPA concentration ([*LDOPA*]) dynamics inside the terminal is given by,

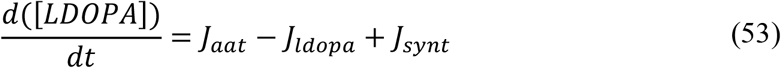

where, *J*_*aat*_ represents the flux of exogenous L-DOPA ([*LDOPA*_*CC*_]) transported into the cytosol, *J*_*ldopa*_ represents the conversion flux of exogenous L-DOPA ([*LDOPA*_*CC*_]) into dopamine, and *J*_*synt*_ represents the flux of synthesized L-DOPA from tyrosine. A detailed description of the dopaminergic terminal is provided in Section S6 of the supplementary information.

### 2.6 Timescales in the Model

Reaching movements, like several other behavioral events, involve dynamics at multiple timescales: the neuronal activity which is generally in milliseconds, and the actual movement which unfolds over the order of seconds. In the present model, the outer (sensory-motor) loop is assumed to run slightly slower than the inner (cortico-basal ganglia) and central (nigro-striatal) loops. As the dynamics of the STN–GPe loop in the indirect pathway needs some time to settle, we run this loop for 2500 iterations (*dt =* 0.02 *ms*), before sending the output to the MC (MC runs for 100 iteration with *dt =* 50 *ms*). Thus, a single update of the MC activity happens after every 50 *ms* during which the BG dynamics run. Similarly, since the dynamics of the SNc neuron needs some time to settle, we run SNc neuron for 2000 iterations (*dt =* 0.025 *ms*), before sending the output to the BG. Thus, a single update of the MC activity happens after every 50 *ms* during which the SNc dynamics run. All the results presented are at the timescale of the MC.

In the present model, the SNc neurons run in milliseconds timescale whereas the pharmacokinetic-pharmacodynamic model of L-DOPA medication runs in hourly timescale. In order to show the drug effect, we sample various points across L-DOPA medication curve (Figure S7.1) and simulated the MCBG model for arm reaching task for each sampled point.

## 3. RESULTS

Here, we showcase the performance of the model starting with training the MCBG model and comparing with previous cortico-basal ganglia model (Figure 4, 5). Next, simulating the PD condition and readout their effects on behavioral outcome (Figure 6). Further, demonstrating the effect of differential dopaminergic axonal loss manifest into some of the cardinal symptoms of PD (Figure 7, 8). Next, assessing the performance in terms of reaching time and verifying the effect of L-DOPA therapeutic intervention (Figure 9, 10). Finally, describing the model results which gave an indicator of how to optimize the drug dosage across the course of the disease progression (Figure 11, 12, 13).

**Figure 4:**
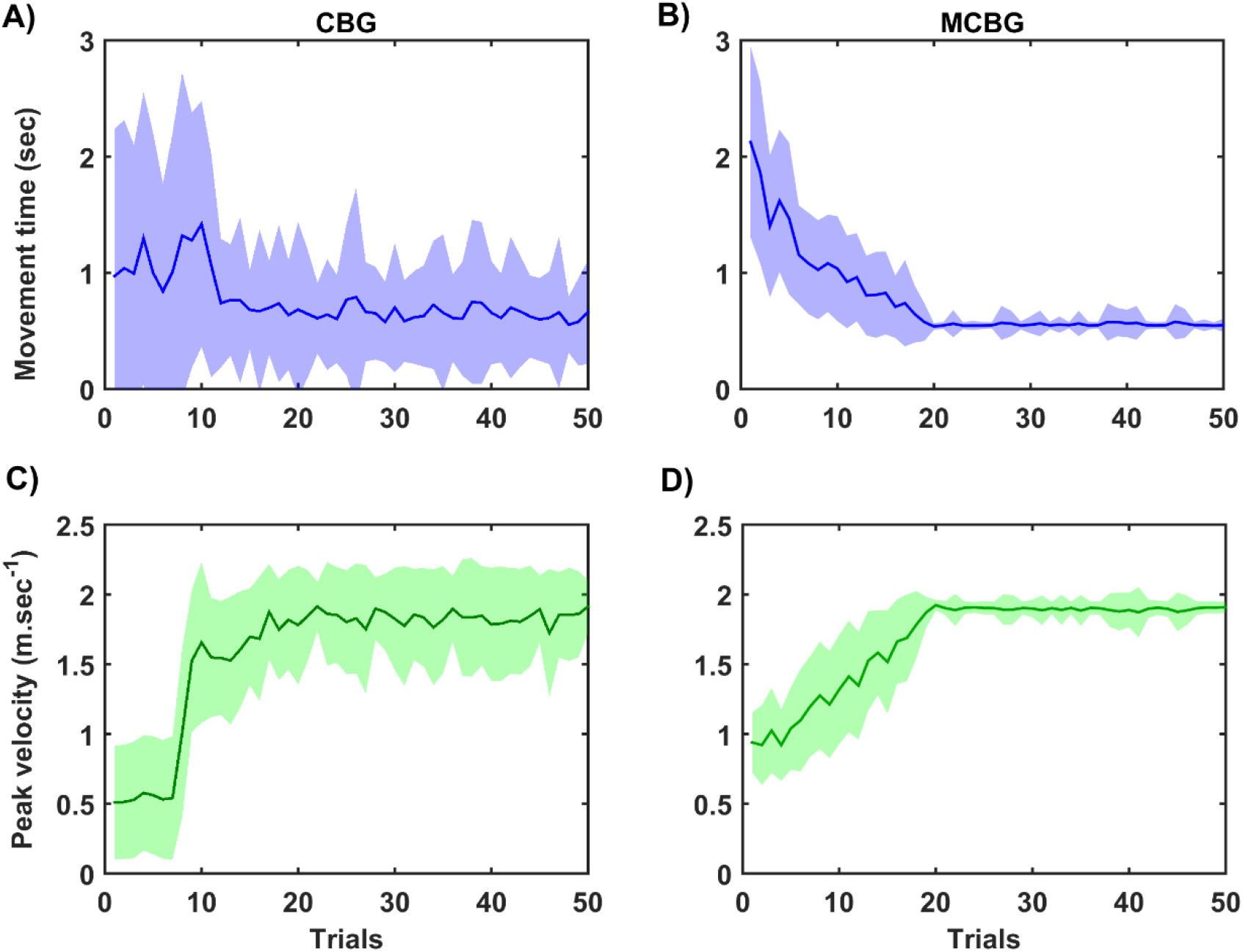
Performance of MCBG model compared with CBG model. A) Movement time and C) Peak velocity in CBG model, B) Movement time and D) Peak velocity in MCBG model. CBG, cortico-basal ganglia model; MCBG, multiscale cortico-basal ganglia model; sec, second; m.sec^-1^, meter per second.

**Figure 5:**
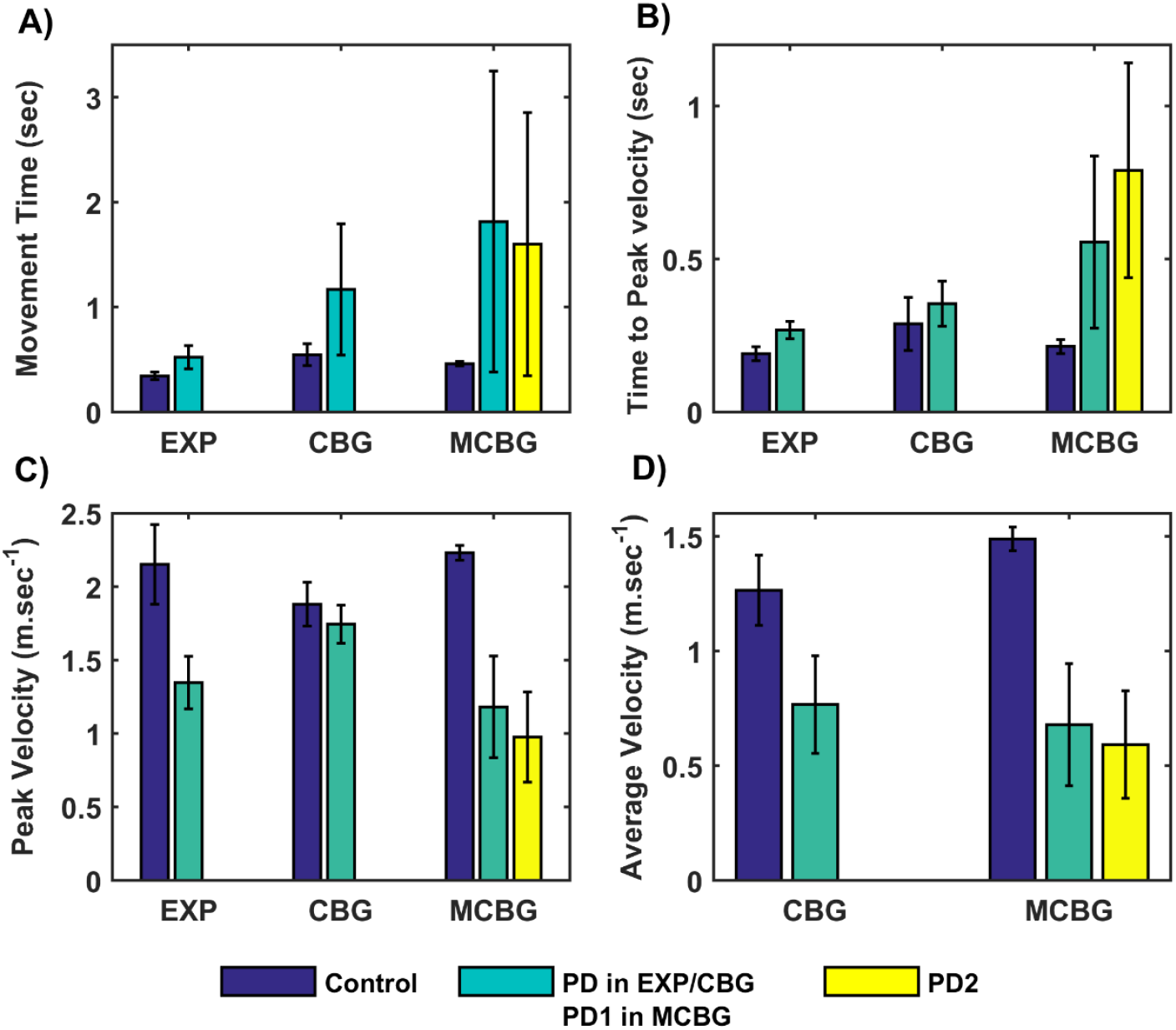
Comparison of performance of the proposed model (during testing phase) with CBG model (Muralidharan et al., 2018) and experimental data adapted from (Majsak et al., 1998). A) Movement time B) Time-to-peak velocity, C) Peak velocity, D) Average velocity. EXP, experiment; CBG, cortico-basal ganglia model; MCBG, multiscale cortico-basal ganglia model; PD1, only striatum affected; PD2, both striatum and subthalamic nucleus affected; sec, second; m/sec, meter per second.

**Figure 6:**
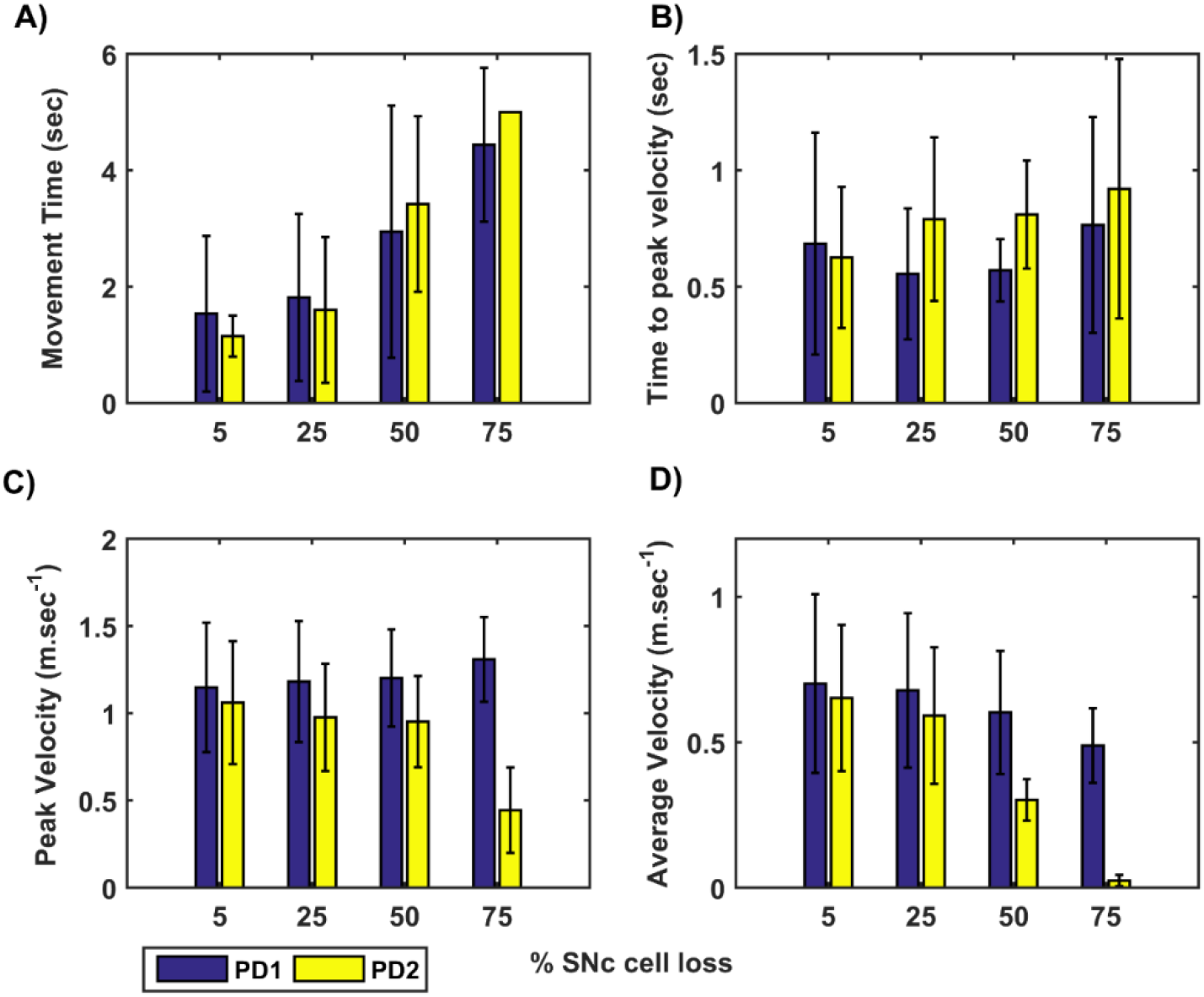
Performance of arm reaching for various PD conditions across different percentage of SNc cell loss. A) Movement time B) Time-to-peak velocity C) Peak velocity D) Average velocity. SNc, substantia nigra pars compacta; PD1, SNc cell loss affecting striatum only; PD2, SNc cell loss affecting both striatum and subthalamic nucleus; sec, second; m.sec^-1^, meter per second.

**Figure 7:**
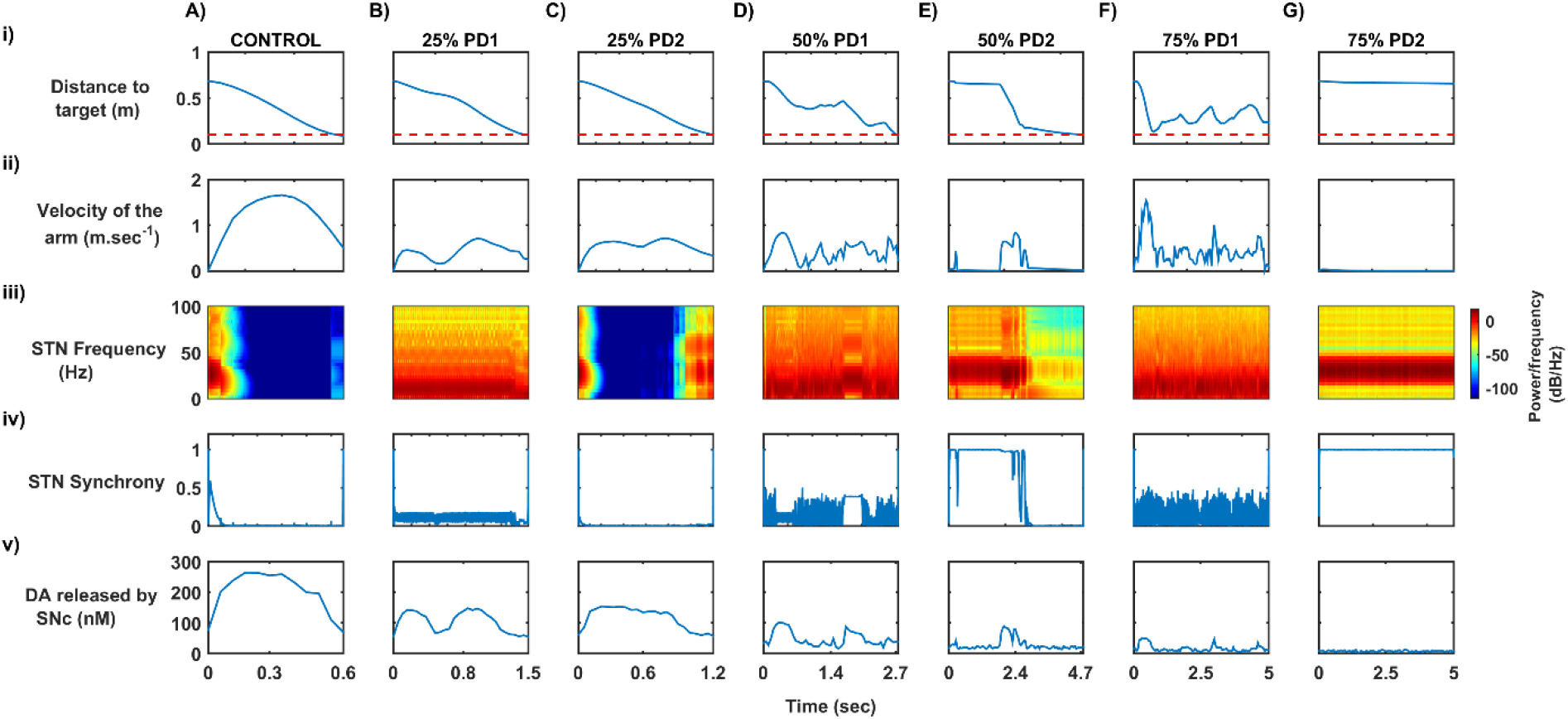
Differential dopaminergic axonal degeneration manifesting in terms of various PD motor symptoms. i) Distance to target ii) Velocity of the arm iii) Spectrogram of STN population iv) Synchrony in STN population v) Dopamine released by SNc extracellularly. SNc, substantia nigra pars compacta; STN, subthalamic nucleus; STR, striatum; DA, dopamine; PD, Parkinson’s disease; sec, second; m/sec, meter per second; Hz, hertz; nM, nanomolar.

**Figure 8:**
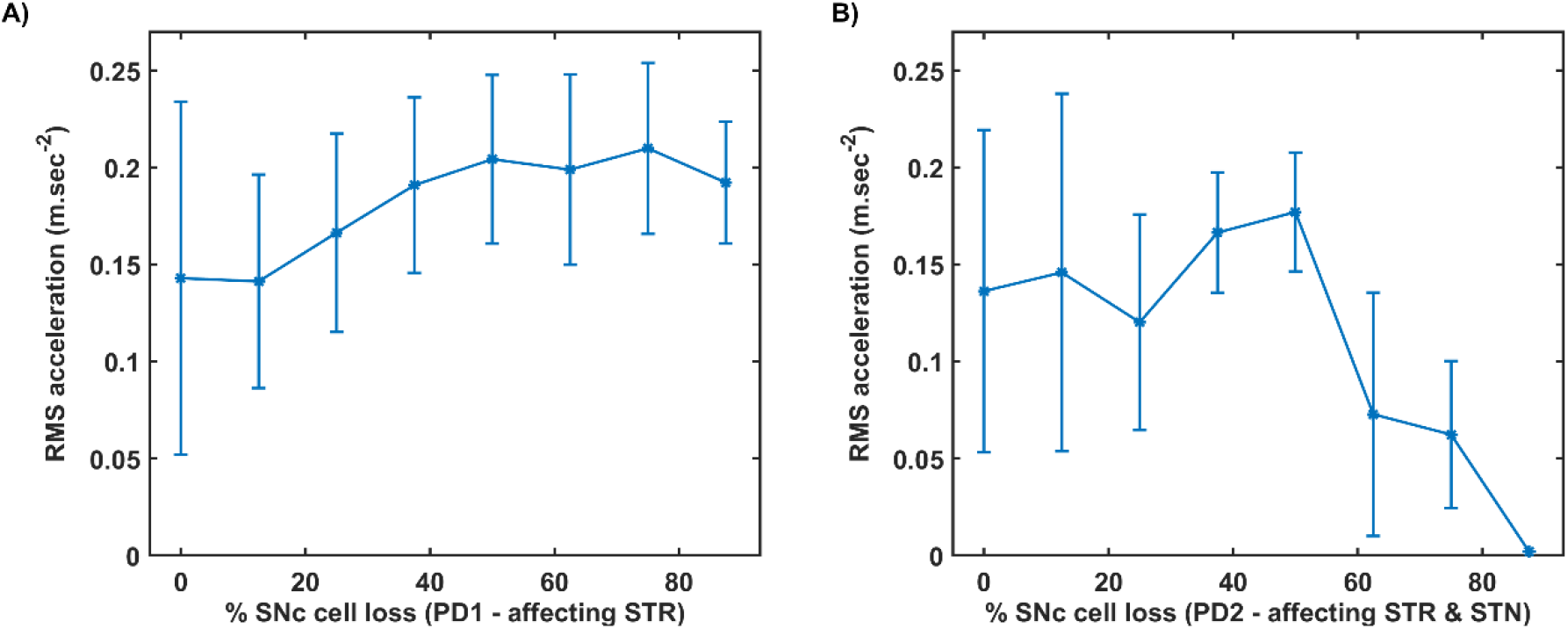
RMS acceleration with respect to percentage loss of SNc cells. A) RMS acceleration when SNc cell loss affecting STR B) RMS acceleration when SNc cell loss affecting STR & STN. SNc, substantia nigra pars compacta; STN, subthalamic nucleus; STR, striatum; PD1, SNc cell loss affecting STR; PD2, SNc cell loss affecting STR & STN; RMS, root mean squared; m/sec^2^, meter per second squared.

**Figure 9:**
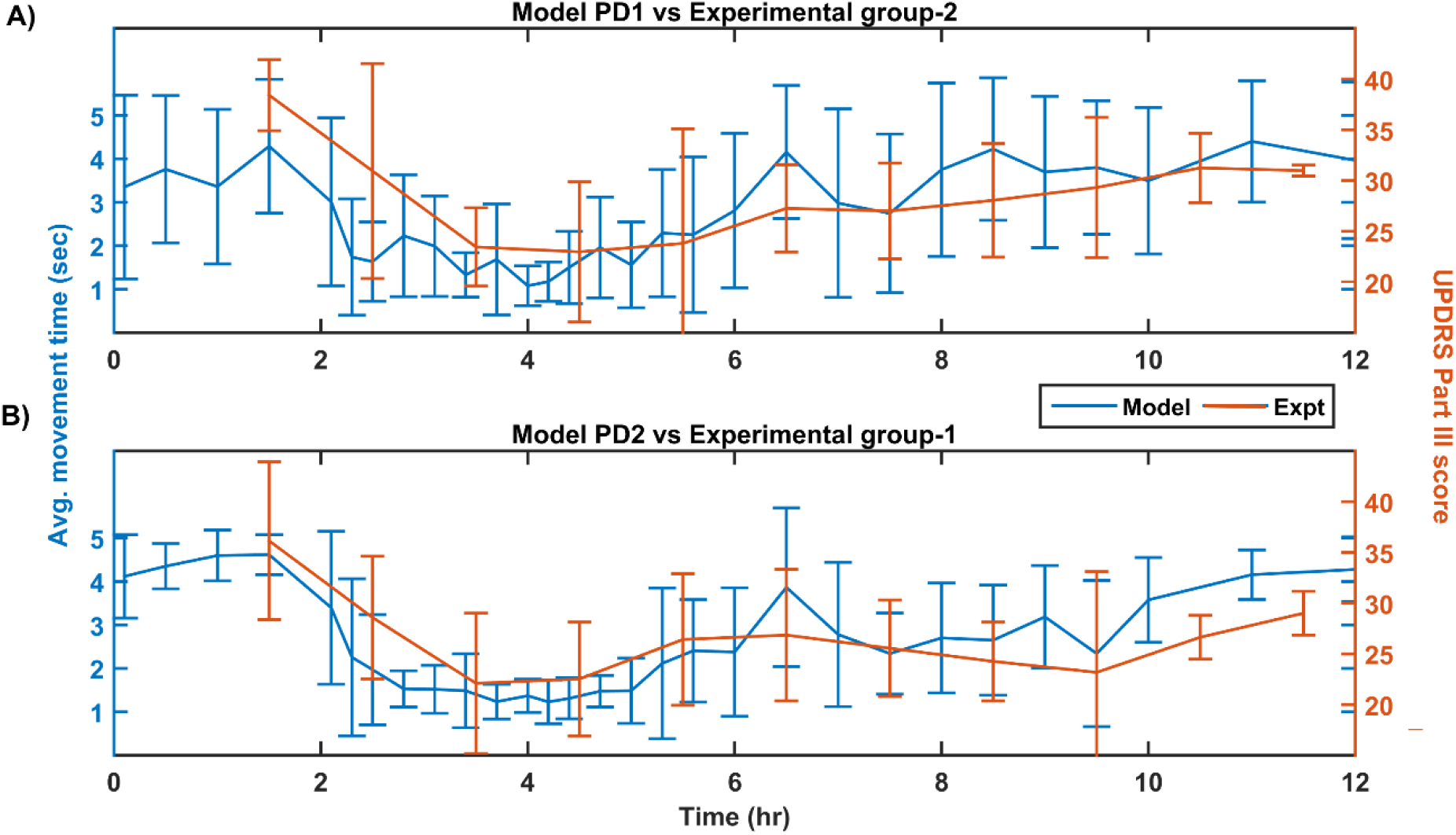
Performance of the model (150 mg L-DOPA and 62% SNc cell loss) compared with experimental study (approx. 140 mg L-DOPA) (Nomoto et al., 2018) for various PD conditions. A) Movement time of PD1 MCBG model was compared with UPDRS Part III score of experimental group-2 after L-DOPA administration B) Movement time of PD2 MCBG model was compared with UPDRS Part III score of experimental group-1 after L-DOPA administration. MCBG, multiscale cortico-basal ganglia model; L-DOPA, levodopa; PD, Parkinson’s disease; PD1, when SNc cell loss affecting STR alone; PD2, when SNc cell loss affecting both STR & STN; SNc, substantia nigra pars compacta; STR, striatum; STN, subthalamic nucleus; UPDRS, unified Parkinson disease rating scale; Expt, experiment; mg, milligram; sec, second; hr, hour.

**Figure 10:**
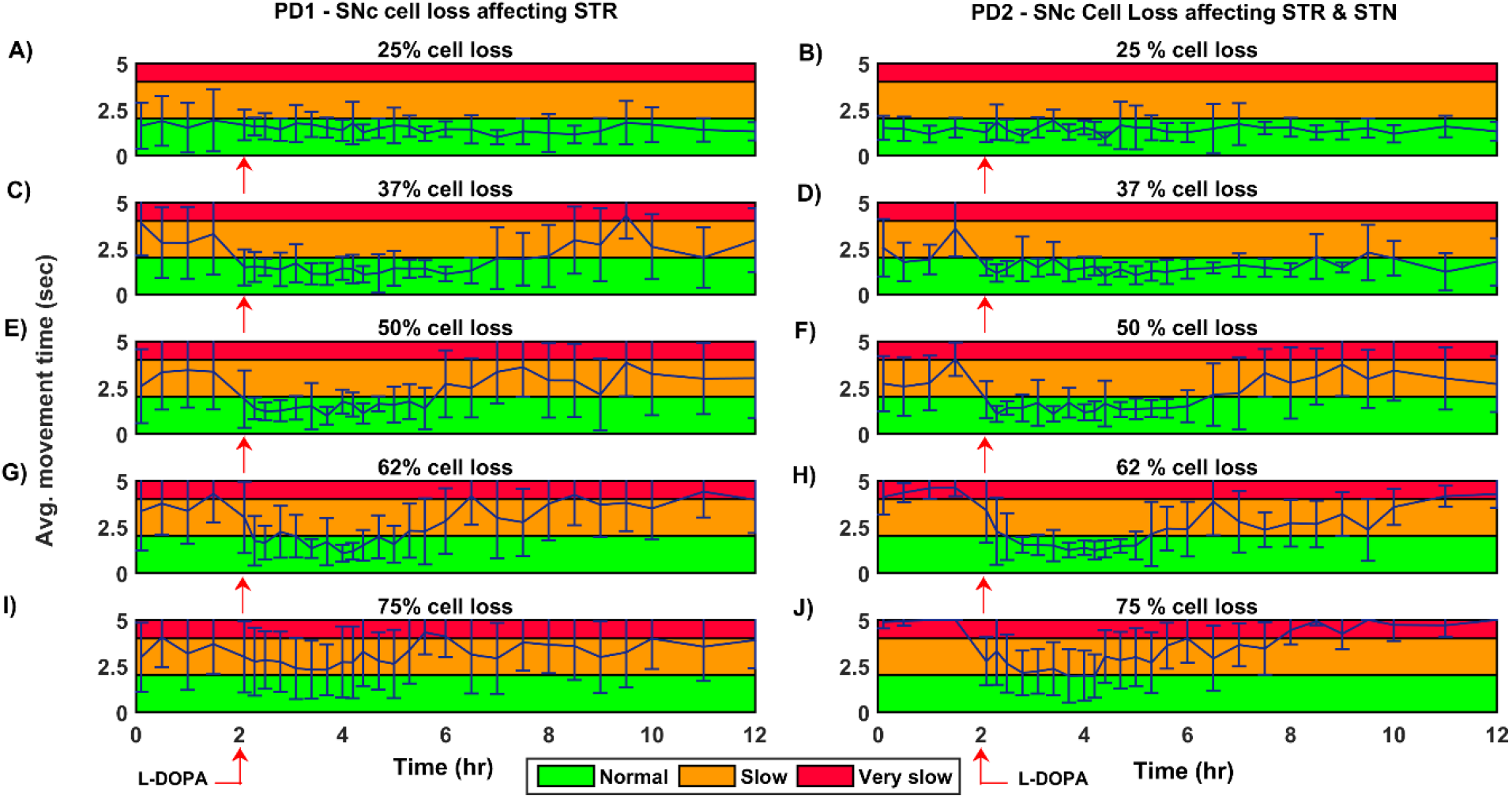
Average time to reach the target for 150 mg L-DOPA medication for various PD conditions. Average movement time for SNc cell loss of 25% (A, B), 37% (C, D), 50% (E, F), 62% (G, H), 75% (I, J) when SNc cell loss affecting STR (PD1) and STR & STN (PD2) during L-DOPA medication (administrated at second hour, indicated by red arrow). The performance of the model during L-DOPA medication is categorized into three regions based on movement time. Green region – when arm reaches the target within 2 seconds; Yellow region – when arm reaches the target between 2 and 4 seconds; Red region – when arm reaches the target beyond 4 seconds. PD1, SNc cell loss affecting STR; PD2, SNc cell loss affecting STR & STN; SNc, substantia nigra pars compacta; STN, subthalamic nucleus; STR, striatum; L-DOPA, levodopa; sec, second; hr, hour.

**Figure 11:**
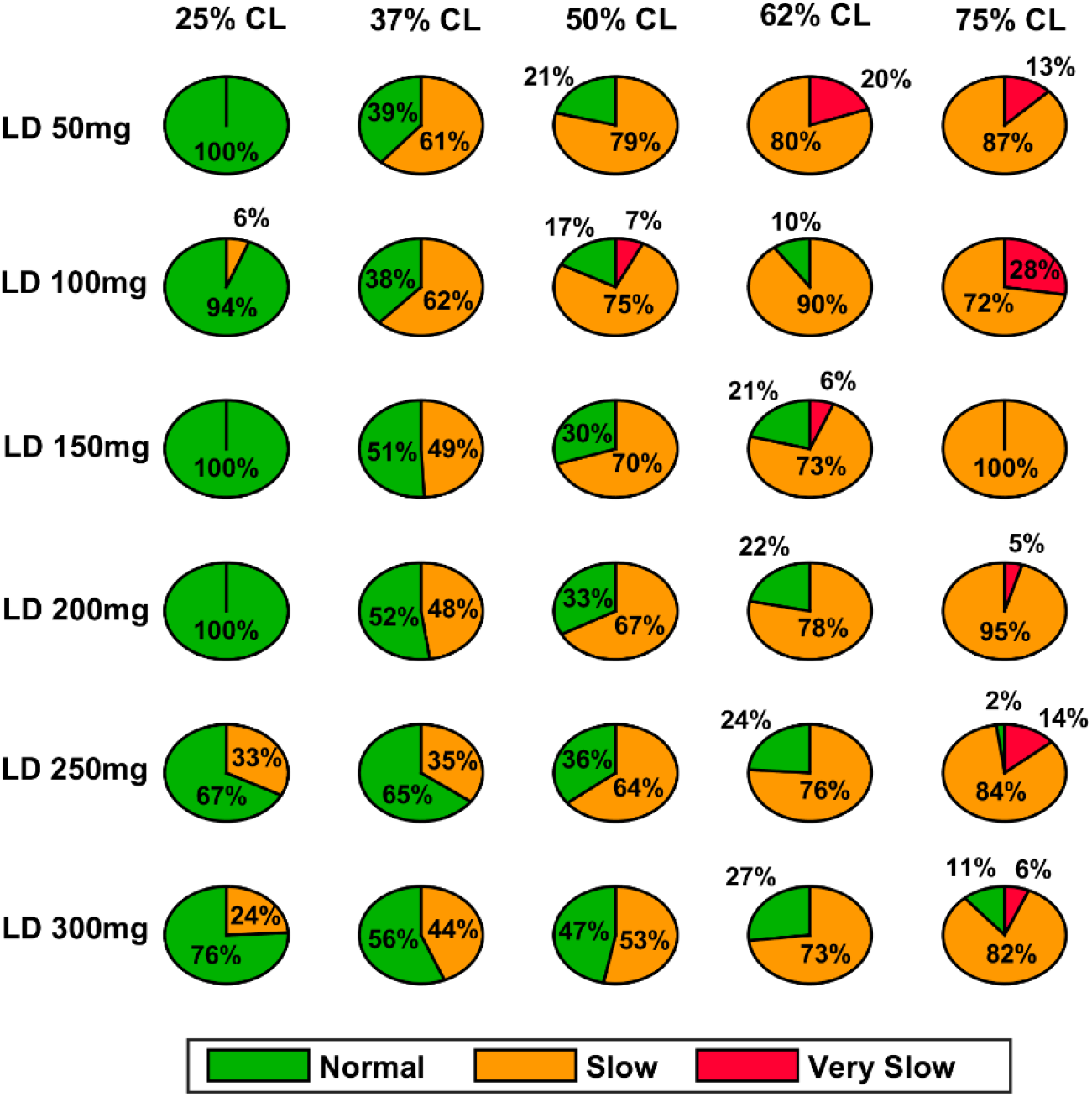
Model performance for different L-DOPA dosage across various percentage SNc cell loss where SNc cell loss affecting STR – PD1. CL, cell loss; LD or L-DOPA, levodopa; SNc, substantia nigra pars compacta; STR, striatum.

**Figure 12:**
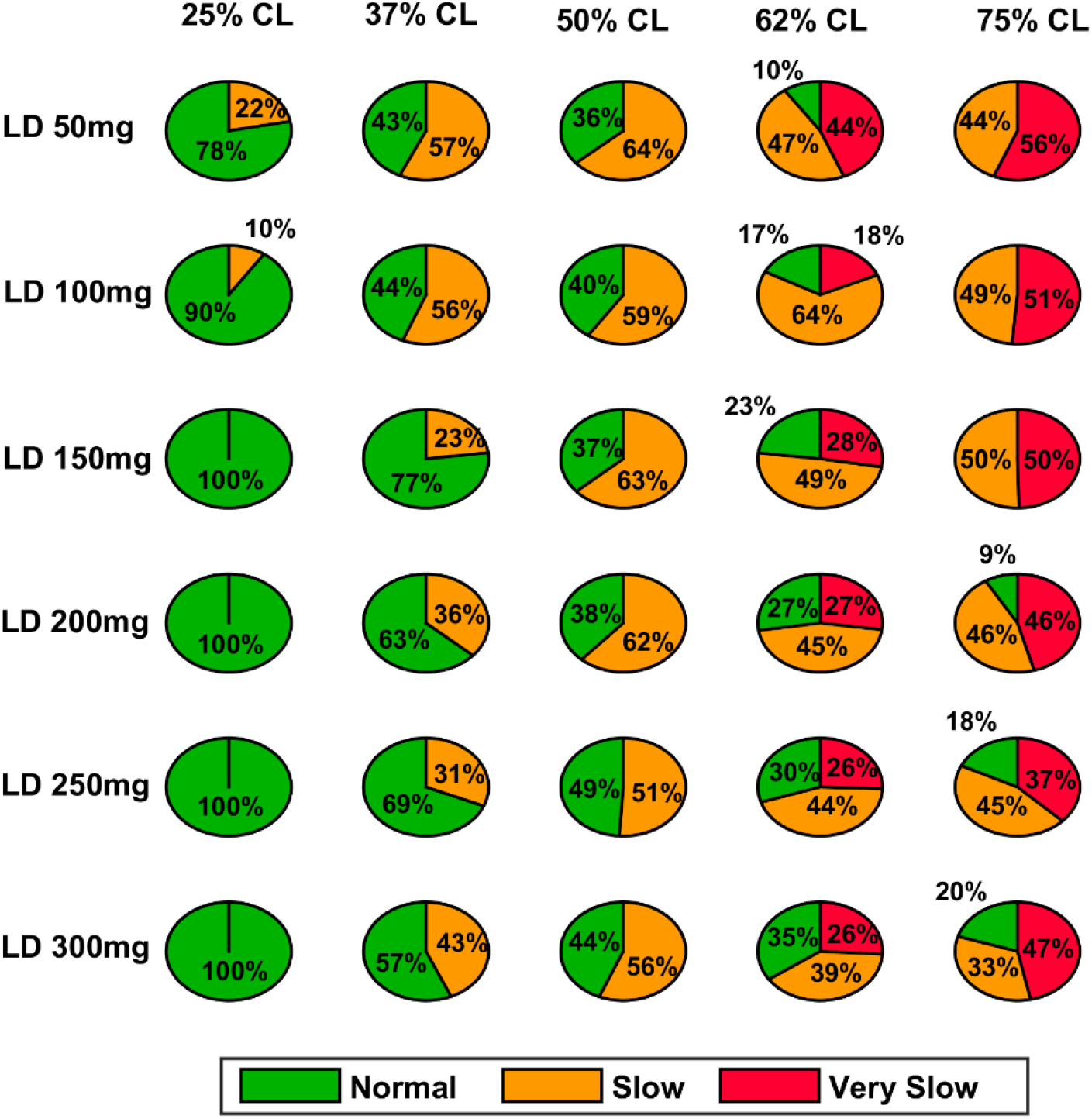
Model performance for different L-DOPA dosage across various percentage SNc cell loss where SNc cell loss affecting STR & STN – PD2. CL, cell loss; LD or L-DOPA, levodopa; SNc, substantia nigra pars compacta; STR, striatum.

**Figure 13:**
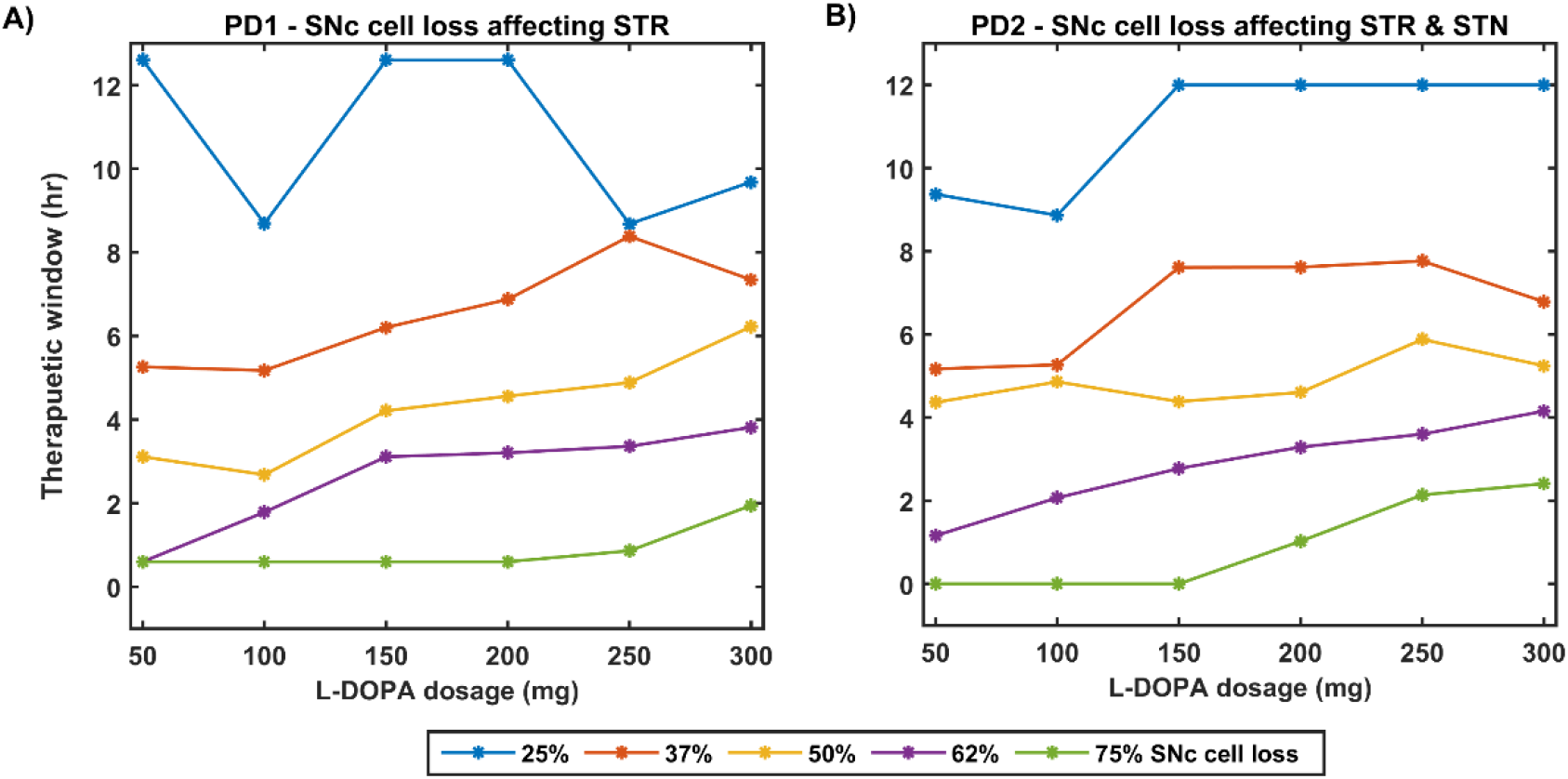
Effect of L-DOPA dosage on therapeutic window for various PD conditions. A) Therapeutic window across different L-DOPA dosage for various percentage of SNc cell loss when SNc cell loss affecting STR B) Therapeutic window across different L-DOPA dosage for various percentage of SNc cell loss when SNc cell loss affecting STR & STN. PD1, SNc cell loss affecting STR; PD2, SNc cell loss affecting STR & STN; SNc, substantia nigra pars compacta; STN, subthalamic nucleus; STR, striatum; L-DOPA, levodopa; mg, milligram; hr, hour.

### 3.1. MCBG Model for Arm Reaching Task

#### 3.1.1. Training Phase

The MCBG model was trained for 50 trials for the arm to reach the target. The performance of MCBG model was compared with the cortico-basal ganglia (CBG) model of (Muralidharan et al., 2018) in arm reaching task during training phase (Figure 4). After 50 trials, MCBG model reaches the target in 0.55 ± 0.05 *sec* compared to CBG model which reaches in 0.67 ± 0.43 *sec* (Figure 4A, 4B). After 50 trials, MCBG model obtained peak velocity of 1.91 ± 0.04 *m. sec*^−1^ compared to CBG model which obtained peak velocity of 1.84 ± 0.34 *m. sec*^−1^ during the arm trajectory towards the target (Figure 4C, 4D). After 50 trials, the performance of MCBG model was better compared to CBG model as the variance in terms of movement time required for the arm to reach the target and peak velocity obtained during the trajectory of the arm moving towards the target was significantly lesser with more number of trials.

#### 3.1.2. Testing Phase

The performance of MCBG model was tested, the model results were compared to that of CBG model (Muralidharan et al., 2018) and the experimental data (Majsak et al., 1998) for both control and PD conditions. In MCBG model, PD conditions simulated were subdivided into two categories: in PD1, the SNc cell loss impacts only striatum whereas in PD2, the SNc cell loss impacts both striatum and STN. The MCBG and the CBG model was tested and the performance was evaluated with respect to movement time, peak velocity, time-to-peak velocity and average velocity along with the experimental results. In control case, MCBG model reaches the target in 0.46 ± 0.02 *sec* compared to CBG model and experimental subject which reaches the target in 0.560 ± 0.10 *sec* and 0.3432 ± 0.04 *sec*, respectively (Figure 5A, dark blue bar). The MCBG model obtained peak velocity of 2.23 ± 0.05 *m. sec*^−1^ compared to CBG model and experimental subject which obtained peak velocity of 1.88 ± 0.15 *m. sec*^−1^ and 2.15 ± 0.27 *m. sec*^−1^, respectively during the arm reaching towards the target in case of control (Figure 5C, dark blue bar). The time taken to reach the peak velocity in case of control was 0.21 ± 0.02 *sec* for MCBG model, 0.29 ± 0.09 *sec* for CBG model and 0.19 ± 0.02 *sec* for experimental subject (Figure 5B, dark blue bar). Finally, the average velocities for MCBG and CBG models were found to be 1.49 ± 0.05 *m. sec*^−1^ and 1.26 ± 0.15 *m. sec*^−1^, respectively in case of control (Figure 5D, dark blue bar).

In case of PD, the experimental subject recorded an average movement time of 0.52 ± 0.63 *sec* respectively (Figure 5A, cyan bar), while CBG model reaches the target in 1.17 ± 0. 3 *sec* (Figure 5A, cyan bar) whereas MCBG model took 1.88 ± 1.42 *sec* and 1.60 ± 1.35 *sec* for PD1 (Figure 5A, cyan bar) and PD2 (Figure 5A, yellow bar), respectively. The experimental subject recorded peak velocity of 1.35 ± 0.18 *m. sec*^−1^ (Figure 5C, cyan bar) compared to CBG model which obtained peak velocity of 1.74 ± 0.13 *m. sec*^−1^ (Figure 5C, cyan bar) whereas MCBG model obtained peak velocities of 1.18 ± 0.35 *m. sec*^−1^ (Figure 5C, cyan bar) and 0.98 ± 0.31 *m. sec*^−1^ (Figure 5C, yellow bar), respectively during the arm trajectory towards the target. The time taken to reach the peak velocity in PD case was 0.27 ± 0.03 *sec* for experimental subject (Figure 5B, cyan bar), 0.35 ± 0.07 *sec* for CBG model (Figure 5B, cyan bar) and 0.56 ± 0.28 *sec*, and 0.79 ± 0.35 *sec* in PD1 (Figure 5B, cyan bar) and PD2 (Figure 5B, yellow bar) cases respectively for MCBG model. Finally, the average velocity for CBG model was found to be 0.77 ± 0.21 *m. sec*^−1^ in PD (Figure 5D, cyan bar), and the average velocities for MCBG model were found to be 0.68 ± 0.27 *m. sec*^−1^ and 0.59 ± 0.23 *m. sec*^−1^ in PD1 (Figure 5D, cyan bar) and PD2 (Figure 5D, yellow bar), respectively.

### 3.2. Simulating Parkinsonian Conditions

To simulate PD conditions in the model, SNc cells were killed and their effects on basal ganglia were considered in two aspects. In the first scenario, only striatum is affected by SNc cell loss (PD1 – cell loss affecting nigrostriatal pathway only) and in second scenario, both striatum and STN are affected by SNc cell loss (PD2 – cell loss affecting both nigrostriatal and nigrosubthalamic pathways).

#### 3.2.1. Effect of SNc Cell Loss on MCBG Behavioral Outcome

To assess the performance metrics with respect to dopaminergic cell loss affecting striatum and both striatum and STN, a comparison study was done with respect to the movement time, peak velocity, time required to peak velocity and average velocity (Figure 6). In both cases (PD1 & PD2), the time required to reach the target (Figure 6A) and time-to-reach the peak velocity (Figure 6B) increases with increase in SNc cell loss. In PD1 case, the peak velocity increases with increase in SNc cell loss when compared to PD2 case where the peak velocity decreases with increase in SNc cell loss (Figure 6C). The reason behind this discrepancy in both cases will be explored in next sections where one leads to tremor-like behavior and other leads to rigidity-like behavior. In both the cases, the average velocity across the trajectory decreases with increase in SNc cell loss (Figure 6D).

#### 3.2.2. Differential Dopaminergic Axonal Degeneration Manifests into Various PD Motor Symptoms

Both the PD scenarios (PD1 & PD2) simulated in the model can be attributed to differential degeneration of dopaminergic projections to various targets in the basal ganglia, and how degeneration manifests into various motor symptoms of PD. In control case, the arm reaches the target in 0.55 *sec* (Figure 7A(i)) with the peak velocity of 1.191 *m. sec*^−1^ (Figure 7A(ii)). The population activity of STN exhibits desynchronous activity during the arm movement which is indicated in the STN spectrogram (Figure 7A(iii)) and synchrony (average value= 0.03) (Figure 7A(iv)) (synchrony measure is described in section S10 of the supplementary information). Dopamine released by SNc neurons in the striatum during the arm reaching peaked at ∼264 *nM* which was in the range of 150 – 400 nM (Schultz, 1998) (Figure 7A(v)).

In 25% PD1, the arm reaches the target in 1.5 *sec* (Figure 7B(i)) with reduced peak velocity of 0.71 *m. sec*^−1^, exhibiting bradykinesia-like behaviour in the arm (Figure 7B(ii)). Population activity of STN exhibits a greater synchrony compared to control case during the arm movement which is also indicated in STN spectrogram (Figure 7B(iii)) and synchrony with average value of 0.11 (Figure 7B(iv)). Dopamine released by SNc neurons in the striatum during the arm reaching peaked at ∼ 148 *nM* which was lesser than in the control case (Figure 7B(v)).

In 25% PD2, the arm reaches the target in 1.2 *sec* (Figure 7C(i)) with the peak velocity of 0.71 *m. sec*^−1^, exhibiting bradykinesia-like behavior in the arm (Figure 7C(ii)). Population activity of STN exhibits desynchronous activity, same as control case during the arm movement which is indicated in the STN spectrogram (Figure 7C(iii)) and synchrony with average value of > 0.01 (Figure 7C(iv)). Dopamine released by SNc neurons in striatum during the arm reaching peaked at ∼ 154 *nM* which was lesser than the control case (Figure 7C(v)).

In 50% PD1, the arm reaches the target in 2.7 *sec* (Figure 7D(i)) with the peak velocity of 0.84 *m. sec*^−1^ showing tremor-like behavior in the arm (Figure 7D(ii)). Population activity of STN exhibits low synchronous activity during the arm movement which indicates in STN spectrogram (Figure 7D(iii)) and synchrony with average value of 0.17 (Figure 7D(iv)). Dopamine released by SNc neurons in the striatum during the arm reaching peaked at ∼ 101 *nM* which was lesser than the control case (Figure 7D(v)). In 50% PD2, the arm reaches the target in 4.7 *sec* (Figure 7E(i)) with the peak velocity of 0.84 *m. sec*^−1^ as a result of cogwheel-like behavior in the arm (Figure 7E(ii)). The population activity of STN exhibits high synchronous activity during the arm movement which indicates in STN spectrogram (Figure 7E(iii)) and synchrony with average value of 0.55 (Figure 7E(iv)). Dopamine released by SNc neurons in striatum during the arm reaching peaked at ∼ 90 *nM* which was lesser than the control case (Figure 7E(v)).

In 75% PD1, the arm did not reach the target within 5 *sec* (Figure 7F(i)) with the peak velocity of 1.54 *m. sec*^−1^ displaying a tremor-like behavior in the arm (Figure 7F(ii)). Population activity of STN exhibits low synchronous activity during the arm movement which indicates in STN spectrogram with increased power in 5 − 25 *Hz* region (Figure 7F(iii)) and synchrony with an average value of 0.15 (Figure 7F(iv)). Dopamine released by SNc neurons in the striatum during the arm reaching peaked at ∼ 51 *nM* which was lesser than the control case (Figure 7F(v)). In 75% PD2, the arm did not reach the target within 5 *sec* (Figure 7G(i)) with zero peak velocity as a result of rigidity-like state of the arm (Figure 7G(ii)). Population activity of STN exhibits high synchronous activity during the arm movement which is indicated in STN spectrogram with increased power in 15 − 50 *Hz* region (Figure 7G(iii)) and synchrony with average value of > 0.99 (Figure 7G(iv)). Dopamine released by SNc neurons in the striatum during the arm reaching peaked at ∼ 13 *nM*, which was lesser than in the control case (Figure 7G(v)).

#### 3.2.3. Quantifying Tremor-Like and Rigidity-Like Motor Symptoms

To quantify between tremor-like and rigidity-like motor symptoms of PD, Root Mean Square (RMS) acceleration was computed across movement trajectory for various PD conditions where RMS acceleration can be used as an indicator of random non-deterministic movements (Figure 8). In PD1 scenario, the RMS acceleration increases with increase in SNc cell loss which indicates irregular changes in velocity of arm movement (Figure 8A). This irregular velocity profile in PD1 is a result of tremor-like motor behavior. In PD2 scenario, the RMS acceleration increases with increase in SNc cell loss till 50% and beyond 50% RMS acceleration decreases with increase in SNc cell loss (Figure 8B). The tremor-like motor behavior is indicated by the RMS acceleration increases until 50% SNc cell loss and from there on, we can see a sudden decrease, which marks the onset of rigidity.

### 3.3. Effect of Levodopa Medication

In order to show L-DOPA medication effect on MCBG model, we simulated different scenarios where various L-DOPA dosages were administrated across various PD conditions and movement time was monitored.

#### 3.3.1. Comparison of MCBG Model with Experimental Results

The L-DOPA therapeutic effect was monitored by recording the performance in terms of the average movement time across the time course of the dosage for the next 10 hours. The performance of the model was also recorded 2 hours prior to the administration of the drug. The MCBG model results were compared with experimental study where PD patients were evaluated based on UPDRS Part III score (Nomoto et al., 2018) (Figure 9). The experimental PD subjects were categorized into two groups based on the UPDRS part III score (motor evaluation) where the group 1 PD subjects have a mean UPDRS III score of 28.0 (13-51) and the group 2 PD have a mean UPDRS III score of 30.3 (22-41) (Nomoto et al., 2018). An average L-DOPA dosage of 141 mg was given to both the experimental groups. The MCBG model was simulated with 62% SNc cell loss and 150 mg of L-DOPA administered at second hour of the simulation.

The PD1 MCBG model performance in terms of movement time (Figure 9A, blue curve) matched with experimental group 2 result in terms of UPDRS III score (Figure 9A, orange curve). Similarly, PD2 MCBG model performance in terms of movement time (Figure 9B, blue curve) matched with experimental group 1 result in terms of UPDRS III score (Figure 9B, orange curve).

#### 3.3.2. Effect of L-DOPA Medication with Disease Progression

The effect of L-DOPA (150 mg) medication on the model performance was studied across different percentages (25%, 37%, 50%, 62% and 75%) of SNc cell loss for both PD1 and PD2 scenarios. The L-DOPA medication was given at the second hour in the simulation. The simulated results show that as SNc cell loss increases, the model performance deteriorates and also the therapeutic effect decreases as the disease progresses in both PD1 and PD2 scenarios (Figure 10). The maximum therapeutic effect of L-DOPA was seen for 50% and 62% SNc cell loss in both PD1 and PD2 scenarios (Figure 10E, 10F). In 75% SNc cell loss, the model performance was poor in case of PD1 when compared to PD2 (Figure 10G, 10H). The model performance was categorized into three regions based on the following criteria: If the arm reaches the target within 2 seconds then that region was marked in green color which indicates the normal movement. If the arm reaches the target between 2 and 4 seconds then that region was marked in yellow color, indicating slow movement or *bradykinesia*. If the arm reaches the target beyond 4 seconds then that region was marked in red color which indicate very slow movement or *akinesia*. The simulated results show that as the SNc cell loss increases the movement time curve shift from green to yellow region when medication was ON and the movement time curve shift from yellow to red region when medication was OFF (Figure 10).

#### 3.3.3. Effect of L-DOPA Dosage and SNc Cell Loss on Therapeutic Window

As discussed in previous section, the model performance was categorized into three regions: green (normal movement), yellow (slow movement, bradykinesia) and red (very slow movement, akinesia). The therapeutic window is computed by taking the time difference between the points when the performance improved after taking medication and entered into the green shaded region until it started wearing off and crosses back to the yellow shaded region (where the effects of L-DOPA start wearing off).

In case of 25%, SNc cell loss (PD1), as the L-DOPA dosage increases the therapeutic window (green region) decreases (Figure 11, first column). But at higher percentage loss of cells (37%, 50%, 62% and 75% SNc cell loss), as the L-DOPA dosage increases the therapeutic window (green region) increased (Figure 11). However, in case of PD2 for all percentages of SNc cell loss, as the L-DOPA dosage increases the therapeutic window (green region) increased (Figure 12).

## 4. DISCUSSION

### 4.1. MCBG Model

The proposed model tries to present a biologically realistic model of the effect of L-DOPA on PD symptoms, specifically in terms of movement parameters. In our modelling approach, a large-scale cortico-basal ganglia model forms the back bone of our network. The two-link arm model that is interfaced to the MNs simulates the movement of the hand and the feedback related to the hand position and distance from the target is processed by the PC and passed on to MC. MC uses the corrective signals from the BG to initiate the next action. The BG dynamics is highly influenced by the dopaminergic input from the SNc and by incorporating a detailed biophysical model of the SNc into the network model, we were able to show the effect of loss of dopaminergic cells on the movement parameters. Going forward we aim to relate the pathological behavior with respect to the dynamics at molecular level happening inside the SNc.

The proposed model was able to explain a wide range of pathological behaviours associated with the PD by controlling the release of dopamine into the extracellular space and reducing the complexity of the STN-GPe network. By reducing the supply of dopamine, the slowness of movement or bradykinesia could be simulated, and in combination with modulating the complexity of STN-GPe network, symptoms like tremor and rigidity were simulated. The complexity of STN-GPe network was varied by controlling the dopaminergic projections of the SNc neurons towards the STN, thereby affecting the lateral connections within the STN subsystem. By progressively reducing the number of dopaminergic cells in SNc, we could replicate some of the cardinal symptoms of PD - bradykinesia, tremor and rigidity.

Once the PD condition and the associated symptoms were simulated, we integrated a pharmacokinetic-pharmacodynamic (PK-PD) model of L-DOPA medication (Baston et al., 2016; Véronneau-Veilleux et al., 2020), which showed improved results in reaching performance. L-DOPA medication is one of the first line treatment methodologies for Parkinson’s disease (Suzuki et al., 2020). Our model incorporates the medication effect by interfacing the SNc with the PK-PD model of L-DOPA drug administration. Depending on the dosage of drug administered, L-DOPA is absorbed into the blood. After interacting with other bodily fluids, a portion of the L-DOPA crosses the BBB and gets absorbed by the dopaminergic terminals. Our results show that consumption of L-DOPA improves the PD symptoms to a great extent. Using our model, we could also see that the extent of improvement on the PD condition depend on the dosage.

A higher level of serum L-DOPA results in dyskinesias and a low-level result in wearing off. Hence, an optimum dosage of medication has to be selected. In order to optimize the drug dosage, we performed our tests with various dosages of L-DOPA medication. We could see that as the percentage of SNc cell loss increases, higher dosage of L-DOPA was required to sustain the medication effect. With increase in percentage of SNc cell loss, the therapeutic effect keeps decreasing. Hence our study focused on the variation of therapeutic effect with respect to the varying percentage SNc cell loss and L-DOPA dosage. The results observed are promising enough to suggest optimal tuning strategies of drug dosage for PD patients (Figure 13). The performance characteristics with respect to the variation in cell loss and the dosage helps us to tune the optimum dosage in terms of the quantity and the frequency of dosage.

From the simulation results, we can explain L-DOPA wearing off mechanism to a great extent. Our hypothesis is that the natural progression of the disease characterized by the increase in loss of SNc cells is one of the mechanisms that contributes to L-DOPA wearing off. There could be other factors as well that can accelerate this wearing off phenomenon. Another hypothesis is that the loss of dopaminergic terminals will lead to synchronized activity in STN which in turn causes overexcitation of SNc neurons resulting in a phenomenon called excitotoxicity in SNc (Muddapu et al., 2019; Muddapu and Chakravarthy, 2020). Thus, fewer dopaminergic terminals and higher L-DOPA dosage results in an accelerated loss of the dopaminergic terminals leading to a faster wearing off. There might be other contributing factors as well that may advance the shortening of the therapeutic window. There is potential scope of carrying out a detailed study on the various causes of the L-DOPA wearing off and we believe our model serves as a good platform to conduct such comprehensive research.

### 4.2. Future Scope

We could reliably replicate some of the cardinal symptoms of PD using our MCBG model. Along with simulating the PD ON/OFF mechanisms, our model could also successfully demonstrate the medication effect of L-DOPA. With the L-DOPA PK-PD model integration with the MCBG model, we could also explain the side effects of L-DOPA medication such as dyskinesias and wearing off. We hypothesize that the natural progression of the disease and the excitotoxicity could be potential factors that result in L-DOPA wearing off. Increase in cytosolic DA will lead to excitotoxicity as unregulated cytosolic DA leads to neurodegeneration (Chen et al., 2008). In this line, the pharmacological model can be extended by incorporating administration of other drugs that blocks the vesicular transporter (Pregeljc et al., 2020). In addition to dopamine-induced excitotoxicity, L-DOPA-induced toxicity can also cause neurodegeneration (Fahn, 2005; Lipski et al., 2011; Witt and Fahn, 2016; Muddapu et al., 2020b). However, there could be other contributing factors too and this model can serve as a starting step to explore research in similar direction. As highlighted in the discussion section, more detailed study of the L-DOPA wearing off mechanism can be carried out to understand the mechanism and devising the alternate or improved medication regimes. Another line of extension is to explore the phenomenon of different types of dyskinesias such as peak dosage and diphasic dyskinesias (Kim et al., 2019b). We also want to extend the model to show the effect of deep brain stimulation (DBS) on motor deficiencies in PD condition and explore the comorbidity effects of both L-DOPA and DBS on PD motor symptoms (Muthuraman et al., 2018; Muddapu et al., 2019; Muddapu and Chakravarthy, 2020; Mueller et al., 2020). One of the limitations of our model is that our model does not consider the influence of hyperdirect pathway, which involves direct cortical connections to the STN (Nambu et al., 2002; Cai et al., 2019). Also, the model does not take into consideration the influence of cholinergic interneurons in the striatum (Crossley et al., 2016; Kim et al., 2019a). These can be considered as further enhancements to the current model. Currently, our model is focusing on the motor deficiencies in the PD pathology. It would be interesting to model PD non-motor symptoms (Goldman and Postuma, 2014; Goldman and Guerra, 2020).

## 5. CONCLUSIONS

A comprehensive test bench for demonstrating the effect of drug action on symptoms can be powerful tool in the therapeutic toolkit of neurodegenerative diseases such as Parkinson’s disease. Our model is a first step towards this bigger goal. In the current study we were able to successfully simulate the relationship between drug dosage, cell loss and PD ON and OFF conditions. We could also demonstrate some of the cardinal symptoms of PD. We also integrated a PK-PD model of L-DOPA medication, which enabled us to simulate the medication effects of the L-DOPA. We also simulated various combinations of L-DOPA medication and percentage of SNc cell loss which enabled us to understand the general trends in drug effects. These modelling results have the potential to optimize the medication in terms of the amount of dosage and the frequency of dosage.

## 6. CODE ACCESSIBILITY

The MATLAB code of the proposed MCBG model (http://modeldb.yale.edu/266907) is available on ModelDB server (McDougal et al., 2017) and access code will be provided on request.

## 7. AUTHOR CONTRIBUTIONS

SSN - Conceptualization; Model development; Data curation; Formal analysis; Investigation; Methodology; Validation; Writing – original draft; VRM - Conceptualization; Model development; Data curation; Formal analysis; Investigation; Methodology; Validation; Visualization; Writing – original draft; VSC - Conceptualization; Model development; Data curation; Formal analysis; Investigation; Methodology; Validation, Writing – review & editing; Supervision. SSN and VRM are equal first authors.

## 8. ACKNOWLEDGEMENTS

We would like to thank Vishant Batta for his contribution in developing the pharmacokinetic-pharmacodynamic model of levodopa medication.

## REFERENCES

Anilkumar, U., Khacho, M., Cuillerier, A., Harris, R., Patten, D. A., Bilen, M., et al. (2020). MCL-1Matrix maintains neuronal survival by enhancing mitochondrial integrity and bioenergetic capacity under stress conditions. Cell Death Dis. 11, 1–12. doi:10.1038/s41419-020-2498-9.

Armstrong, M. J., and Okun, M. S. (2020). Diagnosis and Treatment of Parkinson Disease: A Review. JAMA - J. Am. Med. Assoc. 323, 548–560. doi:10.1001/jama.2019.22360.

Bakshi, S., Chelliah, V., Chen, C., and van der Graaf, P. H. (2019). Mathematical Biology Models of Parkinson’s Disease. CPT Pharmacometrics Syst. Pharmacol. 8, 77–86. doi:10.1002/psp4.12362.

Balestrino, R., and Schapira, A. H. V. (2020). Parkinson disease. Eur. J. Neurol. 27, 27–42. doi:10.1111/ene.14108.

Baston, C., Contin, M., Calandra Buonaura, G., Cortelli, P., and Ursino, M. (2016). A Mathematical Model of Levodopa Medication Effect on Basal Ganglia in Parkinson’s Disease: An Application to the Alternate Finger Tapping Task. Front. Hum. Neurosci. 10. doi:10.3389/fnhum.2016.00280.

Bereczki, D. (2010). The description of all four cardinal signs of Parkinson’s disease in a Hungarian medical text published in 1690. Parkinsonism Relat. Disord. 16, 290–293. doi:10.1016/j.parkreldis.2009.11.006.

Cai, W., Duberg, K., Padmanabhan, A., Rehert, R., Bradley, T., Carrion, V., et al. (2019). Hyperdirect insula-basal-ganglia pathway and adult-like maturity of global brain responses predict inhibitory control in children. Nat. Commun. 10, 1–13. doi:10.1038/s41467-019-12756-8.

Chakravarthy, V. S., and Balasubramani, P. P. (2014). “Basal Ganglia System as an Engine for Exploration,” in Encyclopedia of Computational Neuroscience (Springer New York), 1–15. doi:10.1007/978-1-4614-7320-6_81-1.

Chakravarthy, V. S., and Moustafa, A. A. (2018). Computational Neuroscience Models of the Basal Ganglia. Singapore: Springer Singapore doi:10.1007/978-981-10-8494-2.

Chen, L., Ding, Y., Cagniard, B., Van Laar, A. D., Mortimer, A., Chi, W., et al. (2008). Unregulated cytosolic dopamine causes neurodegeneration associated with oxidative stress in mice. J. Neurosci. 28. doi:10.1523/JNEUROSCI.3602-07.2008.

Crossley, M. J., Horvitz, J. C., Balsam, P. D., and Ashby, F. G. (2016). Expanding the role of striatal cholinergic interneurons and the midbrain dopamine system in appetitive instrumental conditioning. J. Neurophysiol. 115, 240–254. doi:10.1152/jn.00473.2015.

Dorsey, E. R., Sherer, T., Okun, M. S., and Bloemd, B. R. (2018). The emerging evidence of the Parkinson pandemic. J. Parkinsons. Dis. 8. doi:10.3233/JPD-181474.

Fahn, S. (2005). Does levodopa slow or hasten the rate of progression of Parkinson’s disease? J. Neurol. 252, iv37–iv42. doi:10.1007/s00415-005-4008-5.

Fu, H., Hardy, J., and Duff, K. E. (2018). Selective vulnerability in neurodegenerative diseases. Nat. Neurosci. 21, 1350–1358. doi:10.1038/s41593-018-0221-2.

Fullard, M. E., Morley, J. F., and Duda, J. E. (2017). Olfactory Dysfunction as an Early Biomarker in Parkinson’s Disease. Neurosci. Bull. 33. doi:10.1007/s12264-017-0170-x.

Giguère, N., Delignat-Lavaud, B., Herborg, F., Voisin, A., Li, Y., Jacquemet, V., et al. (2019). Increased vulnerability of nigral dopamine neurons after expansion of their axonal arborization size through D2 dopamine receptor conditional knockout. PLoS Genet. 15, e1008352. doi:10.1371/journal.pgen.1008352.

Gillies, A., Willshaw, D., Atherton, J., and Arbuthnott, G. (2002). “Functional Interactions within the Subthalamic Nucleus,” in Brain (Oxford University Press), 359–368. doi:10.1007/978-1-4615-0715-4_36.

Goldman, J. G., and Guerra, C. M. (2020). Treatment of Nonmotor Symptoms Associated with Parkinson Disease. Neurol. Clin. 38, 269–292. doi:10.1016/j.ncl.2019.12.003.

Goldman, J. G., and Postuma, R. (2014). Premotor and nonmotor features of Parkinson’s disease. Curr. Opin. Neurol. 27, 434–441. doi:10.1097/WCO.0000000000000112.

Gonzalez-Rodriguez, P., Zampese, E., and Surmeier, D. J. (2020). “Selective neuronal vulnerability in Parkinson’s disease,” in Progress in Brain Research (Elsevier B.V.), 61–89. doi:10.1016/bs.pbr.2020.02.005.

Izawa, J., Kondo, T., and Ito, K. (2004). Biological arm motion through reinforcement learning. Biol. Cybern. 91, 10–22. doi:10.1007/s00422-004-0485-3.

Jagodnik, K. M., and van den Bogert, A. J. (2010). Optimization and evaluation of a proportional derivative controller for planar arm movement. J. Biomech. 43, 1086–1091. doi:10.1016/j.jbiomech.2009.12.017.

Kawahara, T. (1980). Coupled Van der Pol oscillators ? A model of excitatory and inhibitory neural interactions. Biol. Cybern. 39, 37–43. doi:10.1007/BF00336943.

Kim, T., Capps, R. A., Hamade, K. C., Barnett, W. H., Todorov, D. I., Latash, E. M., et al. (2019a). The Functional Role of Striatal Cholinergic Interneurons in Reinforcement Learning From Computational Perspective. Front. Neural Circuits 13, 10. doi:10.3389/fncir.2019.00010.

Kim, Y. E., Jeon, B., Yun, J. Y., Yang, H. J., and Kim, H. J. (2019b). Chronological View of Peak and Diphasic Dyskinesia, Wearing off and Freezing of Gait in Parkinson’s Disease. J. Parkinsons. Dis. 9, 741–747. doi:10.3233/JPD-191624.

Kohonen, T. (2001). Self-Organizing Maps. 3rd ed. Berlin, Heidelberg: Springer-Verlag Berlin Heidelberg doi:10.1007/978-3-642-56927-2.

Lipski, J., Nistico, R., Berretta, N., Guatteo, E., Bernardi, G., and Mercuri, N. B. (2011). L-DOPA: A scapegoat for accelerated neurodegeneration in Parkinson’s disease? Prog. Neurobiol. 94, 389–407. doi:10.1016/j.pneurobio.2011.06.005.

Magdoom, K. N., Subramanian, D., Chakravarthy, V. S., Ravindran, B., Amari, S., and Meenakshisundaram, N. (2011). Modeling Basal Ganglia for Understanding Parkinsonian Reaching Movements. Neural Comput. 23, 477–516. doi:10.1162/NECO_a_00073.

Majsak, M. J., Kaminski, T., Gentile, A. M., and Flanagan, J. R. (1998). The reaching movements of patients with Parkinson’s disease under self-determined maximal speed and visually cued conditions. Brain 121, 755–766. doi:10.1093/brain/121.4.755.

Marino, B. L. B., de Souza, L. R., Sousa, K. P. A., Ferreira, J. V., Padilha, E. C., da Silva, C. H. T. P., et al. (2020). Parkinson’s Disease: A Review from Pathophysiology to Treatment. Mini-Reviews Med. Chem. 20, 754–767. doi:10.2174/1389557519666191104110908.

Marras, C., Beck, J. C., Bower, J. H., Roberts, E., Ritz, B., Ross, G. W., et al. (2018). Prevalence of Parkinson’s disease across North America. npj Park. Dis. 4. doi:10.1038/s41531-018-0058-0.

McDougal, R. A., Morse, T. M., Carnevale, T., Marenco, L., Wang, R., Migliore, M., et al. (2017). Twenty years of ModelDB and beyond: building essential modeling tools for the future of neuroscience. J. Comput. Neurosci. 42, 1–10. doi:10.1007/s10827-016-0623-7.

Michel, P. P., Hirsch, E. C., and Hunot, S. (2016). Understanding Dopaminergic Cell Death Pathways in Parkinson Disease. Neuron 90, 675–691. doi:10.1016/j.neuron.2016.03.038.

Morley, J. F., and Duda, J. E. (2010). Olfaction as a biomarker in Parkinsons disease.Biomark. Med. 4. doi:10.2217/bmm.10.95.

Muddapu, V. R., and Chakravarthy, V. S. (2020). A Multi-Scale Computational Model of Excitotoxic Loss of Dopaminergic Cells in Parkinson’s Disease. Front. Neuroinform. 14, 34. doi:10.3389/fninf.2020.00034.

Muddapu, V. R., and Chakravarthy, V. S. (2021). Influence of Energy Deficiency on the Subcellular Processes of Substantia Nigra Pars Compacta Cell for Understanding Parkinsonian Neurodegeneration. Sci. Rep. 11, 1754. doi:10.1038/s41598-021-81185-9.

Muddapu, V. R., Dharshini, S. A. P., Chakravarthy, V. S., and Gromiha, M. M. (2020a). Neurodegenerative Diseases – Is Metabolic Deficiency the Root Cause? Front. Neurosci. 14, 213. doi:10.3389/fnins.2020.00213.

Muddapu, V. R., Mandali, A., Chakravarthy, V. S., and Ramaswamy, S. (2019). A Computational Model of Loss of Dopaminergic Cells in Parkinson’s Disease Due to Glutamate-Induced Excitotoxicity. Front. Neural Circuits 13, 11. doi:10.3389/fncir.2019.00011.

Muddapu, V. R., Vijaykumar, K., Ramakrishnan, K., and Chakravarthy, V. S. (2020b). A Computational Model of Levodopa-Induced Toxicity in Substantia Nigra Pars Compacta in Parkinson’s Disease. bioRxiv, 1–55. doi:10.1101/2020.04.05.026807.

Mueller, K., Urgošík, D., Ballarini, T., Holiga, Š., Möller, H. E., Růžička, F., et al. (2020). Differential effects of deep brain stimulation and levodopa on brain activity in Parkinson’s disease. Brain Commun. 2. doi:10.1093/braincomms/fcaa005.

Muralidharan, V., Mandali, A., Balasubramani, P. P., Mehta, H., Srinivasa Chakravarthy, V., and Jahanshahi, M. (2018). “A Cortico-Basal Ganglia Model to Understand the Neural Dynamics of Targeted Reaching in Normal and Parkinson’s Conditions,” in Computational Neuroscience Models of the Basal Ganglia, eds. V. S. Chakravarthy and A. A. Moustafa (Singapore: Springer Singapore), 167–195. doi:10.1007/978-981-10-8494-2_10.

Muthuraman, M., Koirala, N., Ciolac, D., Pintea, B., Glaser, M., Groppa, S., et al. (2018). Deep brain stimulation and L-DOPA therapy: Concepts of action and clinical applications in parkinson’s disease. Front. Neurol. 9, 711. doi:10.3389/fneur.2018.00711.

Nambu, A., Tokuno, H., and Takada, M. (2002). Functional significance of the cortico-subthalamo-pallidal “hyperdirect” pathway. Neurosci. Res. 43, 111–117. doi:10.1016/S0168-0102(02)00027-5.

Narayanamurthy, R., Jayakumar, S., Elango, S., Muralidharan, V., and Chakravarthy, V. S. (2019). A Cortico-Basal Ganglia Model for choosing an optimal rehabilitation strategy in Hemiparetic Stroke. Sci. Rep. 9. doi:10.1038/s41598-019-49670-4.

Nomoto, M., Nagai, M., Nishikawa, N., Ando, R., Kagamiishi, Y., Yano, K., et al. (2018). Pharmacokinetics and safety/efficacy of levodopa pro-drug ONO-2160/carbidopa for Parkinson’s disease. eNeurologicalSci 13, 8–13. doi:10.1016/j.ensci.2018.09.003.

Pacelli, C., Giguère, N., Bourque, M.-J., Lévesque, M., Slack, R. S., and Trudeau, L.-É. (2015). Elevated Mitochondrial Bioenergetics and Axonal Arborization Size Are Key Contributors to the Vulnerability of Dopamine Neurons. Curr. Biol. 25, 2349–2360. doi:10.1016/j.cub.2015.07.050.

Pissadaki, E. K., and Bolam, J. P. (2013). The energy cost of action potential propagation in dopamine neurons: clues to susceptibility in Parkinson’s disease. Front. Comput. Neurosci. 7, 13. doi:10.3389/fncom.2013.00013.

Poewe, W., Seppi, K., Tanner, C. M., Halliday, G. M., Brundin, P., Volkmann, J., et al. (2017). Parkinson disease. Nat. Rev. Dis. Prim. 3, 1–21. doi:10.1038/nrdp.2017.13.

Pregeljc, D., Teodorescu-Perijoc, D., Vianello, R., Umek, N., and Mavri, J. (2020). How Important Is the Use of Cocaine and Amphetamines in the Development of Parkinson Disease? A Computational Study. Neurotox. Res. 37. doi:10.1007/s12640-019-00149-0.

Reed, M. C., Nijhout, H. F., and Best, J. A. (2012). Mathematical Insights into the Effects of Levodopa. Front. Integr. Neurosci. 6, 1–24. doi:10.3389/fnint.2012.00021.

Schultz, W. (1998). Predictive Reward Signal of Dopamine Neurons. J. Neurophysiol. 80, 1– 27. doi:10.1152/jn.1998.80.1.1.

Shanbhag, T., and Shenoy, S. (2020). Pharmacology for Medical Graduates. 4th ed. New Delhi: Elsevier India Available at: https://www.elsevier.com/books/pharmacology-for-medical-graduates-4th-updated-edition/shanbhag/978-81-312-6259-7 [Accessed December 5, 2020].

Sridharan, D., Prashanth, P. S., and Chakravarthy, V. S. (2006). The role of the basal ganglia in exploration in A neural model based on reinforcement learning. Int. J. Neural Syst. 16. doi:10.1142/S0129065706000548.

Surmeier, D. J. (2018). Determinants of dopaminergic neuron loss in Parkinson’s disease. FEBS J. 285, 3657–3668. doi:10.1111/febs.14607.

Suzuki, M., Arai, M., Hayashi, A., and Ogino, M. (2020). Adherence to treatment guideline recommendations for Parkinson’s disease in Japan: A longitudinal analysis of a nationwide medical claims database between 2008 and 2016. PLoS One 15, e0230213. doi:10.1371/journal.pone.0230213.

Trappenberg, T. P. (2011). “Continuous Attractor Neural Networks,” in Recent Developments in Biologically Inspired Computing (IGI Global), 398–425. doi:10.4018/978-1-59140-312-8.ch016.

Véronneau-Veilleux, F., Ursino, M., Robaey, P., Lévesque, D., and Nekka, F. (2020). Nonlinear pharmacodynamics of levodopa through Parkinson’s disease progression. Chaos 30, 093146. doi:10.1063/5.0014800.

Witt, P. A. L., and Fahn, S. (2016). Levodopa therapy for Parkinson disease: A look backward and forward. Neurology 86, S3–S12. doi:10.1212/WNL.0000000000002509.

Zadravec, M., and Matjačić, Z. (2013). Planar arm movement trajectory formation: An optimization based simulation study. Biocybern. Biomed. Eng. 33, 106–117. doi:10.1016/j.bbe.2013.03.006.

