## Supplementary_Information for "A Multiscale, Systems-level, Neuropharmacological Model of Cortico-Basal Ganglia System for Arm Reaching under Normal, Parkinsonian and Levodopa Medication Conditions"

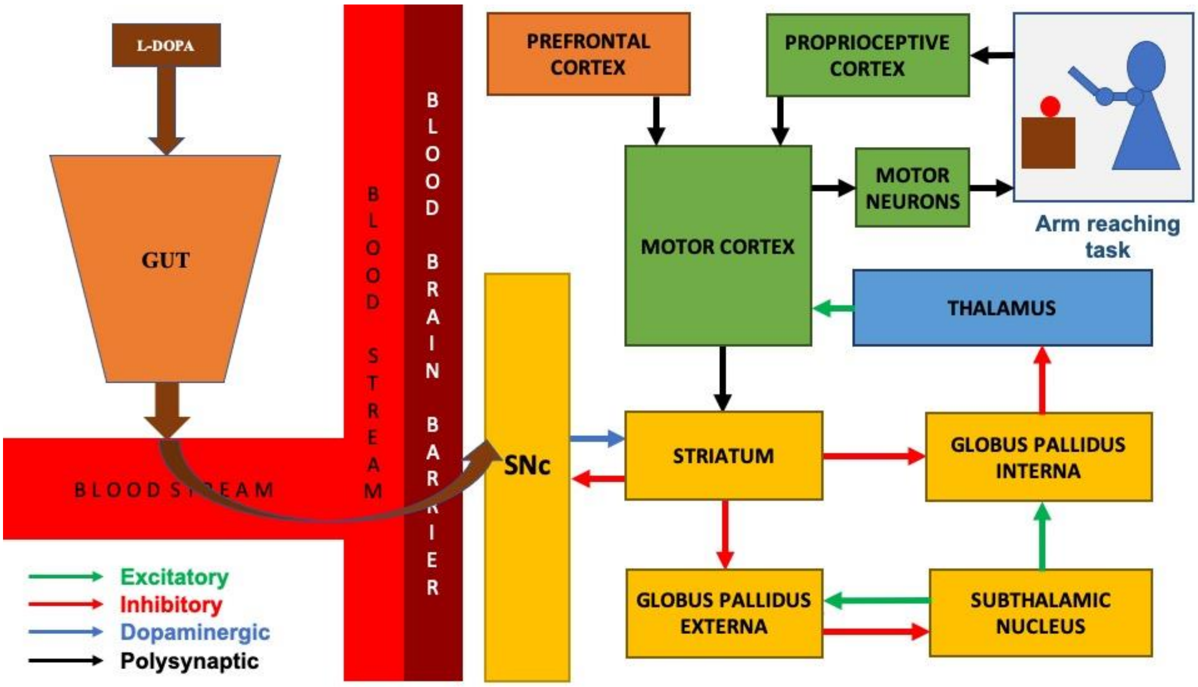


***Figure S1: Model architecture of multiscale cortico-basal ganglia model for arm reaching.*** *SNc, substantia nigra pars compacta; L-DOPA, levodopa.*

### S2. Planar Human Arm Model (Izawa et al., 2004; Zadravec and Matjačić, 2013)

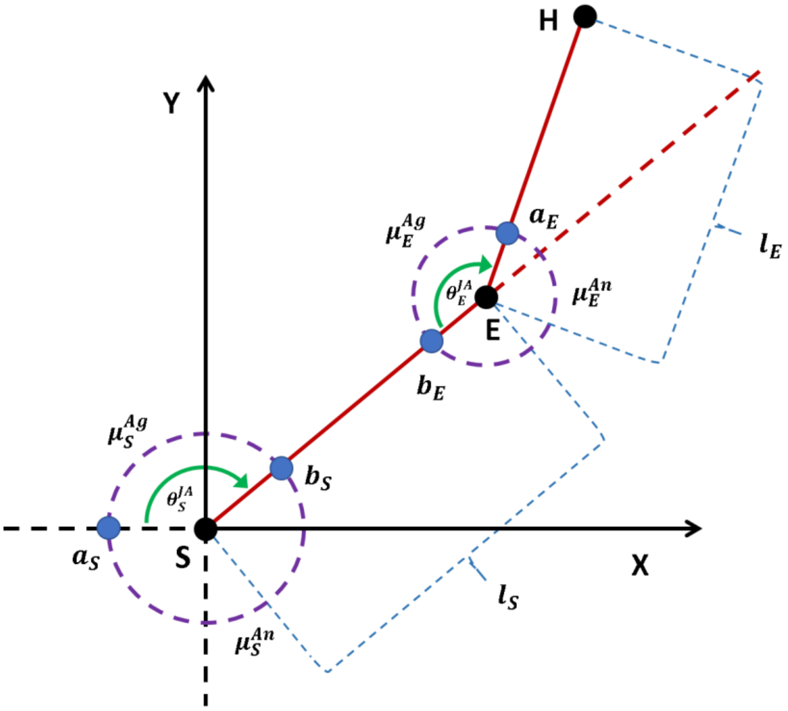


***Figure S2: Planar human arm model.*** *$S$ is the shoulder joint; $E$ is the elbow joint; $H$ is the end effector; $X$ and $Y$ are axes of the two-dimensional plane; $l_S$ is the distance between the shoulder joint center ($S$) and elbow joint center ($E$),; $l_E$ is the distance between the elbow joint center ($E$) and end effector*

$(H)$; $\theta_S^{JA}$ and $\theta_E^{JA}$ are the joint angles for shoulder and elbow, respectively; $\mu_{Ag}^S$ and $\mu_{An}^S$ are the shoulder muscle lengths for agonist (anterior deltoid, $M_1$) and antagonist (posterior deltoid, $M_2$) muscles, respectively; $\mu_{Ag}^E$ and $\mu_{An}^E$ are the elbow muscle lengths for agonist (brachialis, $M_3$) and antagonist (triceps brachii, $M_4$) muscles, respectively; $a_S$ is the distance between shoulder joint center and $M_1$ or $M_2$ moment lever; $b_S$ is the distance between shoulder joint center and $M_1$ or $M_2$ moment lever; $a_E$ is the distance between elbow joint center and $M_3$ or $M_4$ moment lever; $b_E$ is the distance between elbow joint center and $M_3$ or $M_4$ moment lever.

### S3. Training the Outer Loop (Sensory-Motor Loop)

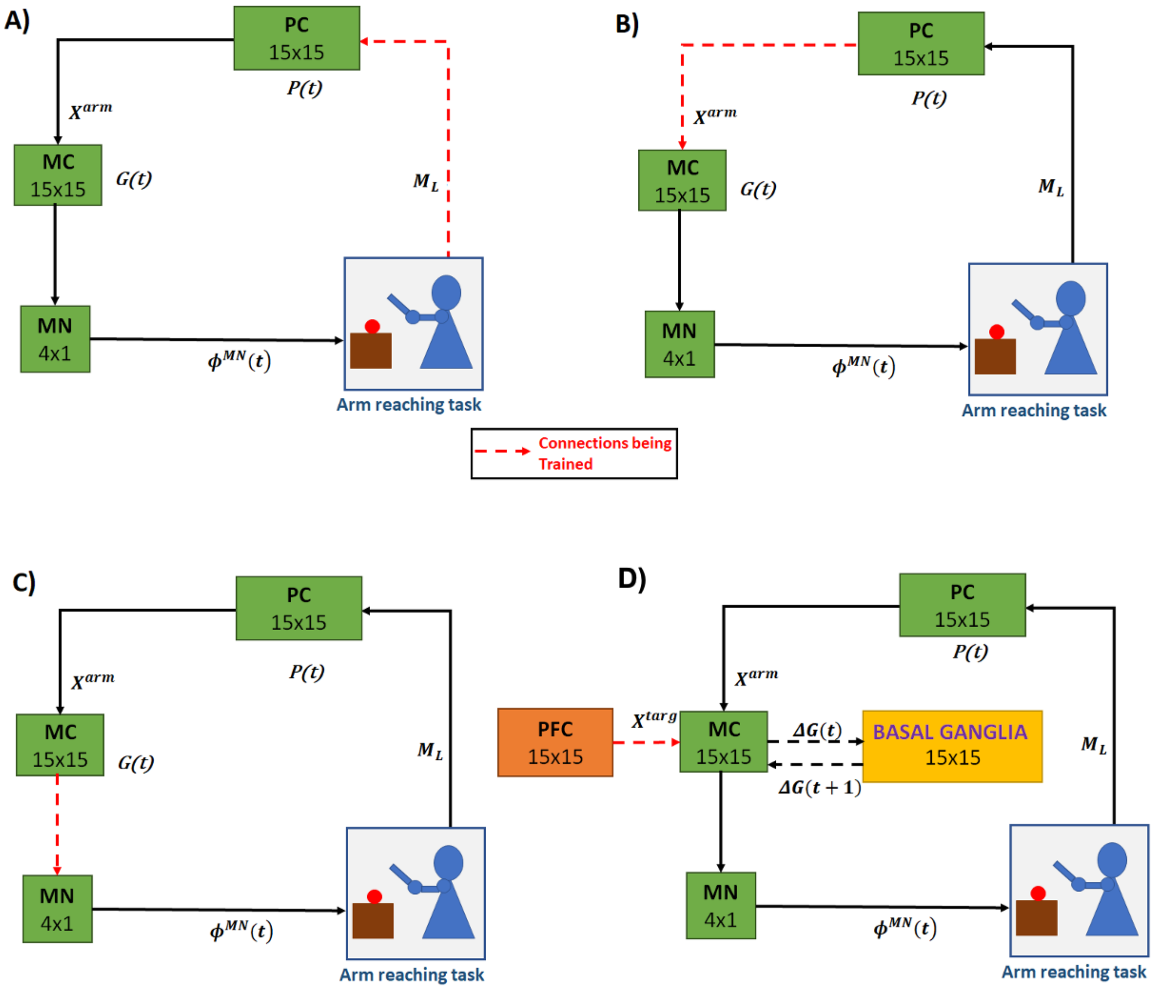


*Figure S3: **The training schema for outer loop (sensory-motor loop).*** *A) Training arm to PC connections B) Training PC to MC connections C) Training MC to MN connections D) BG module is introduced and the PFC to MC connections are trained. In every figure, the dashed arrows indicate the connections that are being trained. PC, proprioceptive cortex; MC, motor cortex; MN, motor neuron; PFC, prefrontal cortex; BG, basal ganglia; $X^{targ}$, the target position; $X^{arm}$, the current arm position; $\phi^{MN}$, the motor neuron activations; $M_L$, muscle lengths; $P(t)$, the PC output; $\Delta G(t)$, the MC output; $\Delta G(t+1)$, the BG-derived activity of thalamus.*

1. First, we generate $n$ random activation vectors for the motor neurons resulting in arm movements. So, in effect, we have $n$ arm configurations. Each such configurations provide the feature vector, Muscle lengths ($M_L$).
2. $M_L$ is provided as the input vector to the PC, which is trained using SOM to form a sensory activation map.
3. The output of PC is presented as an input to MC, and the loop is closed by presenting the output of MC as input to MN in order to obtain the desired activation vector ($\phi^{MN}$), which was initially given to the arm.
4. In the first stage (step 1-3), the sensory map was formed by training the connections between the arm and PC. In the second stage, the connections between PC and MC are trained, and in the third stage, the connections between the MC and MN are trained.
5. After this, BG module is introduced and PFC to MC connections are trained (Figure S3).

**S4. Biophysical Model of SNc**

We have adapted the single-compartmental biophysical model of SNc (Muddapu and Chakravarthy, 2021), where ion-channel dynamics are dependent on ATP levels. The ionic currents which were considered in the soma (Figure S4) are voltage-dependent sodium currents ($I_{Na}$), voltage-dependent potassium currents ($I_K$), voltage-dependent L-type calcium current ($I_{CaL}$), calcium-dependent potassium current $\left(I_{K(Ca)}\right)$, leak current ($I_L$), sodium-potassium ATPase ($I_{NaK}$), calcium ATPase $\left(I_{pmca}\right)$ and sodium-calcium exchanger ($I_{NaCaX}$).

The membrane potential equation for the SNc soma ($V_{SNc}$) is given by,

$$\frac{d(V_{SNc})}{dt} = \frac{F * vol_{cyt}}{C_{snc} * AR_{pmu}} * \left[J_{m,Na} + 2 * J_{m,Ca} + J_{m,K} + J_{inp}\right] \quad (1)$$

where, $F$ is the Faraday's constant, $C_{snc}$ is the SNc membrane capacitance, $vol_{cyt}$ is the cytosolic volume, $AR_{pmu}$ is the cytosolic area, $J_{m,Na}$ is the sodium membrane ion flux, $J_{m,Ca}$ is the calcium membrane ion flux, $J_{m,K}$ is the potassium membrane ion flux, and $J_{inp}$ is the overall input current flux.

*Plasma Membrane Ion Channels*

The intracellular calcium concentration dynamics ($[Ca_i]$) is given by,

$$\frac{d([Ca_i])}{dt} = J_{m,Ca} - J_{calb} - 4 * J_{cam} \tag{2}$$

$$J_{m,Ca} = -\frac{1}{z_{Ca} * F * vol_{cyt}} * \left(I_{CaL} + 2 * I_{pmca} - 2 * I_{NaCaX}\right) \tag{3}$$

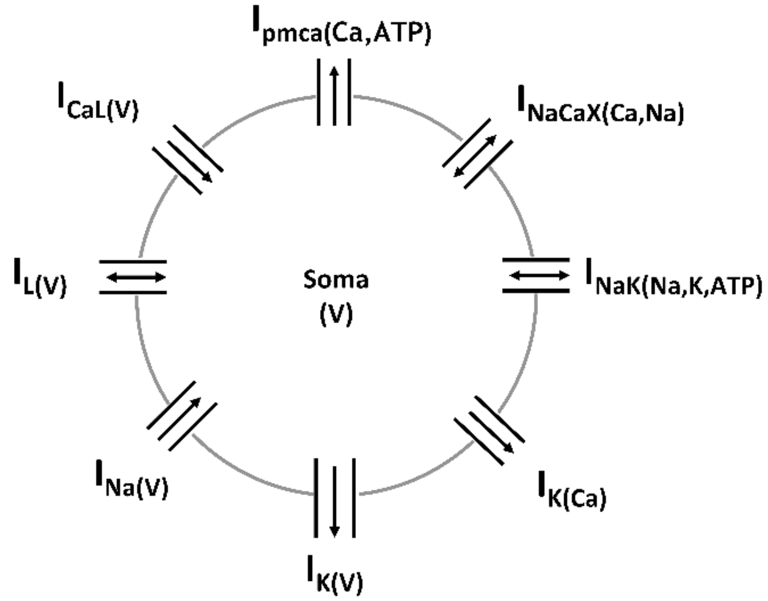


***Figure S4: Schematic of the single-compartment DA neuron model** demonstrating the various ion currents in the proposed model of SNc cell (Muddapu and Chakravarthy, 2021)**.***

$$J_{calb} = k_{1,calb} * [Ca_i] * [Calb] - k_{2,calb} * [CaCalb] \tag{4}$$

$$[CaCalb] = [Calb_{tot}] - [Calb] \tag{5}$$

$$\frac{d([Calb])}{dt} = -J_{calb} \tag{6}$$

$$J_{cam} = \alpha_{cam} * [Cam] - \beta_{cam} * [CaCam] \tag{7}$$

$$[CaCam] = [Cam_{tot}] - [Cam] \tag{8}$$

$$\frac{d([Cam])}{dt} = -J_{cam} \tag{9}$$

$$\alpha_{cam} = K_{cam}^{cb} * K_{cam}^{nb} * \left[\frac{1}{K_{cam}^{cb} + k_{cam}^{nd}} + \frac{1}{k_{cam}^{cd} + k_{cam}^{nd}}\right] \tag{10}$$

$$\beta_{cam} = k_{cam}^{cd} * k_{cam}^{nd} * \left[\frac{1}{K_{cam}^{cb} + k_{cam}^{nd}} + \frac{1}{k_{cam}^{cd} + k_{cam}^{nd}}\right] \tag{11}$$

$$K_{cam}^{cb} = k_{cam}^{cb} * [Ca_i]^2; \qquad K_{cam}^{nb} = k_{cam}^{nb} * [Ca_i]^2 \tag{12}$$

where, $z_{Ca}$ is the valence of calcium ion, $I_{CaL}$ is the L-type calcium channel current, $I_{pmca}$ is the ATP-dependent calcium pump current, $I_{NaCaX}$ is the sodium-potassium exchanger current, $F$ is the Faraday's constant, $vol_{cyt}$ is the cytosolic volume, $J_{m,Ca}$ is the flux of calcium ion channels, $J_{calb}$ is the calcium buffering flux by calbindin, $J_{cam}$ is the calcium buffering flux by calmodulin, $(k_{1,calb}, k_{2,calb})$ are the calbindin reaction rates, $[Ca_i]$ is the intracellular calcium concentration, $[Calb]$ is the calbindin concentration, $[CaCalb]$ is the calcium-bound calbindin concentration, $[Calb_{tot}]$ is the total cytosolic calbindin concentration, $(k_{cam}^{nd}, k_{cam}^{cd}, k_{cam}^{cb}, k_{cam}^{cb})$ are the calmodulin reaction rates, $[Cam]$ is the calmodulin concentration, $[CaCam]$ is the calcium-bound calmodulin concentration, and $[Cam_{tot}]$ is the total cytosolic calmodulin concentration.

The voltage-dependent L-type calcium channel current $(I_{CaL})$ is given by,

$$I_{CaL}(V) = \left(\bar{g}_{Ca,L} * O_{Ca,L}\right) * \left(\sqrt{[Ca_i] * [Ca_e]}\right) * \left(\frac{\sinh(V_D - V_{Ca})}{\left(\frac{\sinh(V_D)}{V_D}\right)}\right) \tag{13}$$

$$O_{Ca,L} = m_{Ca,L} * h_{Ca,L} \tag{14}$$

where, $\bar{g}_{Ca,L}$ is the maximal conductance for calcium channel, $O_{Ca,L}$ is the gating variable of calcium channel, $m_{Ca,L}$ is the activation gate of the L-type calcium channel, $h_{Ca,L}$ is the inactivation gate of L-type calcium channel, $[Ca_i]$ is the intracellular calcium concentration,

$[Ca_e]$ is the extracellular calcium concentration, $V_{Ca}$ is the reversal potential for calcium ion, and $V_D$ is the voltage defined thermodynamic entity.

$$\frac{d(m_{Ca,L})}{dt} = \frac{\frac{1}{\left(1+e^{\left(-\frac{(V+15)}{7}\right)}\right)} - m_{Ca,L}}{7.68 * e^{\left(-\left[\frac{V+65}{17.33}\right]^2\right)} + 0.723} \tag{15}$$

$$h_{Ca,L} = \frac{0.00045}{0.00045 + [Ca_i]} \tag{16}$$

The intracellular sodium concentration ($[Na_i]$) dynamics is given by,

$$\frac{d([Na_i])}{dt} = J_{m,Na} \tag{17}$$

$$J_{m,Na} = -\frac{1}{z_{Na} * F * vol_{cyt}} * (I_{NaT} + 3 * I_{NaK} + 3 * I_{NaCaX}) \tag{18}$$

where, $z_{Na}$ is the valence of sodium ion, $I_{NaT}$ is the total sodium channel current, $I_{NaK}$ is the ATP-dependent sodium-potassium pump current, $I_{NaCaX}$ is the sodium-potassium exchanger current, $F$ is the Faraday's constant, and $vol_{cyt}$ is the cytosolic volume.

The total sodium channel current is given by,

$$I_{NaT} = I_{Na} + I_{NaHCN} + I_{Nalk} \tag{19}$$

where, $I_{Na}$ is the voltage-dependent sodium channel current, $I_{NaHCN}$ is the hyperpolarization-activated cyclic nucleotide-gated sodium channel current, and $I_{Nalk}$ is the leaky sodium channel current.

The voltage-dependent sodium channel current $(I_{Na})$ is given by,

$$I_{Na}(V) = (\bar{g}_{Na} * O_{Na}) * \left(\sqrt{[Na_i] * [Na_e]}\right) * \left(\frac{\sinh\left(\frac{1}{2} * (V_D - V_{Na})\right)}{\left(\frac{\sinh\left(\frac{1}{2} * V_D\right)}{\left(\frac{1}{2} * V_D\right)}\right)}\right) \tag{20}$$

$$O_{Na} = m_{Na}^3 * h_{Na} \tag{21}$$

where, $\bar{g}_{Na}$ is the maximal conductance for sodium channel, $O_{Na}$ is the gating variable of sodium channel, $m_{Na}$ is the activation gate of the sodium channel, $h_{Na}$ is the inactivation gate of the sodium channel, $[Na_i]$ is the intracellular sodium concentration, $[Na_e]$ is the extracellular sodium concentration, $V_{Na}$ is the reversal potential for sodium ion, and $V_D$ is the voltage-defined thermodynamic entity.

$$\frac{d(m_{Na})}{dt} = 1.965 * e^{(1.7127*V_D)} * (1 - m_{Na}) - 0.0424 * e^{(-1.5581*V_D)} * (m_{Na}) \tag{22}$$

$$\frac{d(h_{Na})}{dt} = 0.00009566 * e^{(-2.4317*V_D)} * (1 - h_{Na}) - 0.5296 * e^{(1.1868*V_D)} * (h_{Na}) \tag{23}$$

The hyperpolarization-activated cyclic nucleotide (HCN) gated sodium channel current ($I_{NaHCN}$) is given by,

$$I_{NaHCN}(V) = (\bar{g}_{NaHCN} * O_{NaHCN}) * \left(\sqrt{[Na_i] * [Na_e]}\right) * \left(\frac{\sinh\left(\frac{1}{2} * (V_D - V_{Na})\right)}{\left(\frac{\sinh\left(\frac{1}{2} * V_D\right)}{\left(\frac{1}{2} * V_D\right)}\right)}\right) \tag{24}$$

where, $\bar{g}_{NaHCN}$ is the maximal conductance for sodium HCN channel, $O_{NaHCN}$ is the gating variable of sodium HCN channel, $[Na_i]$ is the intracellular sodium concentration, $[Na_e]$ is the extracellular sodium concentration, $V_{Na}$ is the reversal potential for sodium ion, $V_D$ is the voltage defined thermodynamic entity, and $[cAMP]$ is the cyclic adenosine monophosphate concentration.

$$\frac{d(O_{NaHCN})}{dt} = k_{f,HCN} * (1 - O_{NaHCN}) - k_{r,HCN} * O_{NaHCN} \tag{25}$$

$$k_{f,HCN} \; = \; k_{f,free} * P_c + k_{f,bnd} * (1 - P_c) \tag{26}$$

$$k_{r,HCN} \; = \; k_{r,free} * P_o + k_{r,bnd} * (1 - P_o) \tag{27}$$

$$P_c = \frac{1}{\left(1 + \frac{[cAMP]}{0.001163}\right)}; \qquad P_o = \frac{1}{\left(1 + \frac{[cAMP]}{0.0000145}\right)} \tag{28}$$

$$k_{f,free} = \frac{0.006}{1 + e^{\left(\frac{V+87.7}{6.45}\right)}}; \qquad k_{f,bnd} = \frac{0.0268}{1 + e^{\left(\frac{V+94.2}{13.3}\right)}} \tag{29}$$

$$k_{r,free} = \frac{0.08}{1 + e^{\left(-\frac{V+51.7}{7}\right)}}; \qquad k_{r,bnd} = \frac{0.08}{1 + e^{\left(-\frac{V+35.5}{7}\right)}} \tag{30}$$

The leaky sodium channel current ($I_{Nalk}$) is given by,

$$I_{Nalk}(V) = \; (\bar{g}_{Nalk}) * \left(\sqrt{[Na_i] * [Na_e]}\right) * \left( \frac{\sinh\left(\frac{1}{2} * (V_D - V_{Na})\right)}{\left( \frac{\sinh\left(\frac{1}{2} * V_D\right)}{\left(\frac{1}{2} * V_D\right)} \right)} \right) \tag{31}$$

where, $\bar{g}_{Nalk}$ is the maximal conductance for leaky sodium channel, $[Na_i]$ is the intracellular sodium concentration, $[Na_e]$ is the extracellular sodium concentration, $V_{Na}$ is the reversal potential for sodium ion, and $V_D$ is the voltage defined thermodynamic entity.

The intracellular potassium concentration dynamics ($[K_i]$) is given by,

$$\frac{d([K_i])}{dt} = J_{m,K} \tag{32}$$

$$J_{m,K} = -\frac{1}{z_K * F * vol_{cyt}} * (I_{KT} - 2 * I_{NaK}) \tag{33}$$

where, $z_K$ is the valence of potassium ion, $I_{KT}$ is the total potassium channel current, $I_{NaK}$ is the ATP-dependent sodium-potassium pump current, $F$ is the Faraday's constant, and $vol_{cyt}$ is the cytosolic volume.

The total potassium channel current is given by,

$$I_{KT} = I_{Kdr} + I_{Kir} + I_{Ksk} \tag{34}$$

where, $I_{Kdr}$ is the voltage-dependent (delayed rectifying, DR) potassium channel current, $I_{Kir}$ is the voltage-dependent (inward rectifying, IR) potassium channel current, and $I_{Ksk}$ is the calcium-dependent (small conductance, SK) potassium channel current.

The voltage-dependent (delayed rectifying) potassium channel current $(I_{Kdr})$ is given by,

$$I_{Kdr}(V) = (\bar{g}_{Kdr} * O_{Kdr}) * (V - V_K * V_\tau) \tag{35}$$

$$O_{Kdr} = m_{Kdr}^3 \tag{36}$$

where, $\bar{g}_{Kdr}$ is the maximal conductance for delayed rectifying potassium channel, $O_{Kdr}$ is the gating variable of voltage-dependent (delayed rectifying) potassium channel, $V_K$ is the reversal potential for potassium ion, and $V_\tau$ is the temperature defined thermodynamic entity.

$$\frac{d(m_{K,dr})}{dt} = \frac{\frac{1}{\left(1 + e^{\left(-\frac{(V+25)}{12}\right)}\right)} - m_{K,dr}}{\frac{18}{\left(1 + e^{\left(-\left[\frac{V+65}{17.33}\right]^2\right)}\right)} + 1} \tag{37}$$

The voltage-dependent (inward rectifying) potassium channel current $(I_{Kir})$ is given by,

$$I_{Kir}(V) = (\bar{g}_{Kir} * O_{Kir}) * (V - V_K * V_\tau) \tag{38}$$

$$O_{Kir} = \frac{1}{\left(1 + e^{\left(\frac{V+85}{12}\right)}\right)} \tag{39}$$

where, $\bar{g}_{Kir}$ is the maximal conductance for inward rectifying potassium channel, $O_{Kir}$ is the gating variable of voltage-dependent (inward rectifying) potassium channel, $V_K$ is the reversal potential for potassium ion, and $V_\tau$ is the temperature defined thermodynamic entity.

The calcium-dependent (small conductance) potassium channel current $(I_{Ksk})$ is given by,

$$I_{Ksk}(V) = (\bar{g}_{Ksk} * O_{Ksk}) * \left(\sqrt{[K_i] * [K_e]}\right) * \left(\frac{\sinh\left(\frac{1}{2} * (V_D - V_K)\right)}{\left(\frac{\sinh\left(\frac{1}{2} * V_D\right)}{\left(\frac{1}{2} * V_D\right)}\right)}\right) \tag{40}$$

$$O_{Ksk} = \frac{[Ca_i]^{4.2}}{[Ca_i]^{4.2} + 0.00035^{4.2}} \tag{41}$$

where, $\bar{g}_{Ksk}$ is the maximal conductance for small conductance potassium channel, $O_{Ksk}$ is the gating variable of calcium-dependent (small conductance) potassium channel, $[K_i]$ is the intracellular potassium concentration, $[K_e]$ is the extracellular potassium concentration, $[Ca_i]$ is the intracellular calcium concentration, $V_K$ is the reversal potential for potassium ion, and $V_D$ is the voltage defined thermodynamic entity.

The overall synaptic input current flux $(J_{syn})$ to SNc neuron is given by,

$$J_{syn} = -\frac{1}{F * vol_{cyt}} * \left(I_{syn}^{+} + I_{syn}^{-} - I_{ext}\right) \tag{42}$$

where, $I_{syn}^{+}$ is the excitatory synaptic current, $I_{syn}^{-}$ is the inhibitory synaptic current, $I_{ext}$ is the external current applied, $F$ is the Faraday's constant, and $vol_{cyt}$ is the cytosolic volume.

*Plasma Membrane ATPases*

The plasma membrane sodium-potassium ATPase $(I_{NaK})$ is given by,

$$I_{NaK} = K_{nak} * \left[k_{1,nak} * \mathcal{P}\left(E^{*}_{1,nak}\right) * y_{nak} - k_{2,nak} * \mathcal{P}\left(E^{*}_{2,nak}\right) * (1 - y_{nak})\right] \quad (43)$$

$$\frac{d(y_{nak})}{dt} = \beta_{nak} * (1 - y_{nak}) - \alpha_{nak} * y_{nak} \quad (44)$$

$$\beta_{nak} = k_{2,nak} * \mathcal{P}\left(E^{*}_{2,nak}\right) + k_{4,nak} * \mathcal{P}\left(E^{\#}_{2,nak}\right) \quad (45)$$

$$\alpha_{nak} = k_{1,nak} * \mathcal{P}\left(E^{*}_{1,nak}\right) + k_{3,nak} * \mathcal{P}\left(E^{\#}_{1,nak}\right) \quad (46)$$

$$\mathcal{P}\left(E^{*}_{1,nak}\right) = \frac{1}{\left[1 + \frac{K_{nak,nai}}{[Na_i]} * \left(1 + \frac{[K_i]}{K_{nak,ki}}\right)\right]} \quad (47)$$

$$\mathcal{P}\left(E^{\#}_{1,nak}\right) = \frac{1}{\left[1 + \frac{K_{nak,ki}}{[K_i]} * \left(1 + \frac{[Na_i]}{K_{nak,nai}}\right)\right]} \quad (48)$$

$$\mathcal{P}\left(E^{*}_{2,nak}\right) = \frac{1}{\left[1 + \frac{K_{nak,nae}}{Na_{eff}} * \left(1 + \frac{[K_e]}{K_{nak,ke}}\right)\right]} \quad (49)$$

$$\mathcal{P}\left(E^{\#}_{2,nak}\right) = \frac{1}{\left[1 + \frac{K_{nak,ke}}{[K_e]} * \left(1 + \frac{Na_{eff}}{K_{nak,nae}}\right)\right]} \quad (50)$$

$$Na_{eff} = [Na_e] * e^{(-0.82 * V_D)} \quad (51)$$

$$k_{1,nak} = \frac{0.37}{1 + \frac{0.094}{[ATP_i]}} \quad (52)$$

where, $K_{nak}$ is the maximal conductance for sodium-potassium ATPase, $[Na_i]$ is the intracellular concentration of sodium ion, $[Na_e]$ is the extracellular concentration of sodium ion, $[K_i]$ is the intracellular concentration of potassium ion, $[K_e]$ is the extracellular concentration of potassium ion, $\left(k_{1,nak}, k_{2,nak}, k_{3,nak}, k_{4,nak}\right)$ are the reaction rates,

$(K_{nak,nae}, K_{nak,nai}, K_{nak,ke}, K_{nak,ki})$ are the dissociation constants, $[ATP_i]$ is the intracellular concentration of adenosine triphosphate (ATP), and $V_D$ is the voltage defined thermodynamic entity.

The plasma membrane calcium ATPase $(I_{pmca})$ is given by,

$$I_{pmca} = K_{pc} * \left[k_{1,pc} * \mathcal{P}\left(E^*_{1,pc}\right) * y_{pc} - k_{2,pc} * \mathcal{P}\left(E^*_{2,pc}\right) * \left(1 - y_{pc}\right)\right] \tag{53}$$

$$\frac{d(y_{pc})}{dt} = \beta_{pc} * \left(1 - y_{pc}\right) - \alpha_{pc} * y_{pc} \tag{54}$$

$$\beta_{pc} = k_{2,pc} * \mathcal{P}\left(E^*_{2,pc}\right) + k_{4,pc} * \mathcal{P}\left(E_{2,pc}\right) \tag{55}$$

$$\alpha_{pc} = k_{1,pc} * \mathcal{P}\left(E^*_{1,pc}\right) + k_{3,pc} * \mathcal{P}\left(E_{1,pc}\right) \tag{56}$$

$$\mathcal{P}\left(E^*_{1,pc}\right) = \frac{1}{\left(1 + \frac{K_{pc,i}}{[Ca_i]}\right)}; \qquad \mathcal{P}\left(E^*_{2,pc}\right) = \frac{1}{\left(1 + \frac{K_{pc,e}}{[Ca_e]}\right)} \tag{57}$$

$$\mathcal{P}\left(E_{1,pc}\right) = 1 - \mathcal{P}\left(E^*_{1,pc}\right); \qquad \mathcal{P}\left(E_{2,pc}\right) = 1 - \mathcal{P}\left(E^*_{2,pc}\right) \tag{58}$$

$$k_{1,pc} = \frac{1}{1 + \frac{0.1}{[ATP_i]}} \tag{59}$$

$$K_{pc,i} = \left[\frac{173.6}{1 + \frac{[CaCam]}{5 * 10^{-5}}} + 6.4\right] * 10^{-5} \tag{60}$$

$$K_{pc} = k_{pmca} * \left[\frac{10.56 * [CaCam]}{[CaCam] + 5 * 10^{-5}} + 1.2\right] \tag{61}$$

where, $(k_{1,pc}, k_{2,pc}, k_{3,pc}, k_{4,pc})$ are the reaction rates, $k_{pmca}$ is the maximal conductance for calcium ATPase, $(K_{pc,e}, K_{pc,i})$ are the dissociation constants, $[ATP_i]$ is the intracellular concentration of ATP, $[Ca_i]$ is the intracellular calcium concentration, and $[CaCam]$ is the intracellular calcium-bound calmodulin concentration.

*Plasma Membrane Exchangers*

The plasma membrane sodium-calcium exchanger $(I_{NaCaX})$ is given by,

$$I_{NaCaX} = k_{xm} * \frac{[Na_i]^3 * [Ca_e] * exp^{(\delta_{xm}*V_D)} - [Na_e]^3 * [Ca_i] * e^{((\delta_{xm}-1)*V_D)}}{\left(1 + \mathcal{D}_{xm} * \left[[Na_i]^3 * [Ca_e] + [Na_e]^3 * [Ca_i]\right]\right) * \left(1 + \frac{[Ca_i]}{0.0069}\right)} \quad (62)$$

where, $k_{xm}$ is the maximal conductance for sodium-calcium exchanger, $[Na_e]$ is the extracellular sodium concentration, $[Na_i]$ is the intracellular sodium concentration, $[Ca_e]$ is the extracellular calcium concentration, $[Ca_i]$ is the intracellular calcium concentration, $\delta_{xm}$ is the energy barrier parameter, $\mathcal{D}_{xm}$ is the denominator factor, and $V_D$ is the voltage defined thermodynamic entity.

**Table S4.1:** Parameter values for ion-channel dynamics of SNc cell model (Muddapu and Chakravarthy, 2021).

| Constant | Symbol | Value | Units |
|---|---|---|---|
| Faraday's constant | $F$ | 96485 | $coulomb * mole^{-1}$ |
| SNc membrane capacitance | $C_{snc}$ | $9\ x\ 10^7$ | $pF * cm^{-2}$ |
| Cytosolic volume | $vol_{cyt}$ | $\phi_{cyt} * vol_{pmu}$ | $pl$ |
| Fraction of cytosolic volume | $\phi_{cyt}$ | 0.5 | $dimensionless$ |
| Pacemaking unit (PMU) volume | $vol_{pmu}$ | 5 | $pl$ |
| PMU area | $AR_{pmu}$ | $\delta_{pmu} * vol_{pmu}$ | $pl * cm^{-1}$ |
| PMU surface area-to-volume ratio | $\delta_{pmu}$ | $1.6667\ x\ 10^4$ | $cm^{-1}$ |
| Voltage defined thermodynamic entity | $V_D$ | $\frac{V}{V_\tau}$ | $dimensionless$ |

| | | | |
|---|---|---|---|
| Temperature defined thermodynamic entity | $V_\tau$ | $\frac{R * T}{F}$ | $mV$ |
| Universal gas constant | $R$ | 8314.472 | $mJ * mol^{-1} * K^{-1}$ |
| Physiological temperature | $T$ | 310.15 | $K$ |
| Maximal conductance of calcium channel | $\bar{g}_{Ca,L}$ | 2101.2 | $pA * mM^{-1}$ |
| Extracellular calcium concentration | $[Ca_e]$ | 1.8 | $mM$ |
| Reversal potential for calcium ion | $V_{Ca}$ | $\frac{1}{2} * \log\left(\frac{[Ca_e]}{[Ca_i]}\right)$ | $dimensionless$ |
| Valence of calcium ion | $z_{Ca}$ | 2 | $dimensionless$ |
| Maximal conductance of sodium channel | $\bar{g}_{Na}$ | 907.68 | $pA * mM^{-1}$ |
| Extracellular sodium concentration | $[Na_e]$ | 137 | $mM$ |
| Reversal potential for sodium ion | $V_{Na}$ | $\log\left(\frac{[Na_e]}{[Na_i]}\right)$ | $dimensionless$ |
| Valence of sodium ion | $z_{Na}$ | 1 | $dimensionless$ |
| Maximal conductance of sodium HCN channel | $\bar{g}_{NaHCN}$ | 51.1 | $pA * mM^{-1}$ |
| Maximal conductance of leaky sodium channel | $\bar{g}_{Nalk}$ | 0.0053 | $pA * mM^{-1}$ |
| Cyclic adenosine monophosphate concentration | $[cAMP]$ | $1 x 10^{-5}$ | $mM$ |
| Maximal conductance of delayed rectifying potassium channel | $\bar{g}_{Kdr}$ | 31.237 | $nS$ |

| | | | |
|---|---|---|---|
| Extracellular potassium concentration | $[K_e]$ | 5.4 | $mM$ |
| Reversal potential for potassium ion | $V_K$ | $\log\left(\frac{[K_e]}{[K_i]}\right)$ | $dimensionless$ |
| Valence of potassium ion | $z_K$ | 1 | $dimensionless$ |
| Maximal conductance of inward rectifying potassium channel | $\bar{g}_{Kir}$ | 13.816 | $nS$ |
| Maximal conductance of small conductance potassium channel | $\bar{g}_{Ksk}$ | 2.2515 | $pA * mM^{-1}$ |
| Maximal conductance for sodium-potassium ATPase | $K_{nak}$ | 1085.7 | $pA$ |
| Reaction rates of $I_{NaK}$ | $k_{2,nak}$ | 0.04 | $ms^{-1}$ |
| | $k_{3,nak}$ | 0.01 | $ms^{-1}$ |
| | $k_{4,nak}$ | 0.165 | $ms^{-1}$ |
| Dissociation constants of $I_{NaK}$ | $K_{nak,nae}$ | 69.8 | $mM$ |
| | $K_{nak,nai}$ | 4.05 | $mM$ |
| | $K_{nak,ke}$ | 0.258 | $mM$ |
| | $K_{nak,ki}$ | 32.88 | $mM$ |
| Maximal conductance for calcium ATPase | $k_{pmca}$ | 2.233 | $pA * ms^{-1}$ |
| Reaction rates of $I_{pmca}$ | $k_{2,pc}$ | 0.001 | $ms^{-1}$ |
| | $k_{3,pc}$ | 0.001 | $ms^{-1}$ |
| | $k_{4,pc}$ | 1 | $ms^{-1}$ |
| Dissociation constants of $I_{pmca}$ | $K_{pc,e}$ | 2 | $mM$ |

| | | | |
|---|---|---|---|
| Maximal conductance for sodium-calcium exchanger | $k_{xm}$ | 0.0166 | $pA * ms^{-1}$ |
| Energy barrier parameter of $I_{NaCaX}$ | $\delta_{xm}$ | 0.35 | $dimensionless$ |
| Denominator factor of $I_{NaCaX}$ | $\mathcal{D}_{xm}$ | 0.001 | $dimensionless$ |
| Calbindin reaction rates | $k_{1,calb}$ | 10 | $mM^{-1} * ms^{-1}$ |
| | $k_{2,calb}$ | $2\ x\ 10^{-3}$ | $ms^{-1}$ |
| Total cytosolic calbindin concentration | $[Calb_{tot}]$ | 0.005 | $mM$ |
| Calmodulin reaction rates | $k_{cam}^{cb}$ | 12000 | $mM^{-2} * ms^{-1}$ |
| | $k_{cam}^{nb}$ | $3.7\ x\ 10^{6}$ | $mM^{-2} * ms^{-1}$ |
| | $k_{cam}^{cd}$ | $3\ x\ 10^{-3}$ | $ms^{-1}$ |
| | $k_{cam}^{nd}$ | 3 | $ms^{-1}$ |
| Total cytosolic calmodulin concentration | $[Cam_{tot}]$ | 0.0235 | $mM$ |

**Table S4.2:** Steady state values of ion-channel dynamics of SNc cell model (Muddapu and Chakravarthy, 2021).

| **Symbol** | **Value** | **Symbol** | **Value** |
|---|---|---|---|
| $V_{SNc}$ | $-49.42\ mV$ | $h_{Na}$ | 0.1848 |
| $[Ca_i]$ | $1.88\ x\ 10^{-4}\ mM$ | $O_{NaHCN}$ | 0.003 |
| $[Na_i]$ | $4.69\ mM$ | $m_{K,dr}$ | 0.003 |
| $[K_i]$ | $126.06\ mM$ | $y_{nak}$ | 0.6213 |
| $m_{Na}$ | 0.0952 | $y_{pc}$ | 0.483 |

| | | | |
|---|---|---|---|
| $[Calb]$ | $26\,x\,10^{-4}\,mM$ | $[Cam]$ | $222\,x\,10^{-4}\,mM$ |

**S5. Biochemical Model of SNc Terminal**

The DA turnover process has been modelled as a three-compartment biochemical model based on Michaelis-Menten kinetics (Muddapu and Chakravarthy, 2021). The three compartments are intracellular compartment representing cytosol, extracellular compartment representing extracellular space (ECS), and vesicular compartment representing a vesicle. In DA turnover processes, L-tyrosine (TYR) is converted into L-3,4-dihydroxyphenylalanine or L-DOPA by tyrosine hydroxylase (TH), which in turn is converted into DA by aromatic L-amino acid decarboxylase (AADC) (Figure S5.1). The cytoplasmic DA ($DA_c$) is stored into vesicles by vesicular monoamine transporter 2 (VMAT-2) (Figure S5.2). Upon the arrival of an action potential, vesicular DA ($DA_v$) is released into extracellular space (Figure S5.3). Most of the extracellular DA ($DA_e$) is taken up into the terminal through DA plasma membrane transporter (DAT) (Figure S5.4), and remaining extracellular DA is metabolized by catechol-O-methyltransferase (COMT) and monoamine oxidase (MAO) into homovanillic acid (HVA) (Figure S5.5). The DA that enters the terminal is again packed into vesicles, and the remaining cytoplasmic DA is metabolized by COMT and MAO enzymes (Figure S5.5). It is known that a DA neuron self-regulates its firing, neurotransmission, and synthesis by autoreceptors (Anzalone et al., 2012; Ford, 2014). In the present model, we included autoreceptors that regulate the synthesis and release of DA (Figure S5.6, S5.7). Along with TYR, external L-DOPA compete for transporting into the terminal through aromatic L-amino acid transporter (AAT) (Figure S5.8).

*Modelling Extracellular DA in the ECS*

The major three mechanisms that determine the dynamics of extracellular DA ($[DA_e]$) in the ECS given by,

$$\frac{d([DA_e])}{dt} = J_{rel} - J_{DAT} - J_{eda}^{o} \tag{63}$$

where, $J_{rel}$ represents the flux of calcium-dependent DA release from the DA terminal, $J_{DAT}$ represents the unidirectional flux of DA translocated from the extracellular compartment (ECS) into the intracellular compartment (cytosol) via DA plasma membrane transporter (DAT), and $J^{o}_{eda}$ represents the outward flux of DA degradation, which clears DA from ECS.

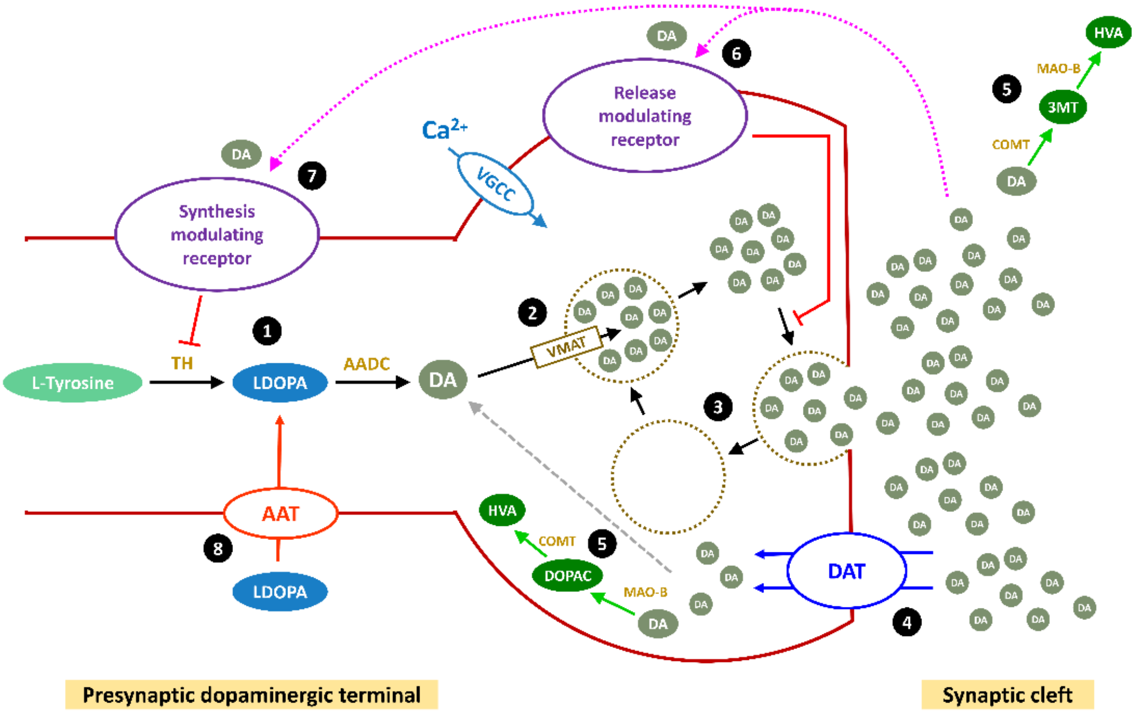


*Figure S5: Schematic of Dopamine turnover processes in SNc terminal (Muddapu and Chakravarthy, 2021).*

*Calcium-Dependent DA Release Flux*

Assuming that calcium-dependent DA release occurs within less than a millisecond after the calcium channels open, the flux of DA release ($J_{rel}$) from the DA terminal is given by,

$$J_{rel} = \psi * n_{RRP} * P_{rel}([Ca_i]) \quad (64)$$

where, $[Ca_i]$ is the intracellular calcium concentration in the DA terminal, $P_{rel}$ is the release probability as a function of intracellular calcium concentration, $n_{RRP}$ is the average number of readily releasable vesicles, and $\psi$ is the average release flux per vesicle within a single synapse.

The flux of calcium-dependent DA release depends on extracellular DA concentration, and intracellular ATP acts as a feedback mechanism, assuming this regulation as extracellular DA and intracellular ATP controls the number of vesicles in the readily releasable vesicle pool ($n_{RRP}$) which is given by,

$$n_{RRP} = \frac{\eta_{nrrp} * e^{\left(\frac{[ATP_i]}{K_{a,RRP}}\right)}}{\left(1 + e^{\left[\frac{-([DA_v]-[DA_{v_o}])}{DA_{v_s}}\right]}\right) * \left(1 + e^{\left[\frac{[DA_e]-DA_{R_a}}{DA_{R_s}}\right]}\right)} \quad (65)$$

$$\eta_{nrrp} = \bar{\eta}_{nrrp} - \beta_{nrrp,asyn_{mis}} * \left(\frac{1}{1 + \left(\frac{K_{asyn_{mis}}}{[ASYN_{mis}]}\right)^4}\right) \quad (66)$$

where, $[DA_{v_o}]$ is the initial vesicular DA concentration, $DA_{v_s}$ is the sensitivity to vesicular concentration, $DA_{R_a}$ is the high-affinity state for DA binding to receptors and $DA_{R_s}$ is the binding sensitivity, $[ATP_i]$ is the intracellular ATP concentration, $K_{a,RRP}$ is the activation constant for ATP, $\eta_{nrrp}$ is the effect of misfolded alpha-synuclein on vesicle recycling (Venda et al., 2010), $\bar{\eta}_{nrrp}$ is the maximal vesicle recycling efficiency, $\beta_{nrrp,asyn_{mis}}$ is the maximum fractional decrease in the vesicle recycling efficiency through $ASYN_{mis}$, $K_{asyn_{mis}}$ is the threshold concentration for damage by $ASYN_{mis}$, and $[ASYN_{mis}]$ is the misfolded alpha-synuclein concentration.

The release probability of DA as a function of intracellular calcium concentration is given by,

$$P_{rel}([Ca_i]) = \bar{P}_{rel} * \frac{[Ca_i]^4}{[Ca_i]^4 + K_{rel}^4} \quad (67)$$

where, $\bar{P}_{rel}$ is the maximum release probability and $K_{rel}$ is the sensitivity of calcium concentration, and $[Ca_i]$ is the intracellular calcium concentration.

*Unidirectional Reuptake Flux of DA*

The unidirectional reuptake flux of extracellular DA into the presynaptic terminal is given by,

$$J_{DAT} = \bar{V}_{eda} * \frac{[DA_e]}{K_{eda} + [DA_e]} \quad (68)$$

where, $\bar{V}_{eda}$ is the maximal velocity of DA transporter (DAT), $K_{eda}$ is the DA concentration at half-maximal velocity, and $[DA_e]$ is the extracellular DA concentration.

*Outward Extracellular Flux*

The flux of extracellular DA enzymatic degradation in the synaptic cleft (ECS) is given by,

$$J_{eda}^{o} = k_{comt} * [DA_e] \tag{69}$$

where, $k_{comt}$ is the rate at which extracellular DA cleared from ECS, and $[DA_e]$ is the extracellular DA concentration.

*<u>Modelling Intracellular DA in the Terminal</u>*

The intracellular DA dynamics ($[DA_i]$) is determined as the sum of DA concentration in cytosolic and vesicular compartments and is given by,

$$\frac{d([DA_i])}{dt} = \frac{d([DA_c])}{dt} + \frac{d([DA_v])}{dt} \tag{70}$$

The cytosolic DA dynamics ($[DA_c]$) is given by,

$$\frac{d([DA_c])}{dt} = J_{DAT} - J_{VMAT} - J_{cda}^{o} + J_{ldopa} \tag{71}$$

where, $J_{DAT}$ represents the unidirectional flux of DA translocated from ECS into the cytosol through DAT, $J_{VMAT}$ represents the flux of cytosolic DA into vesicle through VMAT-2, $J_{ida}^{o}$ represents the outward flux of DA degradation, which clears DA from the cytosol, and $J_{ldopa}$ represents the flux of synthesized cytosol DA from L-DOPA.

The vesicular DA dynamics ($[DA_v]$) is given by,

$$\frac{d([DA_v])}{dt} = J_{VMAT} - J_{rel} \tag{72}$$

where, $J_{rel}$ represents the flux of calcium-dependent DA release from the DA terminal, $J_{VMAT}$ represents the flux of cytosolic DA into a vesicle.

*L-DOPA Synthesis Flux*

The flux of synthesized L-DOPA whose velocity is the function of intracellular calcium concentration and L-DOPA synthesis is regulated by the substrate (TYR) itself, extracellular DA (via autoreceptor) and intracellular DA concentrations are given by,

$$J_{synt} = \frac{V_{synt}}{1 + \frac{K_{TYR}}{[TYR]} * \left(1 + \frac{[DA_c]}{K_{i,cda}} + \frac{[DA_e]}{K_{i,eda}}\right)} \tag{73}$$

where, $V_{synt}$ is the velocity of synthesizing L-DOPA, $[TYR]$ is the tyrosine concentration in terminal bouton, $K_{TYR}$ is the tyrosine concentration at which half-maximal velocity was attained, $K_{i,cda}$ is the inhibition constant on $K_{TYR}$ due to cytosolic DA concentration, $K_{i,eda}$ is the inhibition constant on $K_{TYR}$ due to extracellular DA concentration, $[DA_c]$ is the cytoplasmic DA concentration, and $[DA_e]$ is the extracellular DA concentration.

In Chen et al.(Chen et al., 2003), neuronal stimulation was linked to DA synthesis through an indirect event, which starts with calcium influx into the terminal bouton. In this model, the velocity of L-DOPA synthesis as a function of calcium levels in the terminal bouton is expressed as,

$$V_{synt}(Ca_i) = \bar{V}_{synt} * \frac{[Ca_i]^4}{K_{synt}^4 + [Ca_i]^4} \tag{74}$$

where, $K_{synt}$ is the calcium sensitivity, $\bar{V}_{synt}$ is the maximal velocity for L-DOPA synthesis, and $[Ca_i]$ is the intracellular calcium concentration.

*Storage Flux of DA into the Vesicle*

The flux of transporting DA in the cytosol into the vesicles, which depends on the intracellular ATP is given by,

$$J_{VMAT} = V_{cda,ATP} * \frac{[DA_c]}{K_{cda} + [DA_c]} \tag{75}$$

$$V_{cda,ATP} = \bar{V}_{cda} * \alpha_{vmat} * e^{(\beta_{vmat} * [ATP_i])} \tag{76}$$

where, $K_{cda}$ is the cytosolic DA concentration at which half-maximal velocity was attained, $\bar{V}_{cda}$ is the maximal velocity with which DA was packed into vesicles, $[DA_c]$ is the cytosolic

DA concentration, $\alpha_{vmat}$ is the scaling factor for VMAT-2, $\beta_{vmat}$ is the scaling factor for $ATP_i$, and $[ATP_i]$ is the intracellular ATP concentration.

*Outward Intracellular Flux*

The flux of intracellular DA enzymatic degradation in synaptic bouton (cytosol) is given by,

$$J_{cda}^{o} = k_{mao} * [DA_c] \tag{77}$$

where, $k_{mao}$ is the rate at which intracellular DA cleared from the cytosol, and $[DA_c]$ is the cytosolic DA concentration.

*L-DOPA to DA Conversion Flux*

The flux of L-DOPA conversion to DA by AADC(Reed et al., 2012) is given by,

$$J_{ldopa} = \bar{V}_{aadc} * \frac{[LDOPA]}{K_{aadc} + [LDOPA]} \tag{78}$$

where, $K_{aadc}$ is the L-DOPA concentration at which half-maximal velocity was attained, $\bar{V}_{aadc}$ is the maximal velocity with which L-DOPA was converted to DA, $[LDOPA]$ is the L-DOPA concentration.

*Transport Flux of Exogenous L-DOPA into the Terminal*

The flux of exogenous L-DOPA transported into the terminal through AAT while competing with other aromatic amino acids (Reed et al., 2012) is given by,

$$J_{aat} = \bar{V}_{aat} * \frac{[LDOPA_e]}{\left(K_{ldopa_e} * \left(1 + \left(\frac{[TYR_e]}{K_{tyr_e}}\right) + \left(\frac{[TRP_e]}{K_{trp_e}}\right)\right) + [LDOPA_e]\right)} \tag{79}$$

where, $K_{ldopa_e}$ is the extracellular L-DOPA concentration at which half-maximal velocity was attained, $\bar{V}_{aat}$ is the maximal velocity with which extracellular L-DOPA was transported into the cytosol, $[LDOPA_e]$ is the extracellular L-DOPA concentration, $[TYR_e]$ is the extracellular TYR concentration, $[TRP_e]$ is the extracellular tryptophan (TRP) concentration, $K_{tyr_e}$ is the affinity constant for $[TYR_e]$, $K_{trp_e}$ is the affinity constant for $[TRP_e]$.

When L-DOPA drug therapy is initiated,

$$[LDOPA_e] = [sLD] \quad (80)$$

When no L-DOPA drug therapy is initiated,

$$LDOPA_e = 0 \quad (81)$$

The L-DOPA concentration ($[LDOPA]$) dynamics inside the terminal is given by,

$$\frac{d([LDOPA])}{dt} = J_{aat} - J_{ldopa} + J_{synt} \quad (82)$$

where, $J_{aat}$ represents the flux of exogenous L-DOPA transported into the cytosol, $J_{ldopa}$ represents the conversion flux of exogenous L-DOPA into DA, $J_{synt}$ represents the flux of synthesized LDOPA from tyrosine, and $[sLD]$ is the serum L-DOPA concentration.

**Table S5.1:** Parameter values for DA turnover processes of SNc cell model (Muddapu and Chakravarthy, 2021).

| Constant | Symbol | Value | Units |
|---|---|---|---|
| Average release flux per vesicle | $\psi$ | 17.4391793 | $mM * ms^{-1}$ |
| Initial vesicular DA concentration | $DA_{v_o}$ | 500 | $mM$ |
| Sensitivity to vesicular DA concentration | $DA_{v_s}$ | 0.01 | $mM$ |
| Affinity constant of DA binding to receptors | $DA_{R_a}$ | $5 \, x \, 10^{-5}$ | $mM$ |
| Binding sensitivity | $DA_{R_s}$ | 0.01 | $mM$ |
| Activation constant for ATP | $K_{a,RRP}$ | 1.4286 | $mM$ |
| Vesicle recycling maximal flux | $\bar{v}_{nrrp}$ | $1 \, x \, 10^{-3}$ | $mM * ms^{-1}$ |
| Maximal vesicle recycling efficiency | $\bar{\eta}_{nrrp}$ | 0.995 | $dimensionless$ |
| Maximal fraction of $asyn^*$ effect on the vesicle | $\beta_{nrrp,asyn_{mis}}$ | 0.08 | $dimensionless$ |

| | | | |
|---|---|---|---|
| Affinity constant for $asyn^*$ | $K_{asyn_{mis}}$ | $8.5\ x\ 10^{-3}$ | $mM$ |
| Reaction constant of $DA_e$ clearance | $k_{comt}$ | 0.0083511 | $ms^{-1}$ |
| Tyrosine concentration | $[TYR]$ | $126\ x\ 10^{-3}$ | $mM$ |
| Affinity constant for $TYR$ | $K_{TYR}$ | $46\ x\ 10^{-3}$ | $mM$ |
| Inhibition constant for $DA_c$ | $K_{i,cda}$ | $11\ x\ 10^{-2}$ | $mM$ |
| Inhibition constant for $DA_e$ | $K_{i,eda}$ | $46\ x\ 10^{-3}$ | $mM$ |
| Maximal velocity of DA synthesis | $\bar{V}_{synt}$ | $25\ x\ 10^{-6}$ | $mM * ms^{-1}$ |
| Affinity constant for $Ca_i$ | $K_{synt}$ | $35\ x\ 10^{-4}$ | $mM$ |
| Maximal velocity of VMAT | $\bar{V}_{cda}$ | $4.67\ x\ 10^{-6}$ | $ms^{-1}$ |
| Affinity constant for $DA_c$ | $K_{cda}$ | $238\ x\ 10^{-4}$ | $mM$ |
| Scaling factor for VMAT | $\alpha_{vmat}$ | $1\ x\ 10^{-3}$ | $dimensionless$ |
| Scaling factor for $ATP_i$ | $\beta_{vmat}$ | 3 | $dimensionless$ |
| Reaction constant of $DA_c$ clearance | $k_{mao}$ | 0.00016 | $ms^{-1}$ |
| Maximal velocity of AADC | $\bar{V}_{aadc}$ | $9.73\ x\ 10^{-5}$ | $mM * ms^{-1}$ |
| Affinity constant for $LDOPA$ | $K_{aadc}$ | 0.13 | $mM$ |
| Maximal velocity of AAT | $\bar{V}_{aat}$ | $5.11\ x\ 10^{-7}$ | $mM * ms^{-1}$ |
| Affinity constant for $LDOPA_e$ | $K_{ldopa_e}$ | $3.2\ x\ 10^{-4}$ | $mM$ |
| Affinity constant for $TYR_e$ | $K_{tyr_e}$ | $6.4\ x\ 10^{-4}$ | $mM$ |
| Affinity constant for $TRP_e$ | $K_{trp_e}$ | $1.5\ x\ 10^{-4}$ | $mM$ |
| Serum concentration of TYR | $[TYR_e]$ | $6.3\ x\ 10^{-4}$ | $mM$ |
| Serum concentration of TRP | $[TRP_e]$ | $8.2\ x\ 10^{-4}$ | $mM$ |
| Serum concentration of LDOPA | $[sLD]$ | $3.6\ x\ 10^{-3}$ | $mM$ |

**Table S5.2:** Steady state values of DA turnover processes of SNc cell model (Muddapu and Chakravarthy, 2021).

| **Symbol** | **Value** | **Symbol** | **Value** |
|---|---|---|---|
| $[DA_e]$ | $4\ x\ 10^{-6}\ mM$ | $[DA_v]$ | $500\ mM$ |
| $[DA_c]$ | $1\ x\ 10^{-4}\ mM$ | $[LDOPA]$ | $3.6\ x\ 10^{-4}\ mM$ |

### S6. Biochemical Model of Dopaminergic Terminal

The dopaminergic terminal model (Reed et al., 2012) includes: transport of tyrosine across the BBB and into the terminal; synthesis of LD by tyrosine hydroxylase (TH), synthesis of cytosolic DA by AADC, packaging of cytosolic DA into vesicles by VMAT, release of vesicular DA into the extracellular space depending on firing rate, reuptake of extracellular DA into the cytosol by the dopamine transporters (DATs), diffusion of extracellular DA out of the system, catabolism of DA in both the extracellular space and the cytosol by monoamine oxidase (MAO), and the effects of extracellular DA on DA synthesis and release via the auto-receptors.

The L-DOPA concentration ($[LDOPA]$) dynamics inside the terminal is given by Equation 84 below as updated in main manuscript,

$$\frac{d([LDOPA])}{dt} = J_{aat} - J_{ldopa} + J_{synt} \tag{83}$$

Where $J_{aat}$ represents the flux of exogenous L-DOPA ($[LDOPA_{CC}]$) transported into the cytosol, $J_{synt}$ represents the flux of synthesized L-DOPA from tyrosine given by, and $J_{ldopa}$ represents the conversion flux of exogenous L-DOPA ($[LDOPA_{CC}]$) into dopamine, as given in Equations 84-87 below,

$$J_{aat} = \bar{V}_{aat} * \frac{[LDOPA_{CC}]}{\left(K_{ldopa_e} * \left(1 + \left(\frac{[TYR_e]}{K_{tyr_e}}\right) + \left(\frac{[TRP_e]}{K_{trp_e}}\right)\right) + [LDOPA_{CC}]\right)} \tag{84}$$

$J_{synt}$ represents the flux of synthesized L-DOPA from tyrosine given by,

$$J_{ldopa} = \bar{V}_{aadc} * \frac{[LDOPA]}{K_{aadc} + [LDOPA]} \tag{85}$$

$$J_{synt} = \bar{V}_{aat}\left[\frac{0.56}{1} + \frac{TYR_e}{K_{tyre}}\right][\frac{4.5}{8\left(\frac{DA_e}{0.002024}\right)^4 + 1} + 0.5] \quad (86)$$

$$\frac{d(DA_c)}{dt} = J_{ldopa} - J_{MAT} + J_{DAT} - k_{CATAB}^{cda}.[DA_c] \quad (87)$$

$$\frac{d(DA_v)}{dt} = J_{MAT} - [DA_v].fire(t) \quad (88)$$

$$\frac{d(DA_e)}{dt} = [DA_v].fire(t) + J_{DAT} + J_{CATAB}^{DA_e} - k_{rem}.[DA_e] + J_{ldopa}^{ext} \quad (89)$$

where $[DA_c]$ is the cytosolic dopamine concentration, $[DA_v]$ is the concentration of vesicular dopamine, $J_{MAT}$ represents the flux of the cytosolic dopamine that is packed into vesicles, $J_{DAT}$ represents the flux of the extracellular dopamine that is reuptake by the terminal, $vda$ is the concentration of the vesicular dopamine, $fire(t)$ represents the firing of the neuron at time instance 't', $[DA_e]$ represents the concentration of extracellular dopamine, $k_{rem}$ is the affinity for the removal of extracellular dopamine, $J_{CATAB}^{DA_e}$ represents the flux of the dopamine that is catabolized in extracellular space and $J_{CATAB}^{cda}$ represents the flux of dopamine that is catabolized inside the terminal.

The effective dopamine released into extracellular space due to medication effect is given be subtracting the basal value of $DA_e^{basal}$ from the total $[DA_e]$, which includes the medication effect. From the trials in our model $DA_e^{basal}$ was found to be 2 nM.

$$DA_e^{med} = DA_e - DA_e^{basal} \quad (90)$$

And some of the dopamine released into the extracellular space is catabolized into homovanillic acid.

$$\frac{d(hva)}{dt} = k_{cda}^{catab}.[DA_c] + J_{CATAB}^{DA_e} - k_{hva}^{catab}.hva \quad (91)$$

**Table S6.2:** Steady state values of DA turnover processes of SNc cell model (Best et al., 2009; Reed et al., 2012; Muddapu and Chakravarthy, 2021).

| Constant | Symbol | Value | Units |
|---|---|---|---|

| | | | |
|---|---|---|---|
| Basal value of Extracellular Dopamine released. | $DA_e^{basal}$ | 2 | $nM$ |
| Removal rate of Extracellular Dopamine | $k_{rem}$ | 400 | $hr^{-1}$ |
| Rate constant of dopamine catabolized inside the terminal | $k_{CATAB}^{cda}$ | 10 | $hr^{-1}$ |
| Rate constant of dopamine catabolized in extracellular space | $k_{hva}^{CATAB}$ | 0.02 | $\mu M hr^{-1}$ |
| Maximal velocity of AAT | $\bar{V}_{aat}$ | $5.11\ x\ 10^{-7}$ | $mM * ms^{-1}$ |

### S7. Levodopa Medication Curve

A three compartmental model of levodopa (L-DOPA) administration (Baston et al., 2016; Véronneau-Veilleux et al., 2020) was integrated to the multiscale large scale basal ganglia model. The compartmental model simulates the condition where the drug is orally consumed by the person, it is absorbed in the bloodstream, and after interacting with other bodily fluids, a proportion of L-DOPA crosses the blood-brain barrier (BBB) and gets absorbed in the terminals. This L-DOPA is added up to the L-DOPA synthesized from tyrosine using the enzyme tyrosine hydroxylase. For a 100 mg of L-DOPA Dosage the amount of extracellular dopamine released due to medication effect follows the curve shown in the Figure S7.1 below. The different curves shown represents the $DA_e^{med}$ released with increasing percentage of cell loss. $DA_e$ released is observed for different dosages and terminal loss as shown in the Figure S7.1.

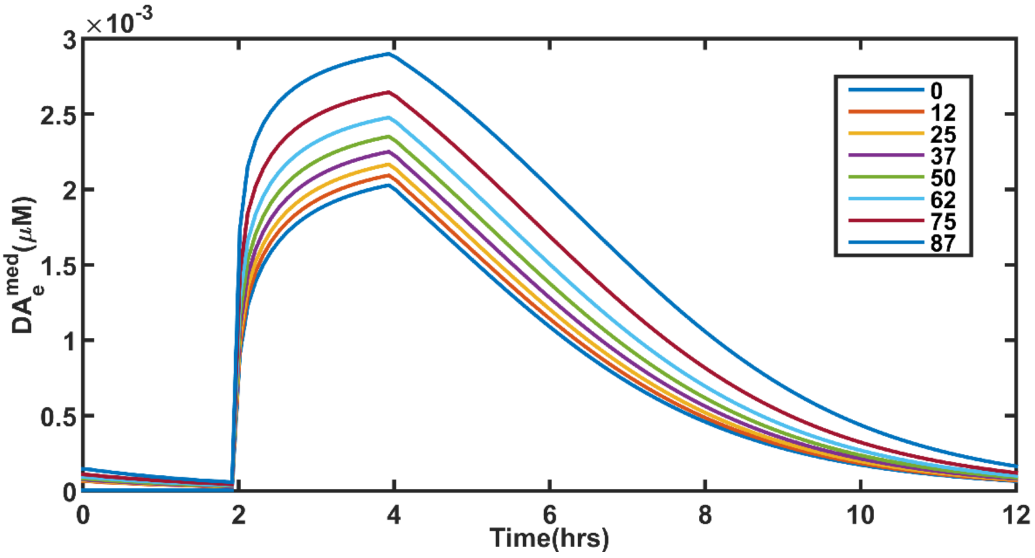


*Figure S7.1: The figure shows the concentration of $DA_e^{med}$ released after consuming 100 mg of L-DOPA with varying percentages of SNc terminals getting killed.*

### S7.1 Variation of Peak $DA_e^{med}$ released with varying L-DOPA Dosages and SNc Terminal Loss

As the number of SNc terminals is lost, there are fewer terminals to absorb and reuptake the excess L-DOPA present in the extracellular space. Therefore, we see increased $DA_e^{med}$ peaks when the dopaminergic terminals are lost. Similarly, as the dosage increased higher concentration of L-DOPA crosses the BBB, and more amount of $DA_e^{med}$ is released (Figure S7.2(A)). Figure S7.2(B) shows the effect of dosage and the terminal loss on peak extracellular DA.

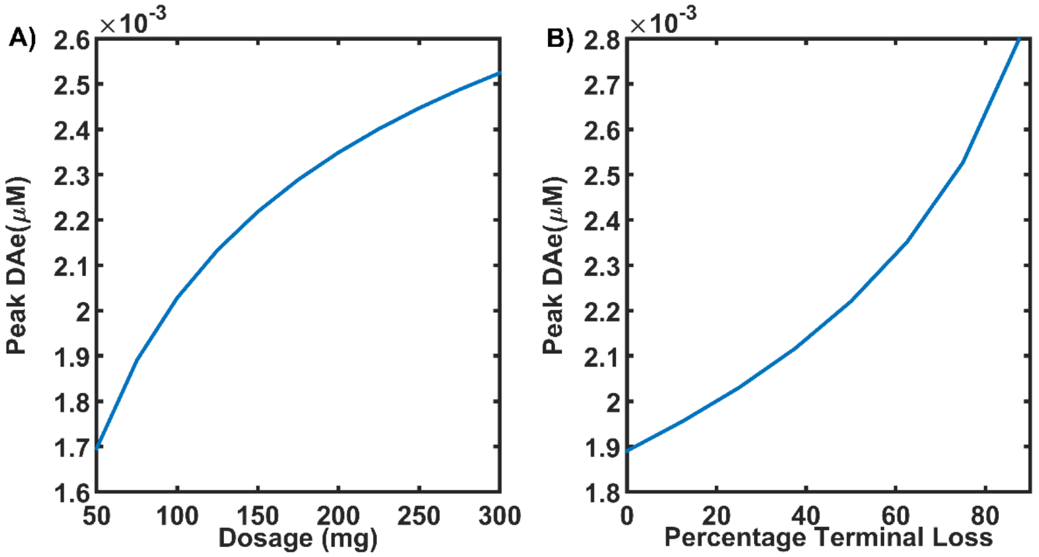


*Figure S7.2: The figure shows the peak value $DA_e^{med}$ released after consuming 100 mg of L-DOPA with varying dosages with no terminal loss (A) and varying percentages of SNc terminals loss (B).*

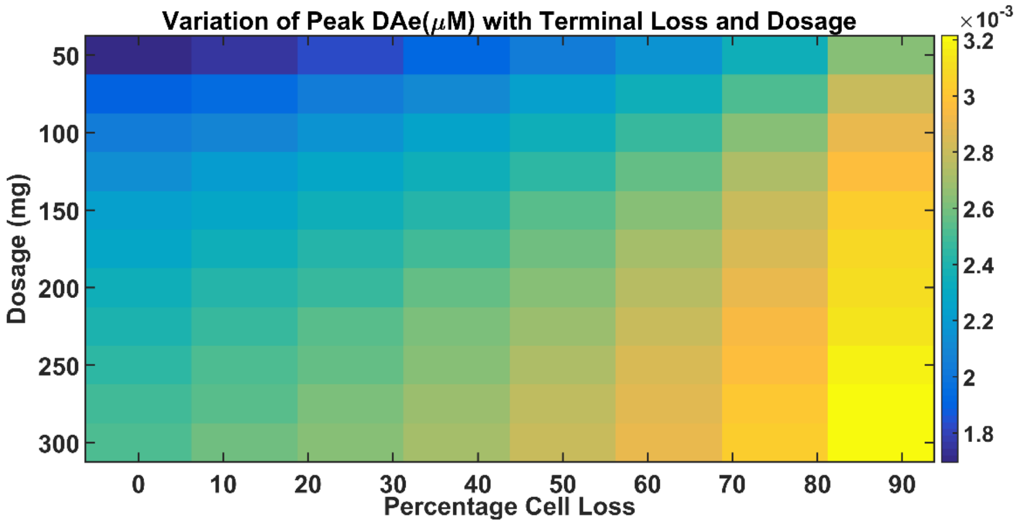


***Figure S7.3: The figure shows the peak value $DA_e^{med}$released with varying percentages of SNc terminals loss (x-axis) and L-DOPA dosages (y-axis).***

Figure S7.2(A) and S7.2(B) shows the effect of SNc terminal loss and the increasing dosages on the concentration of $DA_e$ released whereas the Figure S7.3 shows the combined effect of both.

For analysing the performance of reaching performance reaching task was carried out at various time points after the L-DOPA medication. The various time instances after the medication were taken by sampling the L-DOPA medication curve as shown in Figure S7.4 below.

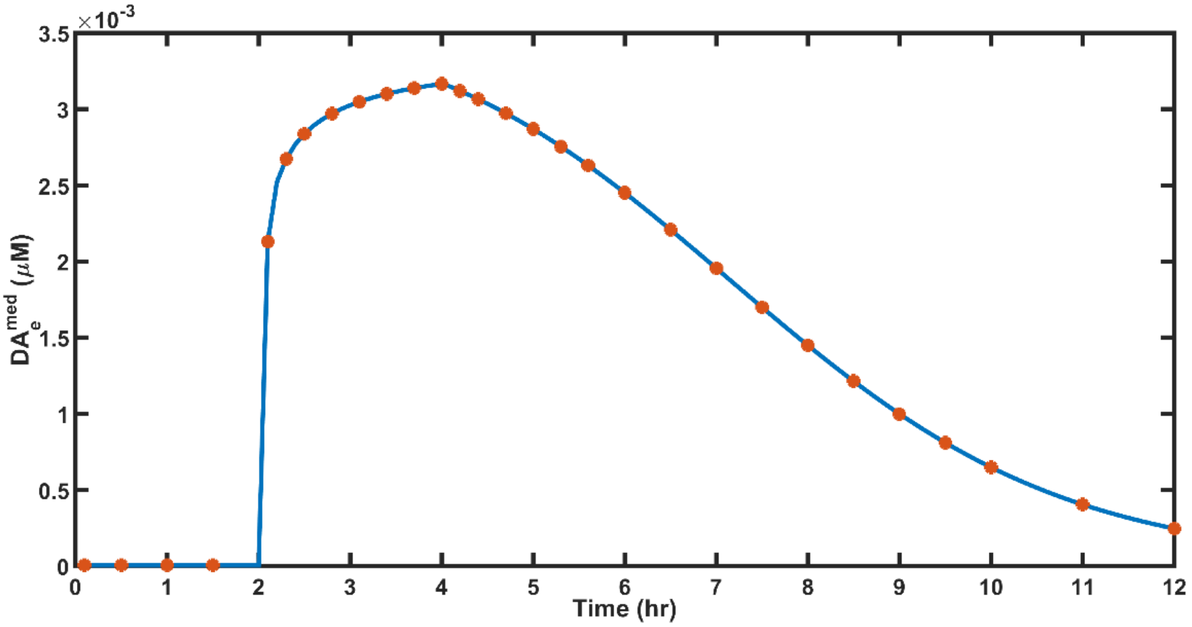

*Figure S7.4: The figure shows the different sample points taken along the $DA_e^{med}$ curve that is taken as a reference for testing the reaching performance after different times following the consumption of the drug. The medication curve is plotted for 100 mg dosage of L-DOPA without any terminal loss.*

### S7.2 Levodopa Medication Effect on Arm Reaching Performance

In this section, the performance of the arm reaching task was evaluated with L-DOPA medication and how the terminal loss and the increasing dosages impact the performance. The L-DOPA dosages ranging from 50 mg to 300 mg in steps of 50 mg are tested in our model with varying cell loss ranging from 0% to 87.5% in steps of 12.5. Here we consider two scenarios a) where dopaminergic loss impacts striatum alone and b) where dopaminergic loss impacts both striatum and STN.

a) With DA loss impacting only the striatum

In this scenario, the dopaminergic loss in the nigrostriatal pathway leads to BG operating predominantly via the indirect pathway resulting in reduced performance. Below Figure S7.5 and S7.6 show the reaching performance with various cell losses (0, 12.5, 25, 37.5, 50, 62.5, 75, 87.5) for L-DOPA dosages 100 and 250 mg, respectively.

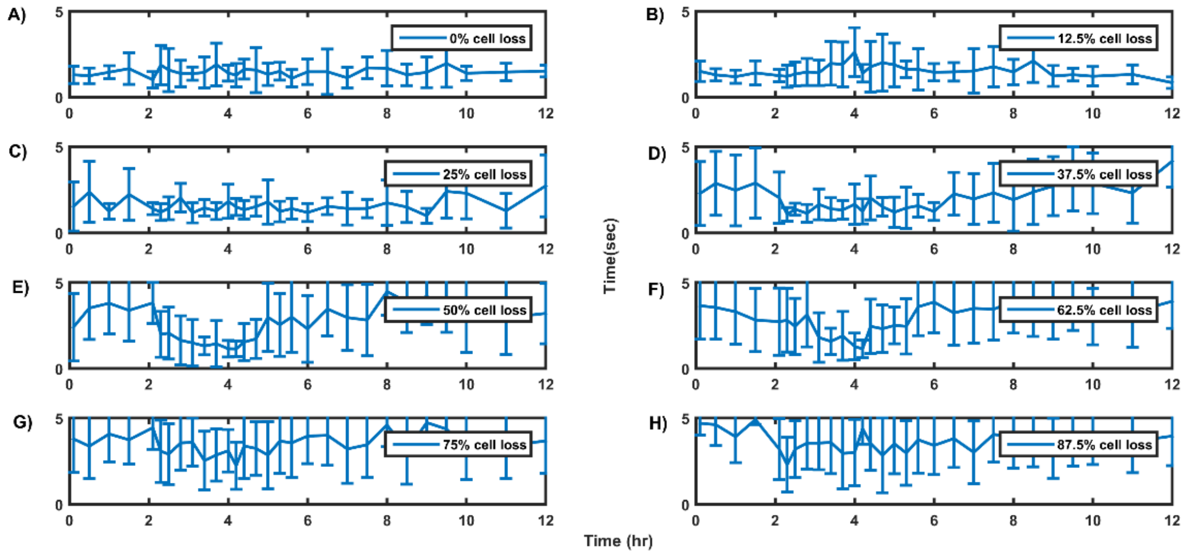


*Figure S7.5: The figure shows the average time to reach the target measured across 10 trials for varying percentage of SNc cell loss impacting only the striatum. The performance metrics is observed for the L-DOPA dosage of 100 mg and reaching performance measured at various times during the 12-hour window where L-DOPA drug is administered.*

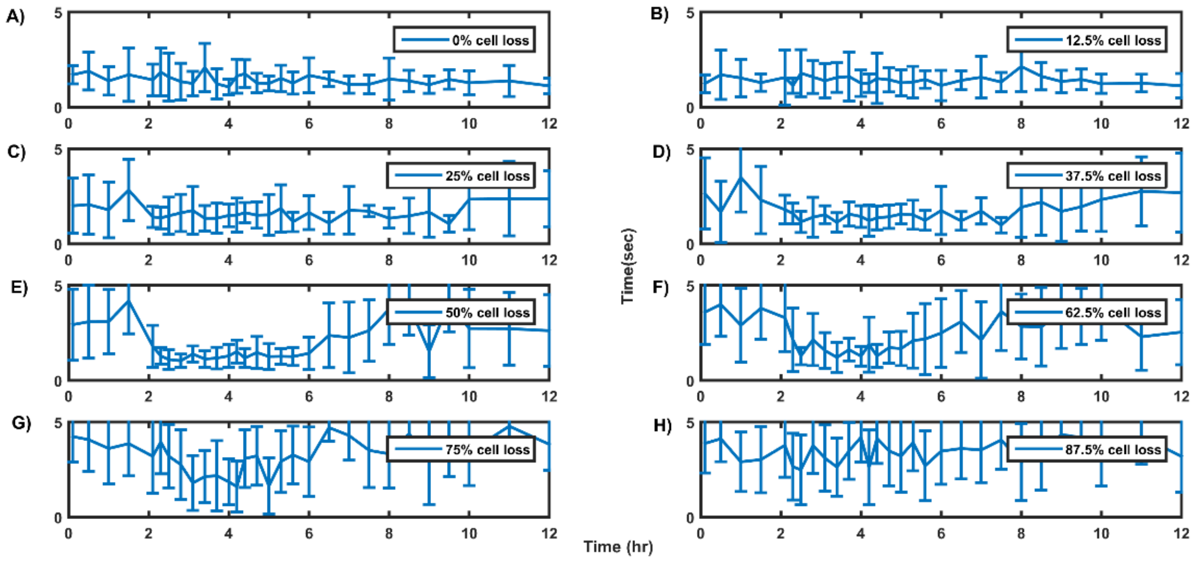


*Figure S7.6: The figure shows the average time to reach the target measured across 10 trials for varying percentage of cell loss impacting only the striatum. The performance metrics is observed for the L-DOPA dosage of 250 mg and reaching performance measured at various times during the 12-hour window where the L-DOPA drug is administered.*

From both the plots, we can see that as the cell loss increases from 12.5% to 87.5 %, the therapeutic effect window keeps reducing. Referring to Figure S7.6, for 25% cell loss (C) the therapeutic effect was nearly there for 8 hrs duration. When the cell loss increases to 37.5% (D) the therapeutic effect came down to 6 hrs, and it further reduced to 4 hrs with 50% cell loss (E). The effect of an increase in dosage can also be noticed in Figure S7.5 and Figure S7.6. At 50% cell loss with L-DOPA dosage 100 mg (Figure S7.5(E)), the therapeutic effect was ~ 3 hrs, whereas when the dosage increased to 250 mg (Figure S7.6(E)), the therapeutic effect was ~ 4 hrs.

b) With DA loss impacting both striatum and STN

In this scenario, both the dopaminergic axonal loss in the nigrostriatal and nigrosubthalamic pathways occurs. As a result, in addition to the reduced supply of dopamine to the striatum, it also results in weak lateral strengths among STN neurons. Below Figure S7.7 and S7.8 show the reaching performance with various cell losses for L-DOPA dosages 100 and 250 mg, respectively.

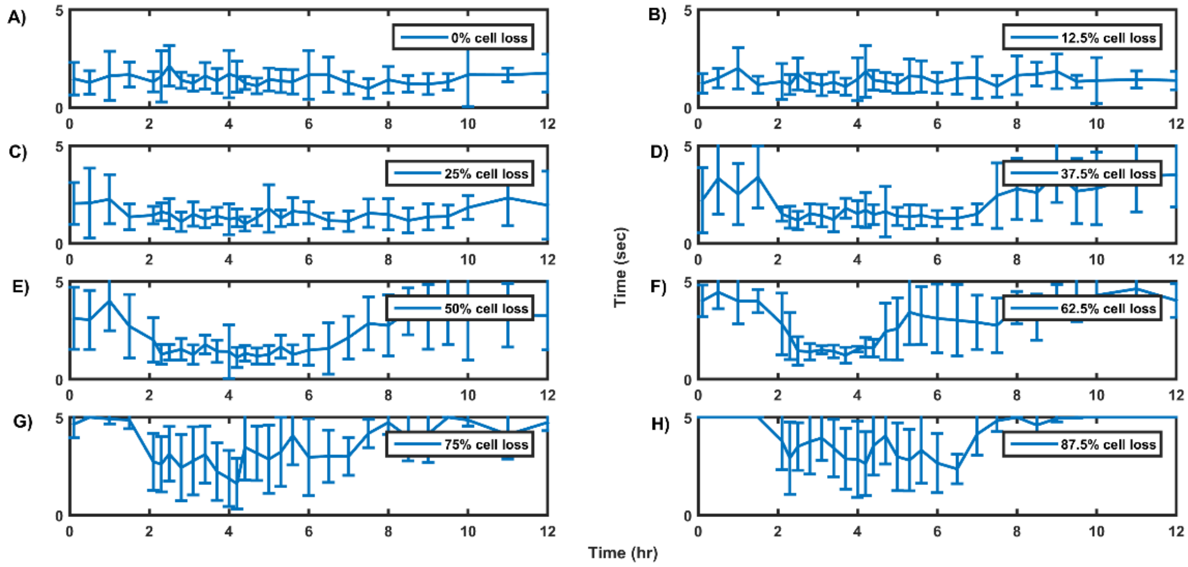


*Figure S7.7 The figure shows the average time to reach the target measured across 10 trials for varying percentage of cell loss impacting both the striatum and STN. The performance metrics is observed for the L-DOPA dosage of 100 mg and reaching performance measured at various times during the 12-hour window where the L-DOPA drug is administered.*

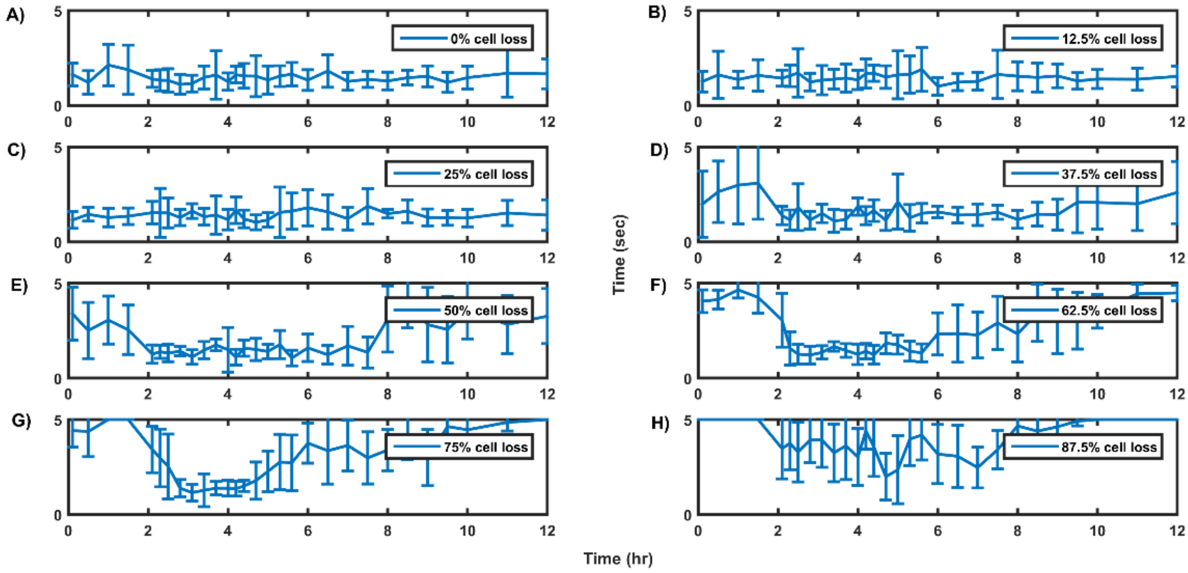


*Figure S7.8: The figure shows the average time to reach the target measured across 10 trials for varying percentage of cell loss impacting both the striatum and STN. The performance metrics is observed for the L-DOPA dosage of 250 mg and reaching performance measured at various times during the 12-hour window where the L-DOPA drug is administered*

From Figure S7.7 and S7.8, we can also see that the performance deteriorates with increasing cell loss. With reference to Figure S7.8(E), we can see that the therapeutic duration for 50% cell loss case under 250 mg L-DOPA dosage was about 6 hours, whereas when the cell loss increased to 75%, the therapeutic window dropped to nearly 2 hours. It is also observed that as the dosage increases, the therapeutic window increases. In reference to Figure S7.7 (F), at 62.5% cell loss with 100 mg L-DOPA dosage, the therapeutic effect was ~ 2.5 hrs, whereas when the dosage was increased to 250 mg, the therapeutic effect increased to ~ 4 hrs.

The performance results for the other dosages (50, 150, 200, and 300 mg) are summarized as shown in Figures S7.9, S7.10, S7.11, and S7.12 below. For simplicity, we have demonstrated the reaching performance across four percentage of cell loss (0%, 25%, 50%, and 75%). The left-hand side plots (A, C, E, and G) indicated the performance when the cell loss affects only the striatum, whereas the right-hand side plots (B, D, F, and H) indicate the performance when the cell loss impacts both striatum and STN.

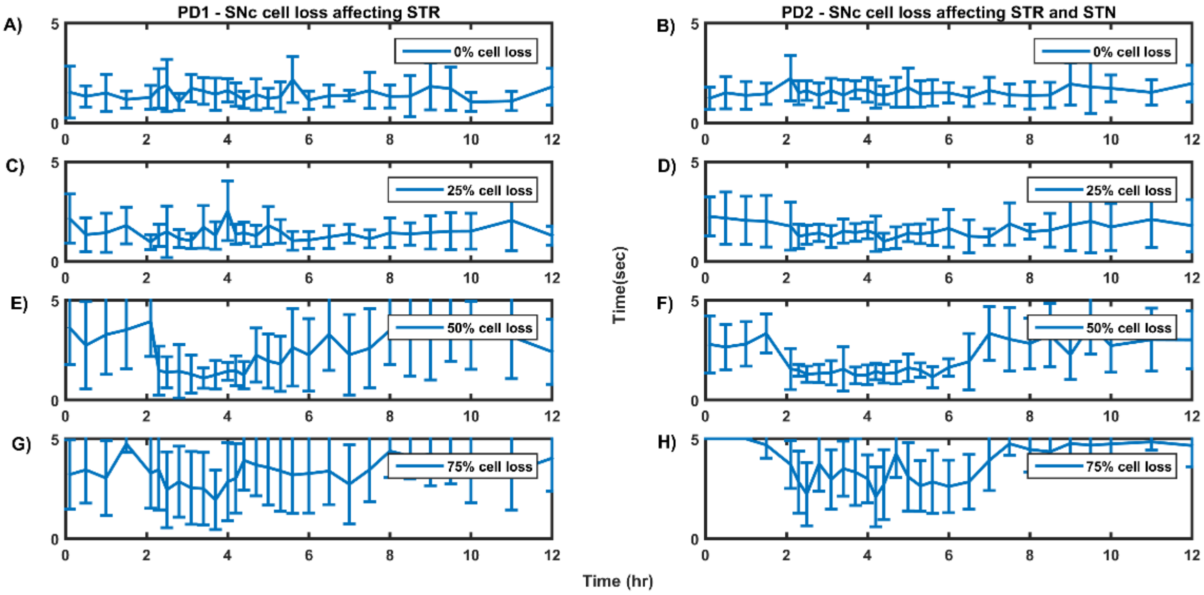


*Figure S7.9: The figure shows the average time to reach the target measured across 10 trials for varying percentage of cell loss impacting only the striatum (A, C, E, G) and impacting both striatum and STN (B, D, F, H). The performance metrics is observed for the L-DOPA dosage of 50 mg and reaching performance measured at various times during the 12-hour window where the L-DOPA drug is administered.*

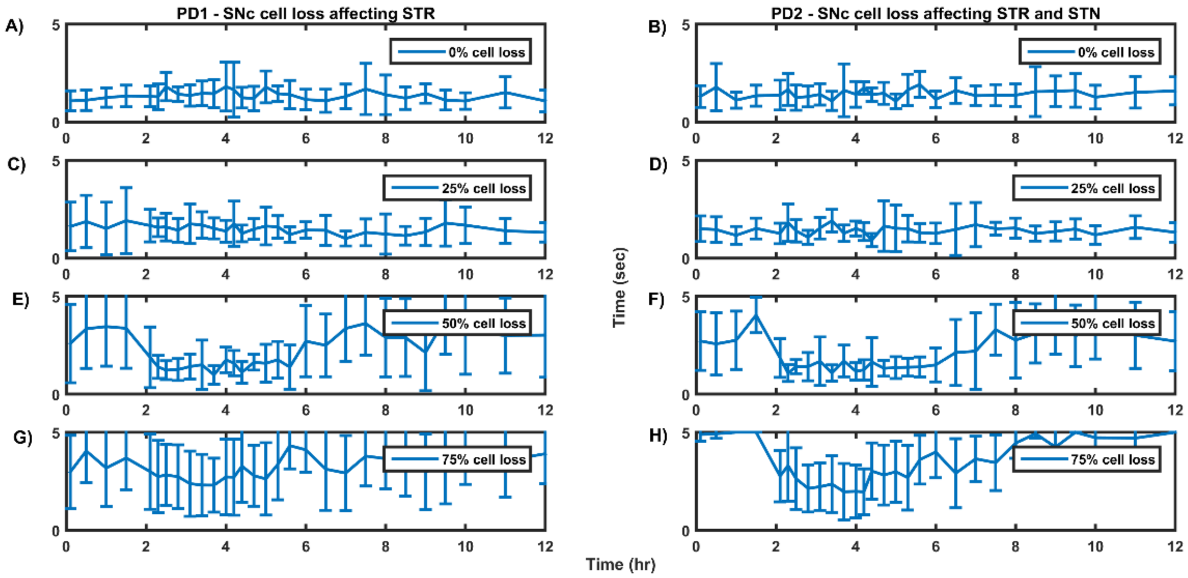


*Figure S7.10: The figure shows the average time to reach the target measured across 10 trials for varying percentage of cell loss impacting only the striatum (A, C, E, G) and impacting both striatum and STN (B, D, F, H). The performance metrics is observed for the L-DOPA dosage of 150mg and reaching performance measured at various times during the 12-hour window where the L-DOPA drug is administered.*

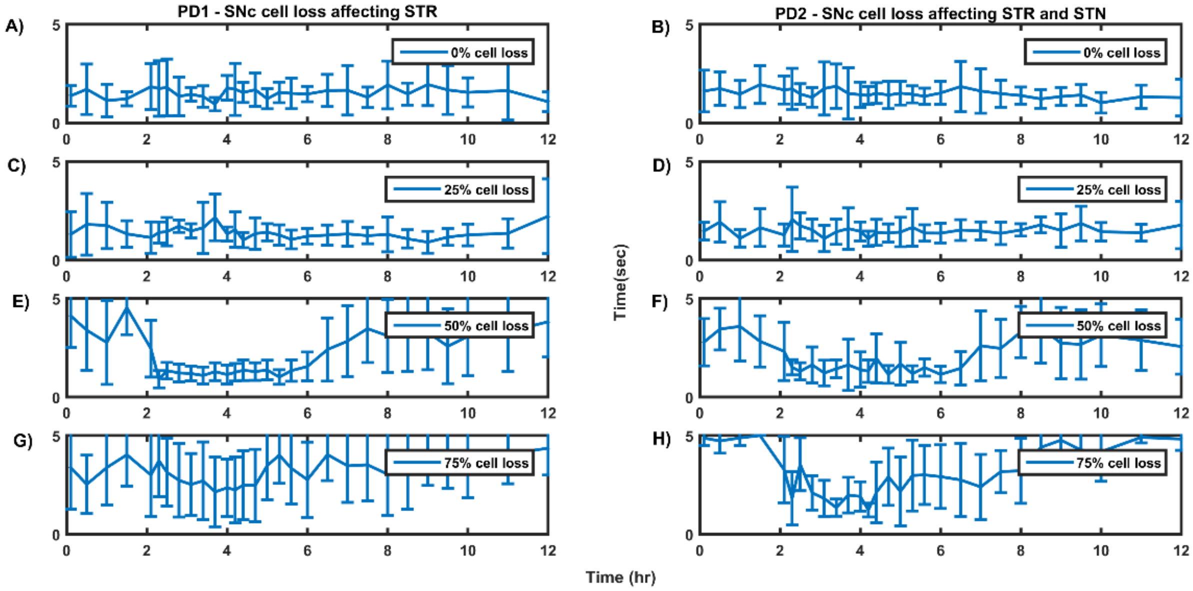


*Figure S7.11: The figure shows the average time to reach the target measured across 10 trials for varying percentage of cell loss impacting only the striatum (A, C, E, G) and impacting both striatum and STN (B, D, F, H). The performance metrics is observed for the L-DOPA dosage of 200mg and reaching performance measured at various times during the 12-hour window where the L-DOPA drug is administered.*

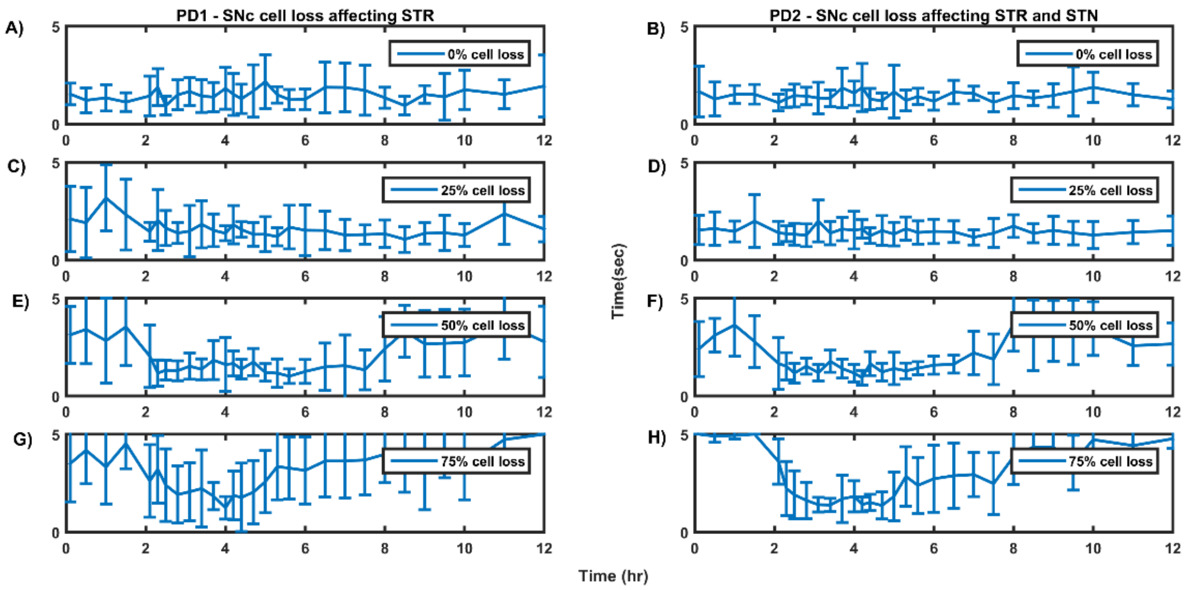


*Figure S7.12: The figure shows the average time to reach the target measured across 10 trials for varying percentage of cell loss impacting only the striatum (A, C, E, G) and impacting both striatum and STN (B, D, F, H). The performance metrics is observed for the L-DOPA dosage of 300mg and reaching performance measured at various times during the 12-hour window where the L-DOPA drug is administered*

c. A consolidated view of the therapeutic effect

Figure S7.13 below provides a consolidated view of the performance across the dosages and the percentage cell loss. Here we divided the performance observed into reach into three categories based on the average time to reach. i) Normal (represented by the green shaded region) ii) Intermediate (represented by the orange shaded region), and iii) Severe (represented by the red shaded region). The upper threshold for the normal region is 2 sec, and the upper and lower thresholds for the intermediate region are 4 sec and 2 sec, respectively. The average time to reach above 4 sec falls into the severe region. The length of therapeutic effect window is calculated by taking the time difference between the points when the performance characteristics entered into the green shaded region (normal) until it started wearing off and crosses back to the orange shaded region (intermediate).

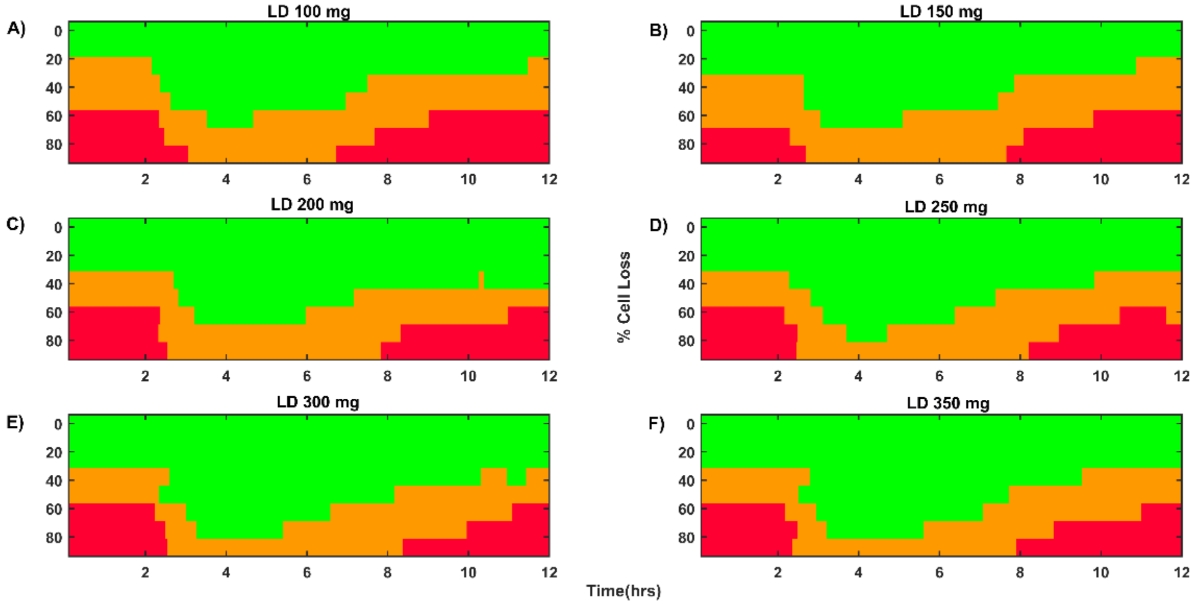


*Figure S7.13: The figure shows the average time to reach the target measured across 10 trials. The overall task performance is divided into three categories (green, orange, and red). The brown line represents the actual recorded reaching time, and the blue line is the smoothened curve after interpolation. Performance metrics is observed for the L-DOPA dosage of 50 mg and reaching performance measured at various times during the 12-hour window where the L-DOPA drug is administered*

### S8. Differential Axonal Degeneration Leading to Various PD Motor Symptoms

This section demonstrates how the nigrostriatal dopaminergic loss and the nigrosubthalamic axonal denervation manifest into various PD symptoms. First, we show the performance without any cell loss, and then we look into the cases where the progressive cell loss impacts the reaching performance. Here we broadly classify the performance evaluation into three categories, a) without any cell loss, b) PD1- with cell loss affecting striatum alone and c) PD2- with cell loss affecting both striatum and STN.

a. Without any cell loss

With no cell loss, the arm reaches the target very quickly. Figure S8.1(B) shows the distance to target reducing from 0.7 meters to less than 0.1 meters in less than a second. The red dotted line represents the threshold of 0.1 meters, and when the distance to the target becomes less than the threshold, the arm is considered to have reached the target. The movement trajectory is found to be very smooth (Figure S8.1(A)), and the velocity of movement forms a perfect bell curve with speed increased upto a point and then decreasing as the arm approaches the target (Figure S8.1(D)). The dopamine released by SNc neurons in the striatum during the arm reaching peaked at $\sim 264\,nM$ which was in the range of $150 - 400\,nM$ (Schultz, 1998) (Figure S8.1(F)). When the arm was in motion, the synchrony was low (Figure S8.1(G)).

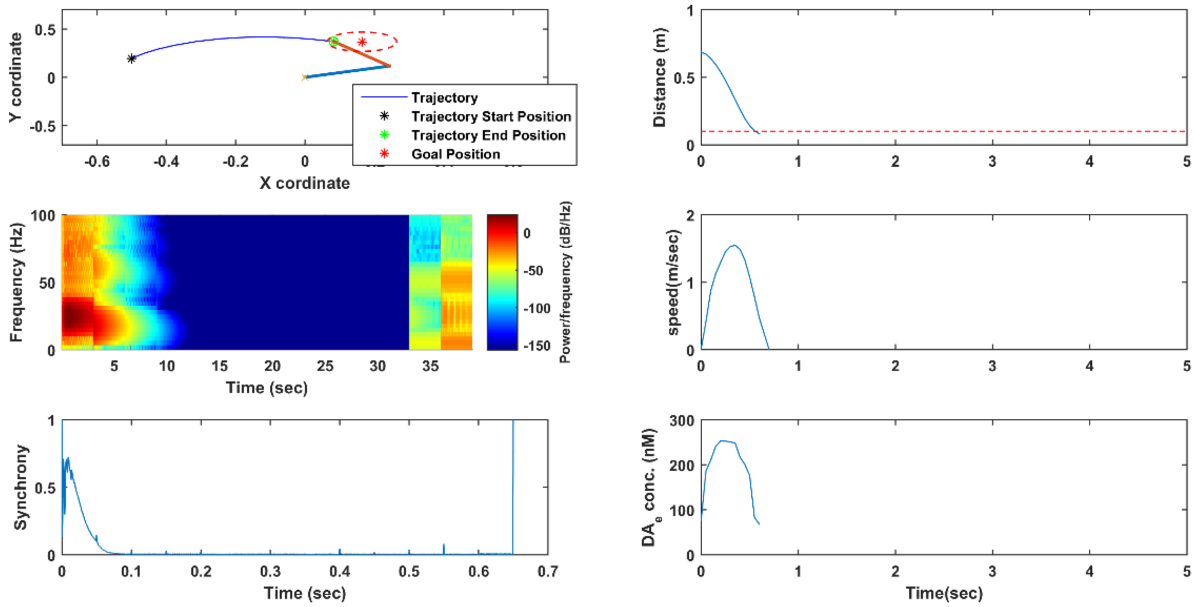


*Figure S8.1: The top left of the figure shows the trajectory followed by the arm end-effector position. The top right plot shows the distance to the target. The middle right figure and the bottom right figure represents the movement speed and the extracellular dopamine released, respectively. The middle left figure represents the time-frequency plot of the STN population activity, and the bottom left figure represents the time-frequency plot of the single STN activity. All these plots are for no cell loss condition.*

b. SNc cell loss impacting striatum alone

With the percentage decline in the dopaminergic neurons in the SNc, the dopamine signal towards the striatum was affected, which resulted in deteriorated performance. With increasing cell loss, the arm took more time reaching the target, and higher percentage of cell loss resulted in tremor-like behavior.

i. With 25% SNc cell loss

When the cell loss was increased to 25%, the arm took more time in reaching the target, and more synchronous activity was observed during movement (Figure S8.2(G)). The spectrogram plot (Figure S8.2(C)) shows higher powers in low-frequency ranges.

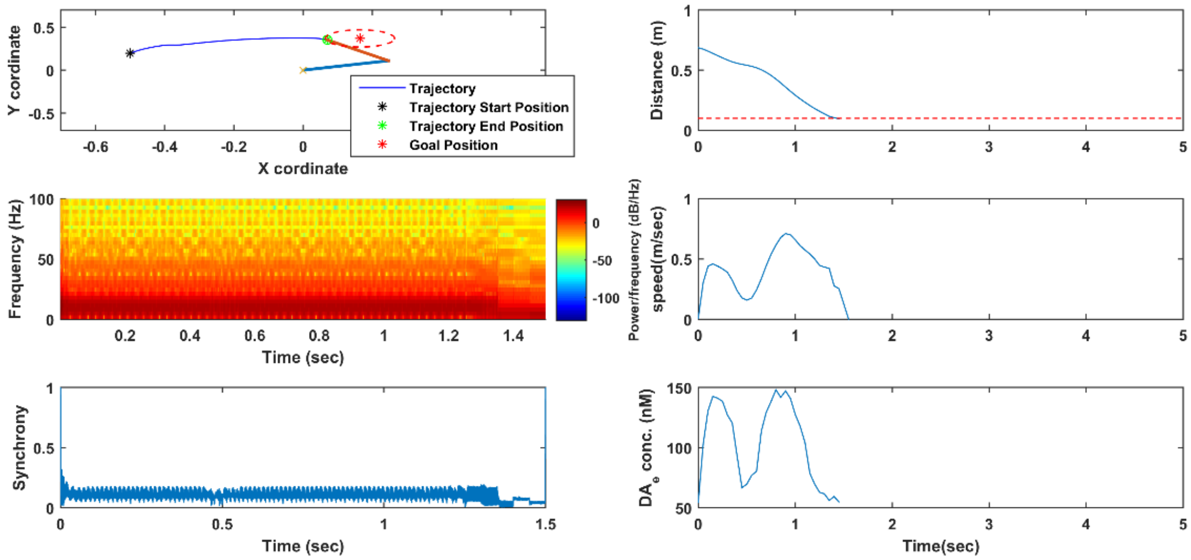


*Figure S8.2: The top left of the figure shows the trajectory followed by the arm end effector position. The top right plot shows the distance to the target. The middle right figure and the bottom right figure represents the movement speed and the extracellular dopamine released, respectively. The middle left figure represents the time-frequency plot of the STN population activity and the bottom left figure represents the time-frequency plot of the single STN activity. All these plots are observed for 25% cell loss condition where the cell loss impacts only the striatum.*

ii. With 50% SNc cell loss

With further increase in cell loss, the time to reach the target increases significantly. With around 50% loss of dopaminergic cells, the time taken to reach the target is considerably more and we can see high powers in slightly higher frequencies around mid-beta region. Synchrony is higher, and the trajectory is not very smooth.

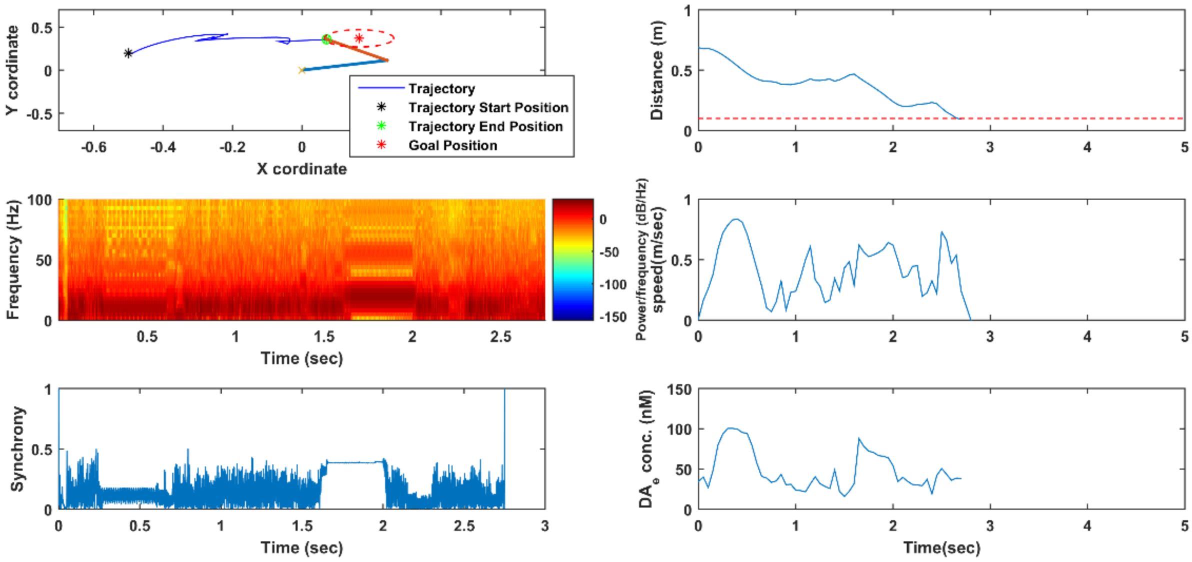


*Figure S8.3: The top left of the figure shows the trajectory followed by the arm end effector position. The top right plot shows the distance to the target. The middle right figure and the bottom right figure represents the movement speed and the extracellular dopamine released respectively. The middle left figure represents the time-frequency plot of the STN population activity and the bottom*

***left figure represents the time-frequency plot of the single STN activity. All these plots are observed for 50% cell loss condition where the cell loss impacts only the striatum.***

iii. With 75% SNc cell loss

With higher cell loss around 75%, the arm hardly reaches the target (Figure S8.4(B)). Tremor-like behavior is observed. The synchrony among the STN neurons is quite high, and the trajectory loses its track as it approaches the target.

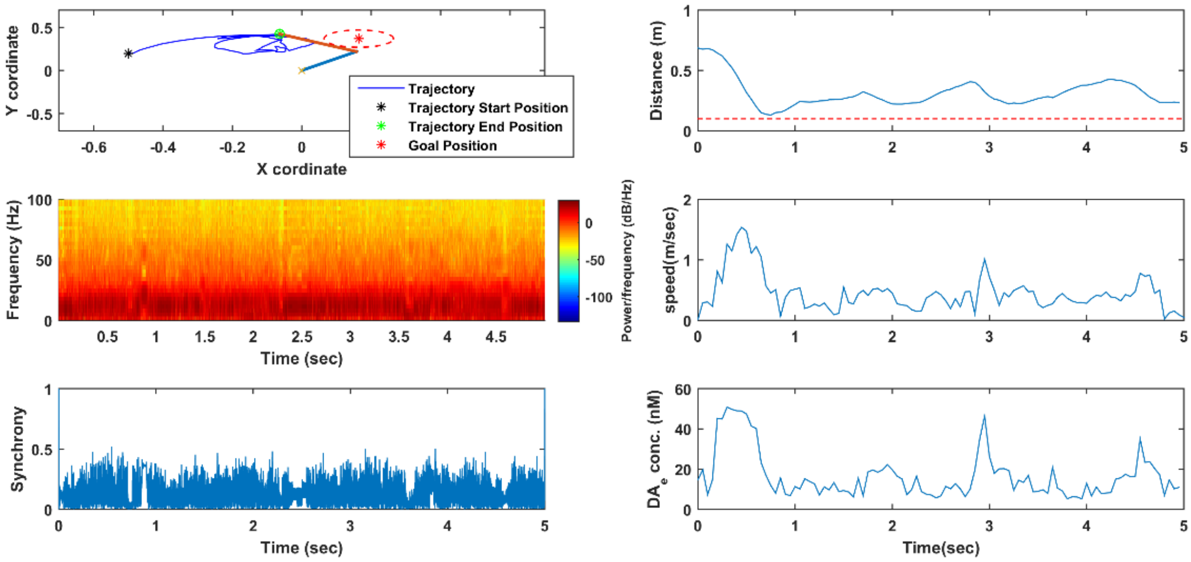


***Figure S8.4: The top left of the figure shows the trajectory followed by the arm end effector position. The top right plot shows the distance to the target. The middle right figure and the bottom right figure represents the movement speed and the extracellular dopamine released respectively. The middle left figure represents the time-frequency plot of the STN population activity and the bottom left figure represents the time-frequency plot of the single STN activity. All these plots are observed for 75% cell loss condition where the cell loss impacts only the striatum.***

c. SNc cell loss impacting both striatum and STN

With dopaminergic cell loss impacting the lateral connections in STN, we observe slowness of movements with increasing cell loss as in the case where the cell loss impacts the striatum alone. However, in this case, when STN lateral connections are also impacted, the behavioral characteristics change from a tremor-like regime towards a rigidity-like regime.

i. With 25% SNc cell loss

With 25% cell loss, the arm reaches the target in just over 1 sec. Synchrony is quite low, and the PD pathology is not noticeably observed.

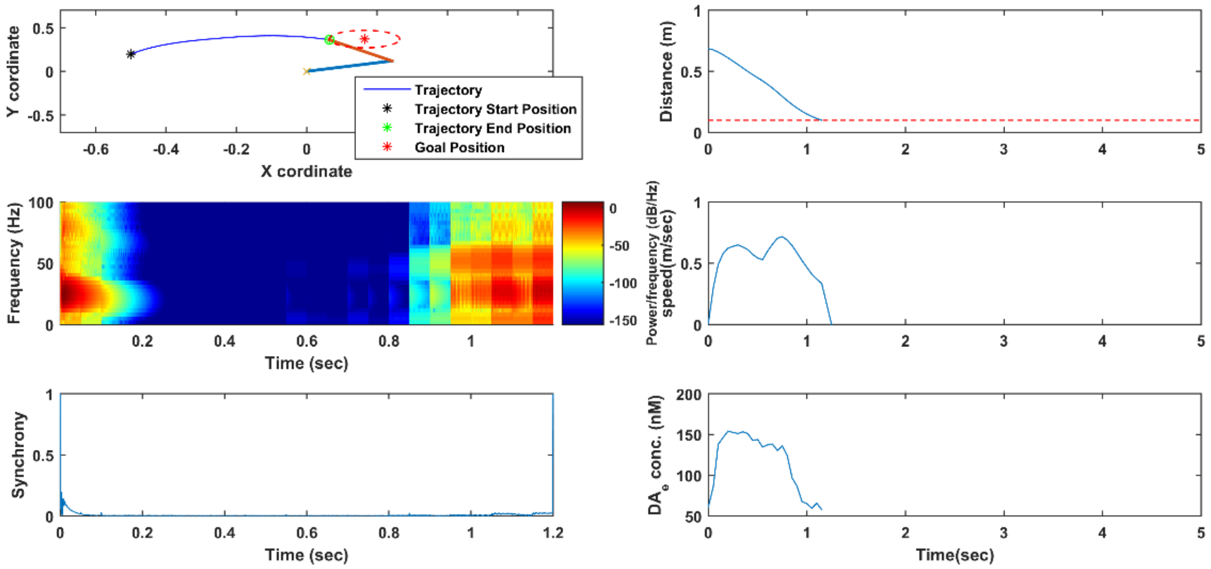


*Figure S8.5: The top left of the figure shows the trajectory followed by the arm end effector position. The top right plot shows the distance to the target. The middle right figure and the bottom right figure represents the movement speed and the extracellular dopamine released respectively. The middle left figure represents the time-frequency plot of the STN population activity and the bottom left figure represents the time-frequency plot of the single STN activity. All these plots are observed for 25% cell loss condition where the cell loss impacts both the striatum and STN.*

ii. With 50% SNc cell loss

With increase in cell loss, say about 50%, the performance deteriorates further. At 50% cell loss, when STN laterals are also impacted in addition to the striatum, the behavior starts to transition from a tremor dominant to a more rigidity dominant regime. We can observe a cogwheel rigidity kind of phenomenon where the arm takes around 0.5 sec to initiate movement (Figure S8.6(D)). At this point of time, we can also see the synchrony to be quite high at the start (Figure S8.6(E)). Once the arm starts picking up the speed, it stops again and resumes after a delay.

iii. With 75% SNc cell loss

At 75% cell loss, the arm hardly moves from its initial position, as seen in the trajectory (Figure S8.7(A)) and the distance to target (Figure S8.7(B)) plots. The synchrony is very high throughout (Figure S8.7(E)), and increased power are observed in gamma regions (Figure S8.7(C)). This resembles a phenomenon like *akinetic rigidity*.

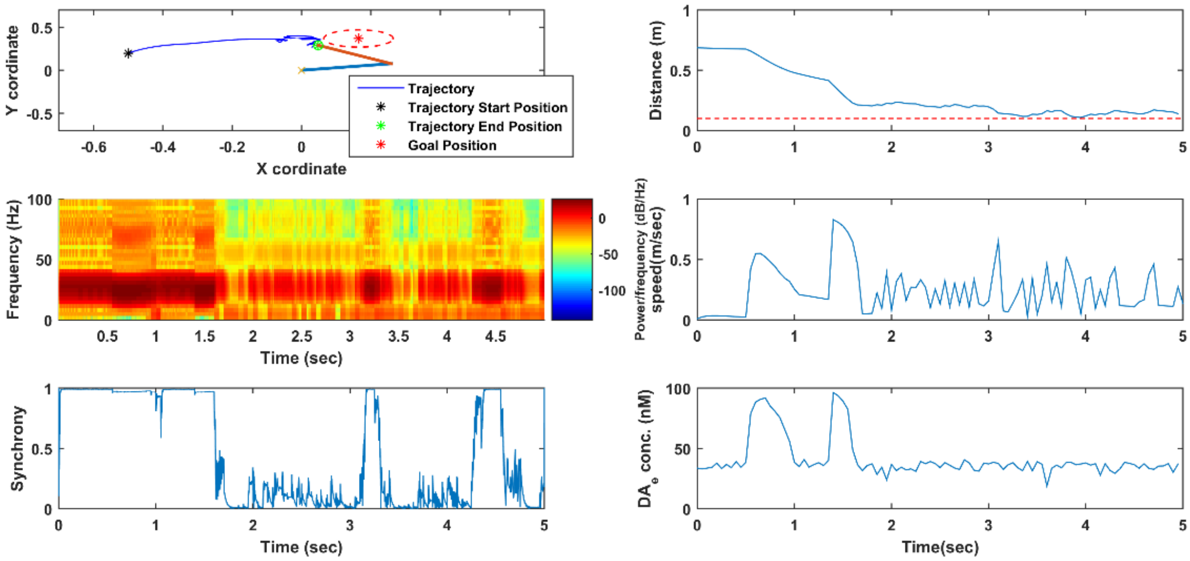


*Figure S8.6: The top left of the figure shows the trajectory followed by the arm end effector position. The top right plot shows the distance to the target. The middle right figure and the bottom right figure represents the movement speed and the extracellular dopamine released respectively. The middle left figure represents the time-frequency plot of the STN population activity and the bottom left figure represents the time-frequency plot of the single STN activity. All these plots are observed for 50% cell loss condition where the cell loss impacts both striatum and STN.*

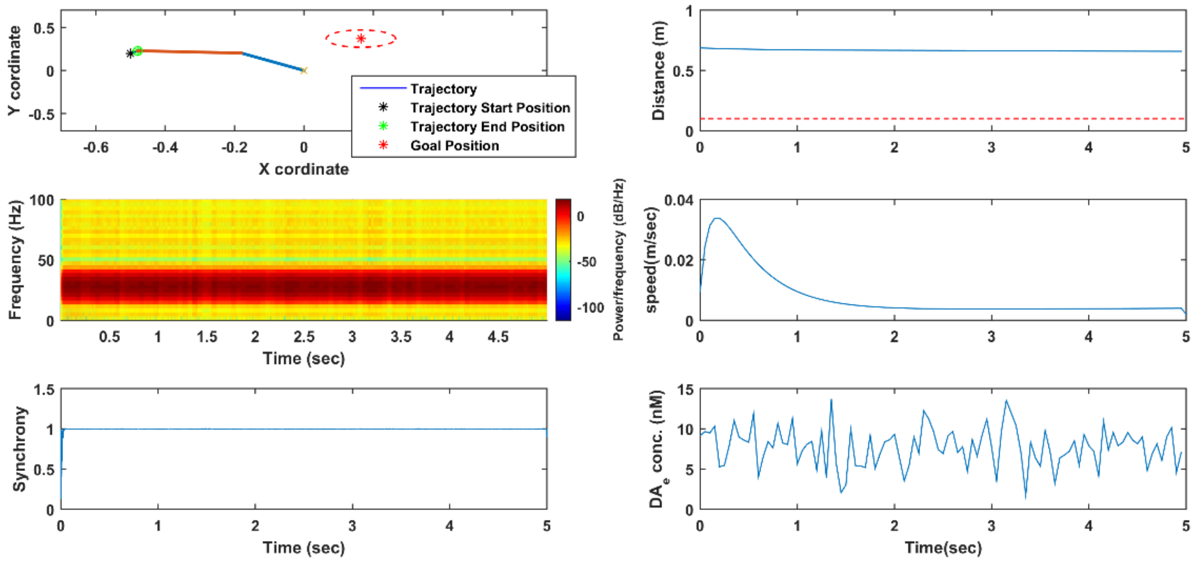


*Figure S8.7: The top left of the figure shows the trajectory followed by the arm end effector position. The top right plot shows the distance to the target. The middle right figure and the bottom right figure represents the movement speed and the extracellular dopamine released respectively. The middle left figure represents the time-frequency plot of the STN population activity and the bottom left figure represents the time-frequency plot of the single STN activity. All these plots are observed for 75% cell loss condition where the cell loss impacts both striatum and STN.*

### S9. STN-GPe Dynamics

The STN-GPe network is modeled using a group of excitatory and inhibitory pair of neurons connected back-to-back that also represents a special type of systems known as the Liénard system (Chakravarthy and Moustafa, 2018). Consider the neuronal pair as shown below.

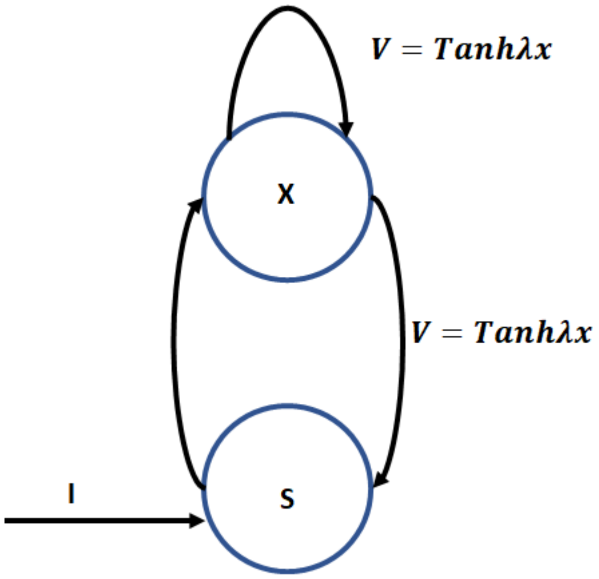


*Figure S9.1: The coupled excitatory-inhibitory pair of STN-GPe neurons where s and x represent GPe and STN neurons, respectively.*

The dynamics of the neuron pair can be described as below

$$\frac{dx}{dt} = -x + V - s \tag{92}$$

$$V = Tanh\lambda x \tag{93}$$

$$\frac{ds}{dt} = -s + V + I \tag{94}$$

$$\frac{d^2x}{dt^2} = -\frac{dx}{dt} + \lambda Sech^2(\lambda x)\frac{dx}{dt} - \frac{ds}{dt} \tag{95}$$

$$\frac{d^2x}{dt^2} = -\frac{dx}{dt} + \lambda Sech^2(\lambda x)\frac{dx}{dt} - V + s - I \tag{96}$$

$$\frac{d^2x}{dt^2} = -\frac{dx}{dt} + \lambda Sech^2(\lambda x)\frac{dx}{dt} - V - I - x + V - \frac{dx}{dt} \tag{97}$$

$$\frac{d^2x}{dt^2} + \frac{dx}{dt}\left(2 - \lambda Sech^2(\lambda x)\right) - (x + I) = 0 \tag{98}$$

Equation 98 is of the form

$\frac{d^2x}{dt^2} + \frac{dx}{dt} f(x) + g(x) = 0$, which represents the generalized form of Liénard system of equations where $f(x) = \lambda Sech^2(\lambda x)$ $and$ $g(x) = -(x + I)$.

STN & GPe neuronal pair is modeled using the same dynamics as given above. A single pair of STN-GPe neurons exhibited the following dynamics as shown in Figures S9.2-S9.5.

a. No external stimulus

With no input current to the neuron pair, the STN and GPe activities exhibit oscillatory behavior (Figure S9.2). As input current varies, the system moves from a periodic oscillatory behaviour to a non-oscillatory behaviour, which can be observed in subsequent Figures.

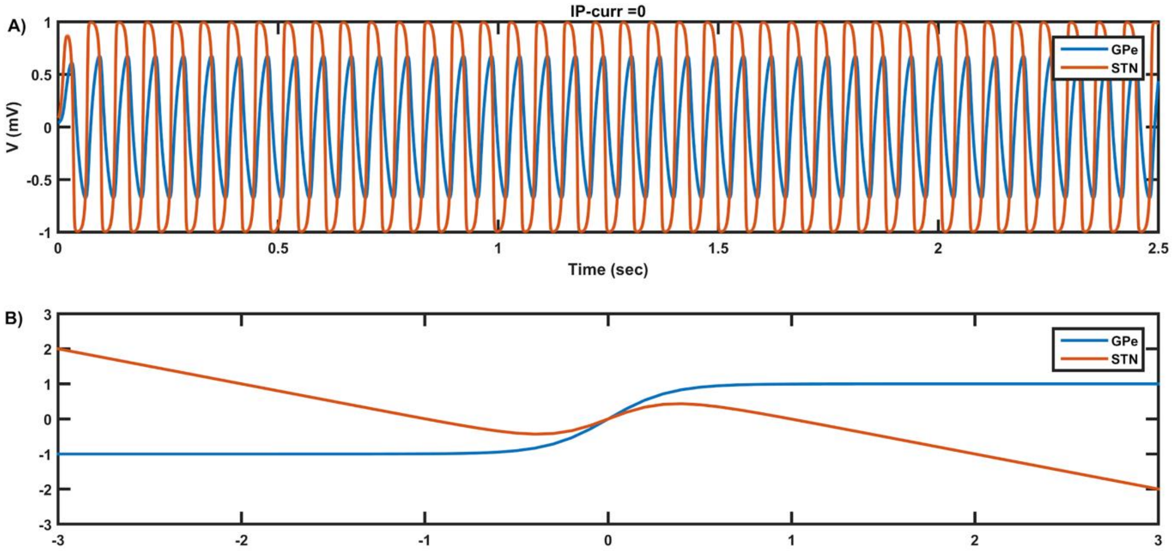


*Figure S9.2: The top portion of the figure shows the activity of STN/GPe neurons over across time. The bottom plot shows the phase plane (blue line represents the GPe nullcline, whereas the red line represents the STN nullcline. The point of intersection of these two nullclines is the fixed point. With no input current, the fixed point is located at the origin. X-axis is x (STN activity, Vstn(t)) and Y-axis is s (GPe activity, Vgpe(t)).*

b. Impact of external stimulus

With external input is applied, the dynamics of the STN and GPe activities transition from periodic to non-periodic. As can be seen in Figures S9.3-S9.5, as the current increases, the frequency of oscillation reduces, and when the current increases beyond 0.34 mA, the oscillations cease (Figure S9.5). Till the current value is increased to 0.34, the system displays oscillatory behavior. However, when the magnitude of the current is increased further, the oscillations cease.

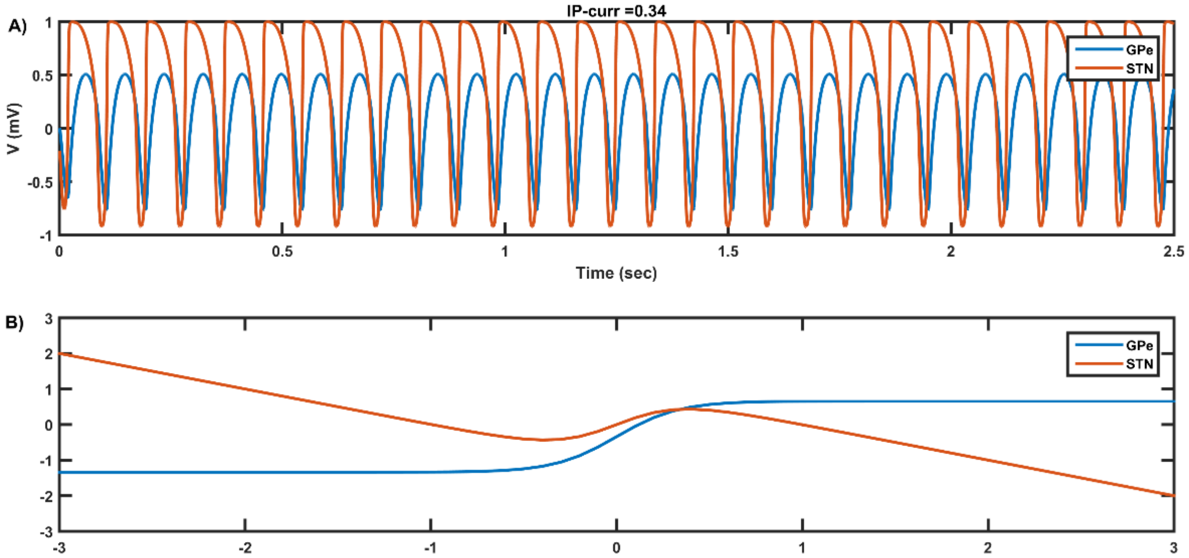


*Figure S9.3: The top portion of the figure shows the activity of STN/GPe neurons over across time. The bottom plot shows the phase plane (blue line represents the GPe nullcline, whereas the red line represents the STN nullcline. The point of intersection of these two nullclines is the fixed point. With a slight increase in the input current, the fixed point is shifted by the magnitude of the current applied and in the direction of the current. X-axis is x (STN activity, Vstn(t)) and Y-axis is s (GPe activity, Vgpe(t)).*

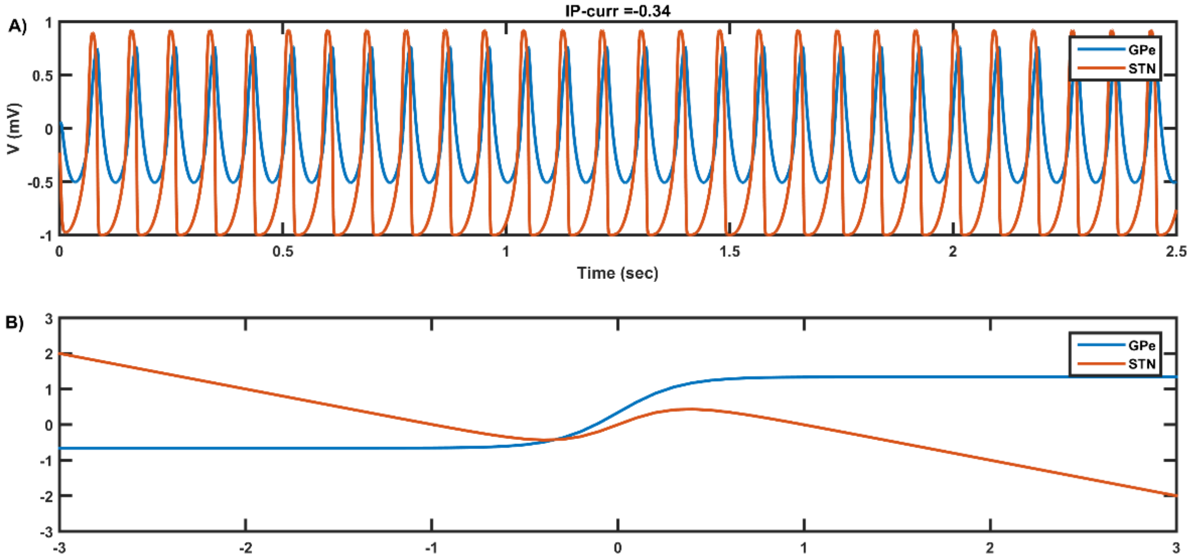


*Figure S9.4: The top portion of the figure shows the activity of STN/GPe neurons over across time. The bottom plot shows the phase plane (blue line represents the GPe nullcline whereas the red line represents the STN nullcline. The point of intersection of these two nullclines is the fixed point. With a slight increase in the input current the fixed point is shifted by the magnitude of the current applied and in the direction of current. Compared to the previous scenario, the current applied here is negative. X-axis is x (STN activity, Vstn(t)) and Y-axis is s (GPe activity, Vgpe(t)).*

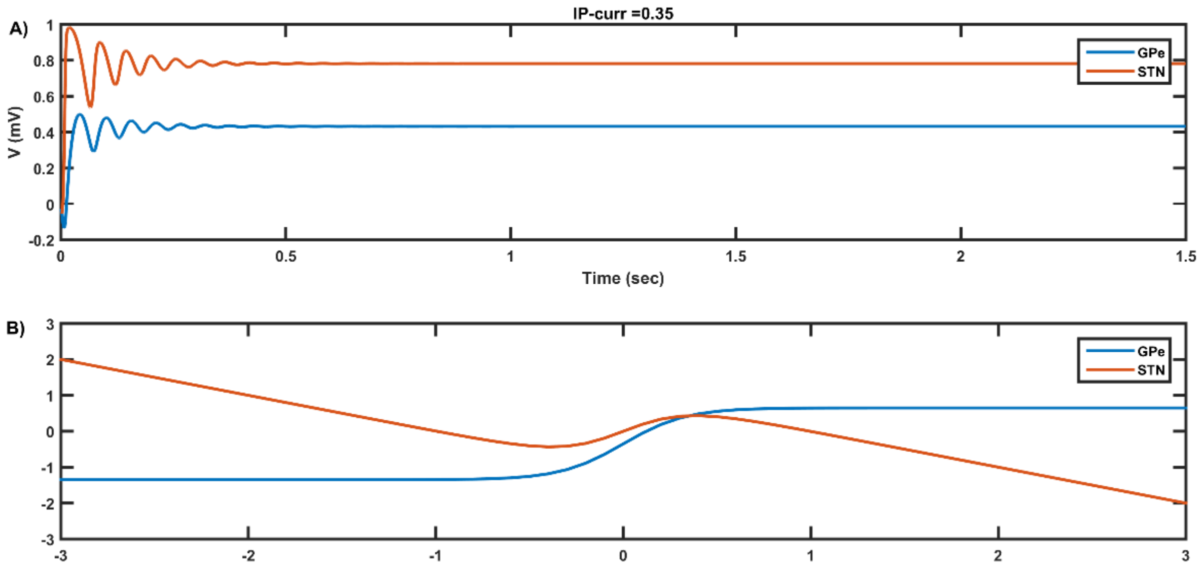


*Figure S9.5: The top portion of the figure shows the activity of STN/GPe neurons over across time. The bottom plot shows the phase plane (blue line represents the GPe nullcline, whereas the red line represents the STN nullcline. The point of intersection of these two nullclines is the fixed point. With further increase in the input current, the behavior changes from periodic to non-periodic. X-axis is x (STN activity, Vstn(t)) and Y-axis is s (GPe activity, Vgpe(t)).*

c. Impact of interconnectivity weights

However, by controlling the interconnectivity weights between the two neurons, we can bring the regime back to the oscillatory mode with the increase in weights. Figure S9.6 and S9.7 shows the impact of increasing the interconnectivity weights between STN and GPe ($W_{sg}$). Also, the frequency of oscillations keeps increasing as $W_{sg}$ is increased (Figure S9.8).

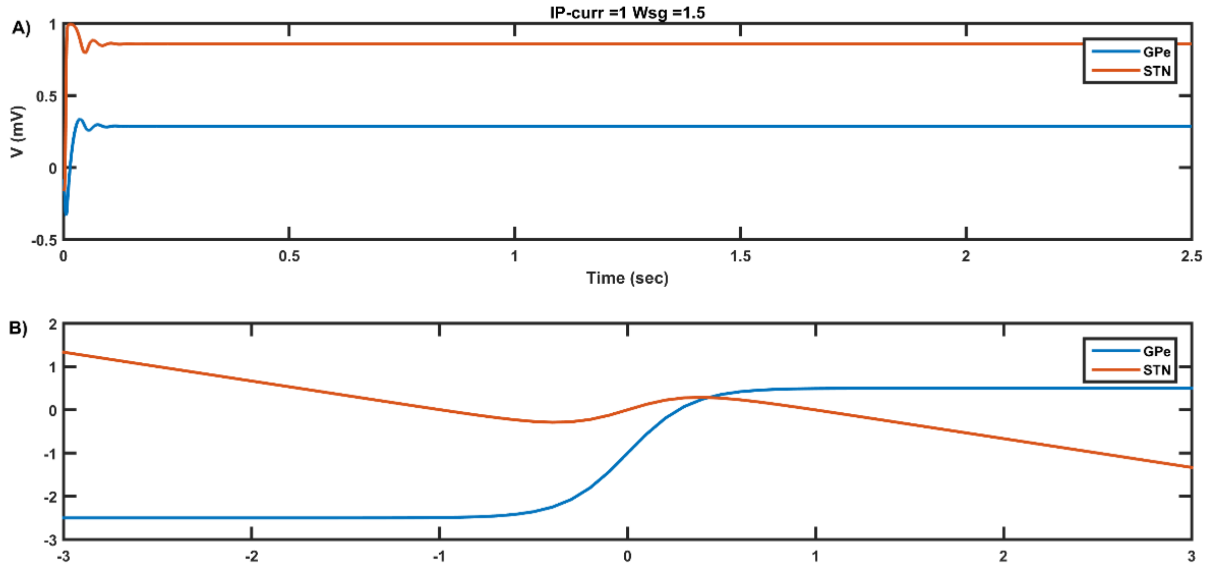


*Figure S9.6: The top portion of the figure shows the activity of STN/GPe neurons over across time. The bottom plot shows the phase plane (blue line represents the GPe nullcline whereas the red line*

***represents the STN nullcline. Increasing the interconnectivity weights results in change in behavioral regime. X-axis is x (STN activity, Vstn(t)) and Y-axis is s (GPe activity, Vgpe(t)).***

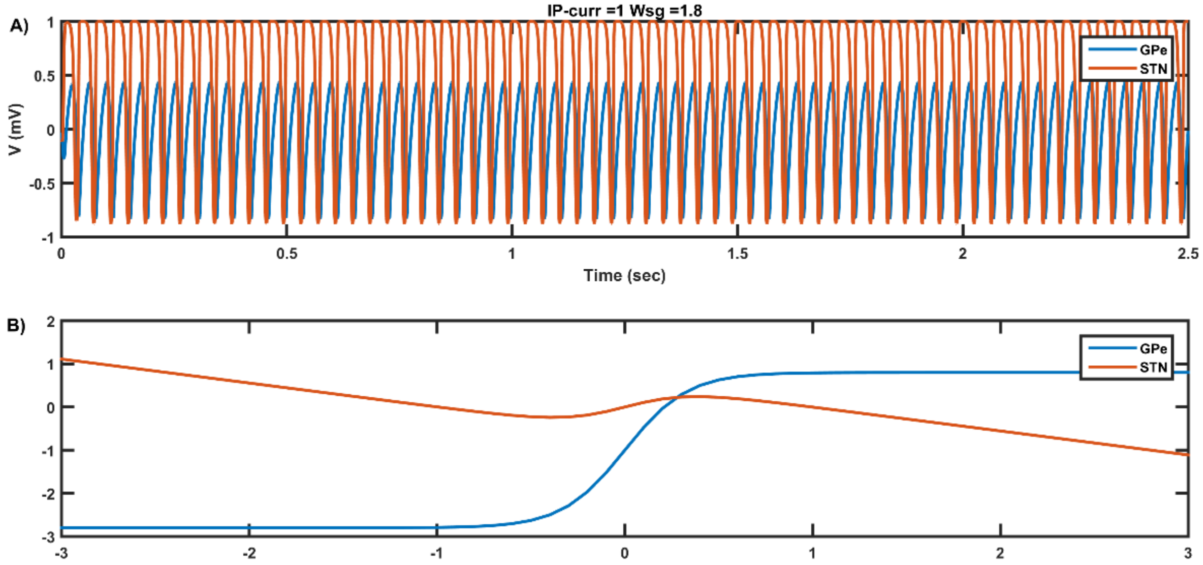


***Figure S9.7: The top portion of the figure shows the activity of STN/GPe neurons over across time. The bottom plot shows the phase plane (blue line represents the GPe nullcline whereas the red line represents the STN nullcline. Increasing the interconnectivity weights results in change in behavioral regime. X-axis is x (STN activity, Vstn(t)) and Y-axis is s (GPe activity, Vgpe(t)).***

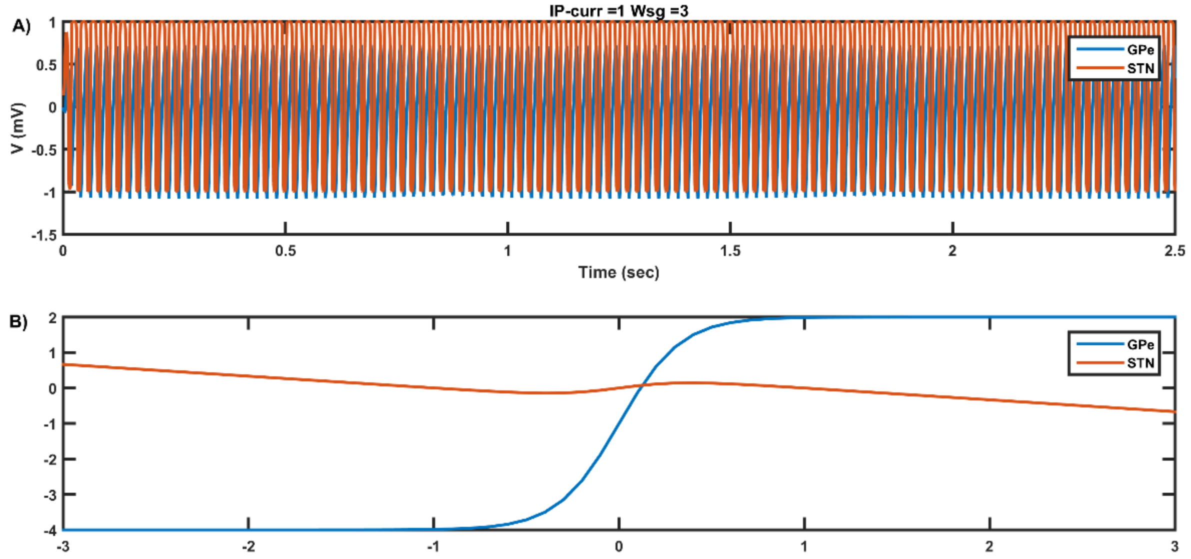


***Figure S9.8: The top portion of the figure shows the activity of STN/GPe neurons over across time. The bottom plot shows the phase plane (blue line represents the GPe nullcline whereas the red line represents the STN nullcline. Increasing the interconnectivity weights results in change in behavioral regime. As the interconnectivity weights are increased, the frequency of oscillations is also found to be increased. X-axis is x (STN activity, Vstn(t)) and Y-axis is s (GPe activity, Vgpe(t)).***

The increase in the frequency of oscillations with increasing interconnectivity weights is illustrated in the frequency spectrum plots shown below in Figure S9.9. With a constant stimulus of ID2 =0.5 mA, the peak frequency is shifted from 25 Hz to 35 Hz, when $W_{sg}$ is changed from 1.5 to 2.

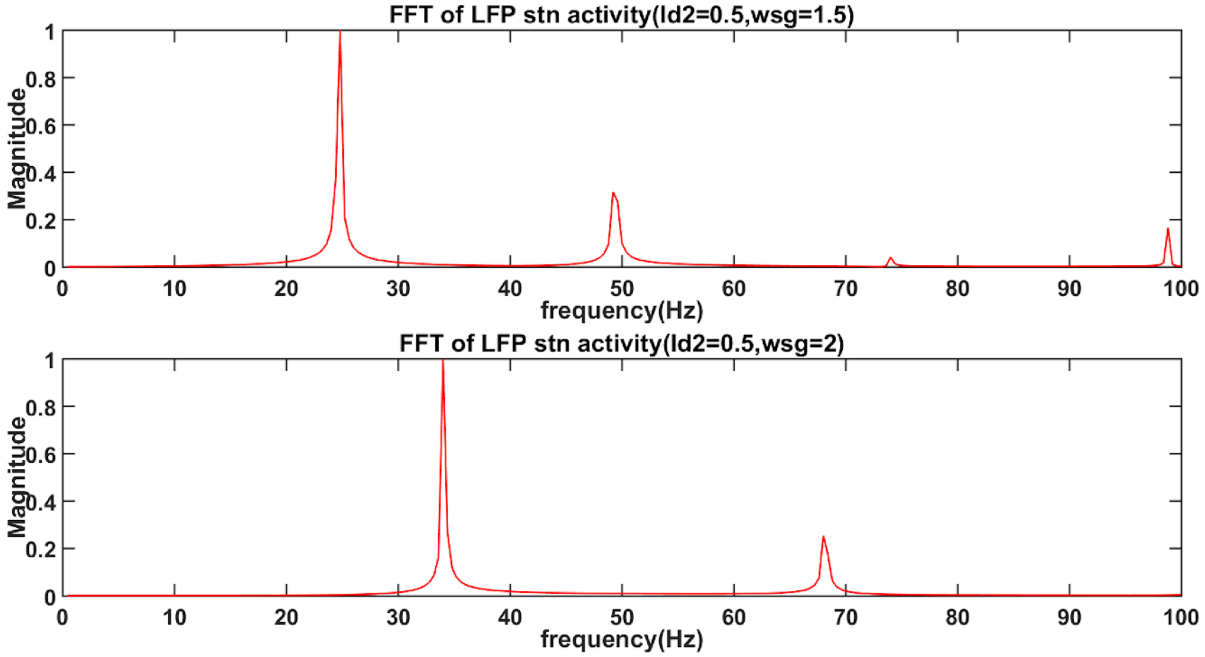


*Figure S9.9: The top portion of the figure shows the FFT for the case where the interconnectivity weight, wsg=1.5. With increase in wsg=2, the peak frequency is shifted to a higher range.*

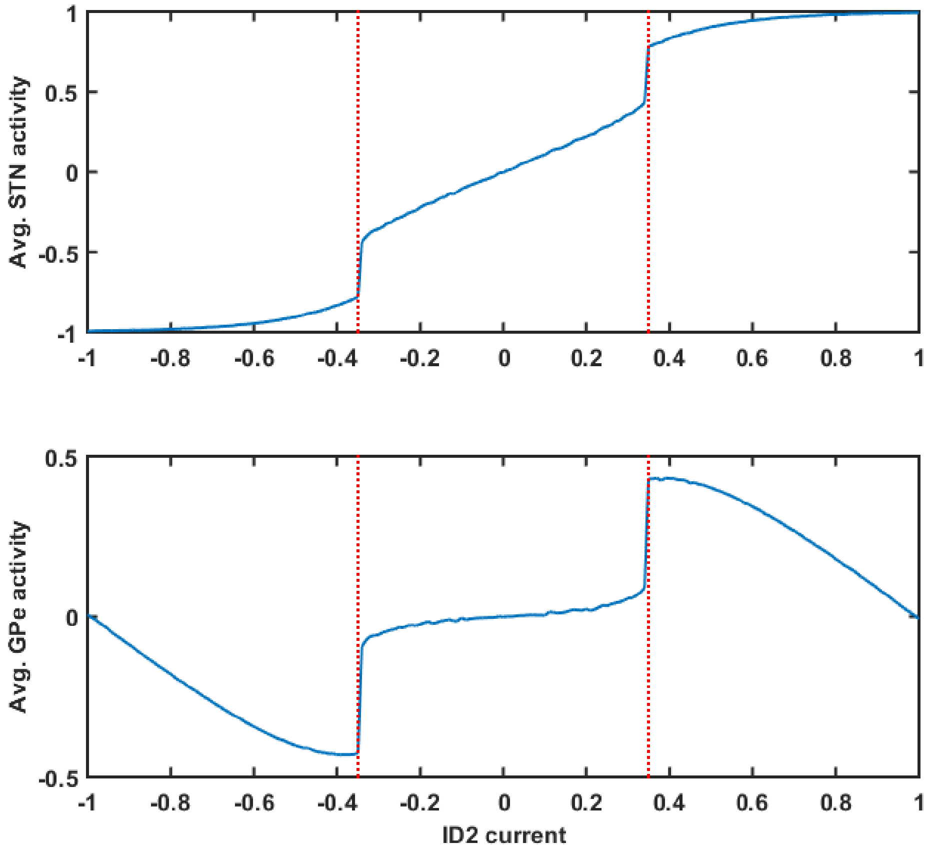

*Figure S9.10: The top portion of the figure shows the average activity of STN neurons over across different values of input stimulus. The bottom plot shows the average activity of GPe neurons over across different values of input stimulus.*

The relationship between the input current and oscillatory behaviour as shown in Figure S9.10 below. As seen for intermediate values of the current, the system exhibits oscillatory behaviour.

Similarly, for a 2D sheet of neurons, the STN neurons have a one-to-one connection to the GPe neurons. Also, the neurons of both the STN and GPe subpopulations are connected among themselves via lateral weights of uniform strength, $\epsilon_s$ and $\epsilon_g$, respectively. The factors influencing the neuronal dynamics are the input current, the interconnectivity weights and the lateral connection strengths, $\epsilon_s$ and $\epsilon_g$. Figure S9.11-S9.13 shows the probability of oscillations (computed according to section S10.b) for the STN and GPe neurons, which is controlled by the input current, lateral weights, and the interconnectivity strength.

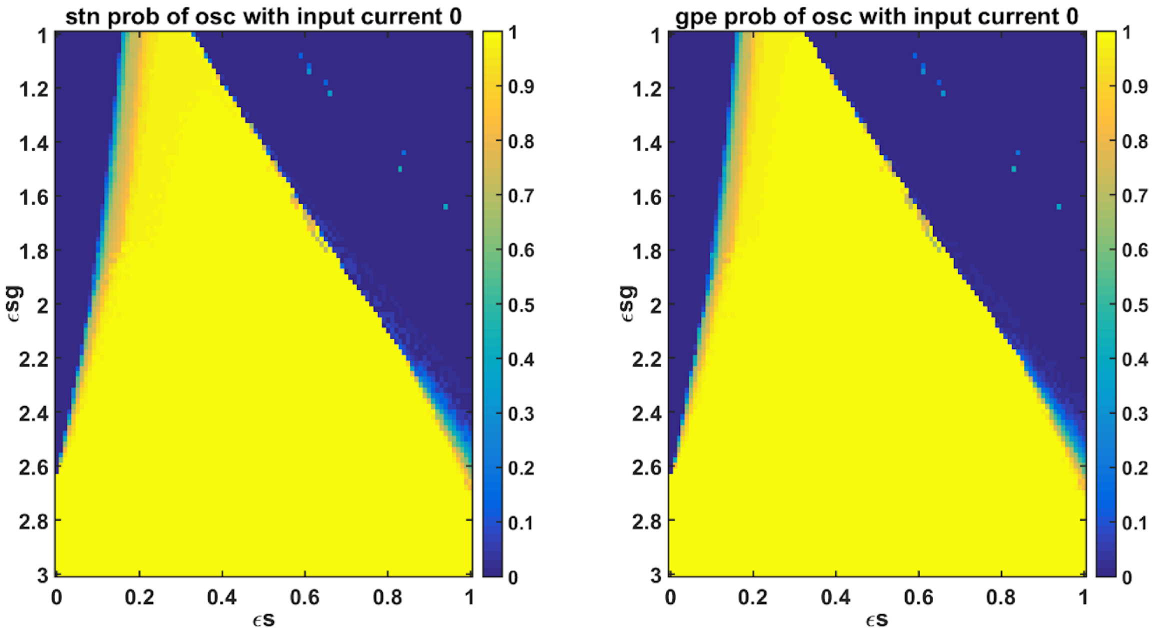


*Figure S9.11: The left portion of the figure shows the probability of oscillations for the STN population over different ranges of $\epsilon_{sg}$ and $\epsilon_s$. The left portion of the figure shows the probability of oscillations for the GPe population over different ranges of $\epsilon_{sg}$ and $\epsilon_s$.*

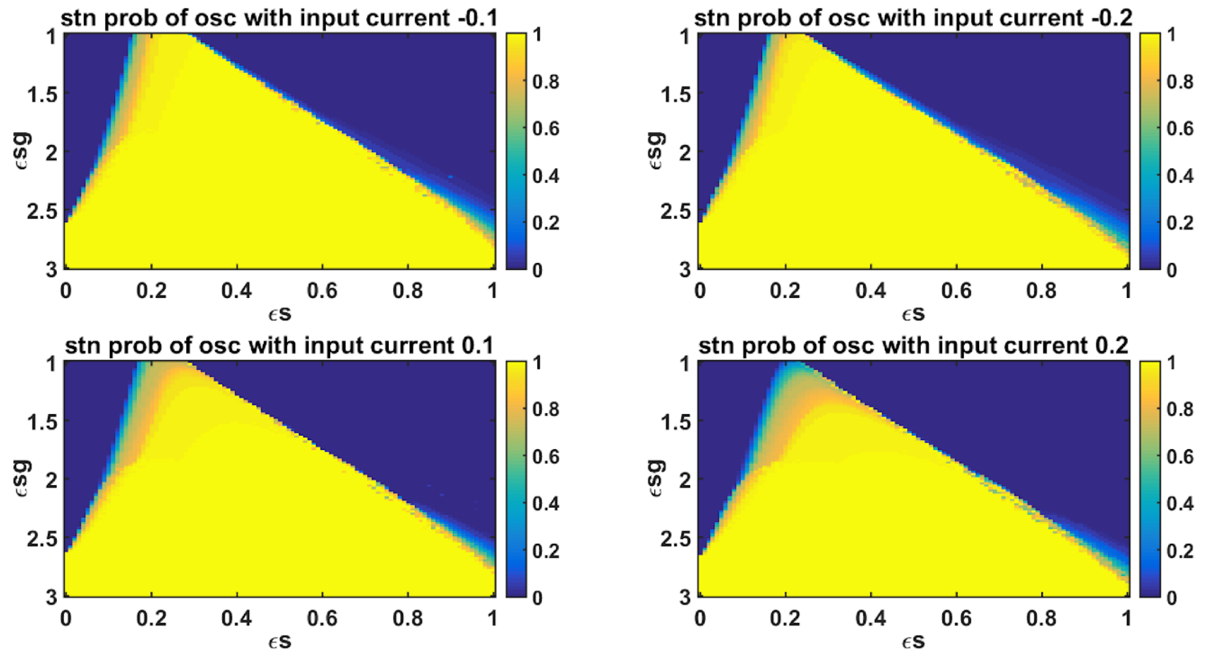


*Figure S9.12: The left portion of the figure shows the probability of oscillations for the STN population with input currents -0.1 and 0.1 over different ranges of $\epsilon_{sg}$ and $\epsilon_s$. The right portion of the figure shows the probability of oscillations for the GPe population with input currents -0.1 and 0.1 over different ranges of $\epsilon_{sg}$ and $\epsilon_s$.*

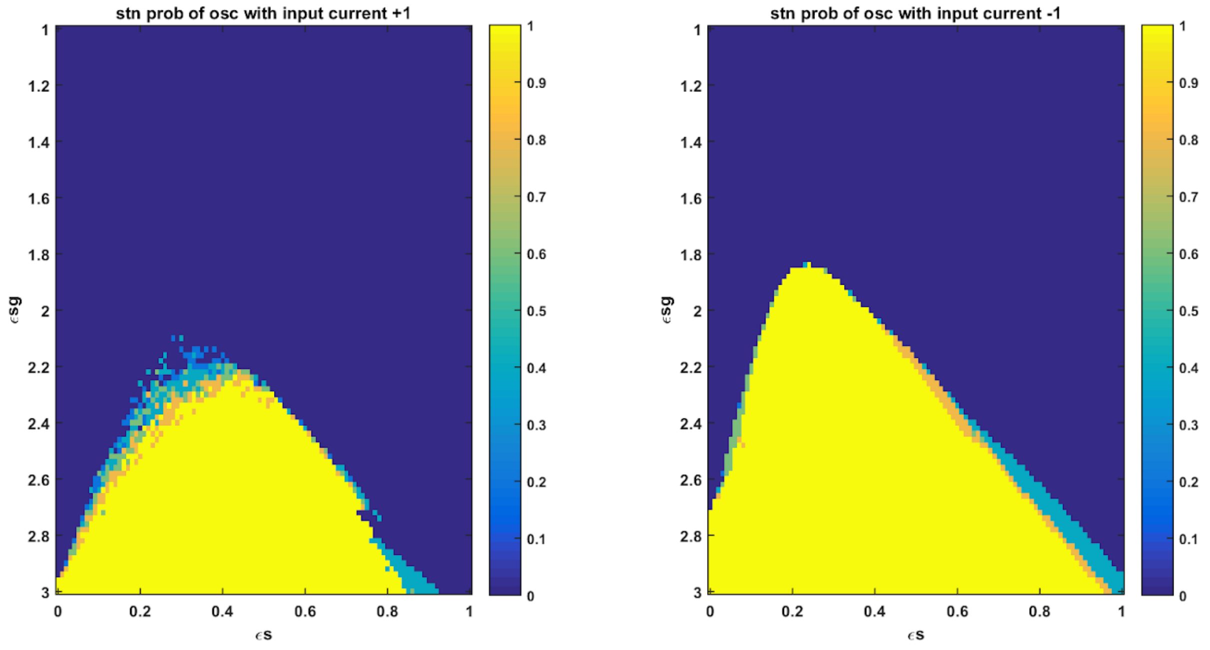


*Figure S9.13: The left portion of the figure shows the probability of oscillations for the STN population with input current = +1mA over different ranges of $\epsilon_{sg}$ and $\epsilon_s$. The right portion of the figure shows the probability of oscillations for the GPe population with input currents =-1 mA over different ranges of $\epsilon_{sg}$ and $\epsilon_s$.*

We can see from the above plots that the probability of oscillations varies with variations in the input current. Also, the lateral STN weights and the STN-GPe interconnectivity weights modulate the probability of oscillations.

Hence in our model, we have tuned both the STN-GPe interconnectivity weights and the STN Lateral weights so as to get the desired mode of operation. Controlling these parameters, we can make the STN-GPe network more complex or less complex and, as seen earlier, reduced complexity results in pathological behavior.

**S10. Network Analysis**

a. Synchronization

Neuronal synchronization is the measure of synchronicity (high synchrony - almost all neurons firing at once, low synchrony - least number of neurons firing at once) in the population of neurons within a network. We had quantified the synchrony in the population of neurons at time $t$ by following equation (Pinsky and Rinzel, 1995; Muddapu et al., 2019),

$$R_x(t) = \frac{1}{N * e^{i*\theta(t)}} \sum_{j=1}^{N} e^{i*\phi_j(t)} \tag{99}$$

$$\phi_j(t) = 2 * \pi * \frac{(T_{j,k} - t_{j,k})}{(t_{j,k+1} - t_{j,k})} \tag{100}$$

where, $R_x(t)$ is the instantaneous synchronization measure ($0 \leq R_x(t) < 1$), $x$ being GPe or STN neuron, $N$ is the number of neurons in the network, $\theta(t)$ is the instantaneous average phase of neurons, $\phi_j(t)$ is the instantaneous phase of jth neuron, $t_{j,k}$ and $t_{j,k+1}$ are the spike times of kth and (k+1)th spike of jth neuron, respectively, $T_{j,k} \in [t_{j,k}, t_{j,k+1}]$.

b. Probability of Oscillations

If the standard deviation of STN and GPe activity during the time interval between 1.5 sec ($t_1 = 1501$) to 2.5 sec ($t_n = 2500$) is greater than 0.2, then it is labeled to show oscillatory activity (Chakravarthy and Moustafa, 2018). This is run over multiple iterations ($Num$), and probability of oscillation ($P_{osc}$) is calculated for both STN and GPe as,

$$x_{mean}^{c} = \frac{1}{t_n - t_1} \sum_{t_1}^{t_n} x_c(t_i) \tag{101}$$

$$x_{std}^{c} = \sqrt{\frac{\sum_{t_1}^{t_n} (x_c(t_i) - x_{mean}^{c})^2}{t_n - t_1}} \tag{102}$$

$$x_{std}^{c}(k) > 0.2 \quad then, \quad OSC_c + 1, \quad where\ k = 1, 2 \ldots Num \tag{103}$$

$$P_{osc}^{c} = \frac{OSC_c}{Num} \tag{104}$$

where, $x_{mean}^{c}$ is the average activity, $x_{std}^{c}$ is the standard deviation, $OSC_c$ is the oscillation count, $P_{osc}^{c}$ is the probability of oscillation ($c = STN\ or\ GPe$).

**References**


Anzalone, A., Lizardi-Ortiz, J. E., Ramos, M., De Mei, C., Hopf, F. W., Iaccarino, C., et al. (2012). Dual control of dopamine synthesis and release by presynaptic and postsynaptic dopamine D2 receptors. *J. Neurosci.* 32, 9023–9024. doi:10.1523/JNEUROSCI.0918-12.2012.

Baston, C., Contin, M., Calandra Buonaura, G., Cortelli, P., and Ursino, M. (2016). A Mathematical Model of Levodopa Medication Effect on Basal Ganglia in Parkinson’s Disease: An Application to the Alternate Finger Tapping Task. *Front. Hum. Neurosci.* 10. doi:10.3389/fnhum.2016.00280.

Best, J. A., Nijhout, H. F., and Reed, M. C. (2009). Homeostatic mechanisms in dopamine synthesis and release: a mathematical model. *Theor. Biol. Med. Model.* 6, 21. doi:10.1186/1742-4682-6-21.

Chakravarthy, V. S., and Moustafa, A. A. (2018). *Computational Neuroscience Models of the Basal Ganglia*. Singapore: Springer Singapore doi:10.1007/978-981-10-8494-2.

Chen, R., Wei, J., Fowler, S. C., and Wu, J.-Y. (2003). Demonstration of functional coupling between dopamine synthesis and its packaging into synaptic vesicles. *J. Biomed. Sci.* 10, 774–781. doi:10.1007/BF02256330.

Ford, C. P. (2014). The role of D2-autoreceptors in regulating dopamine neuron activity and transmission. *Neuroscience* 282, 13–22. doi:10.1016/j.neuroscience.2014.01.025.

Izawa, J., Kondo, T., and Ito, K. (2004). Biological arm motion through reinforcement learning. *Biol. Cybern.* 91, 10–22. doi:10.1007/s00422-004-0485-3.

Muddapu, V. R., and Chakravarthy, V. S. (2021). Influence of Energy Deficiency on the Subcellular Processes of Substantia Nigra Pars Compacta Cell for Understanding Parkinsonian Neurodegeneration. *Sci. Rep.* 11, 1754. doi:10.1038/s41598-021-81185-9.

Muddapu, V. R., Mandali, A., Chakravarthy, V. S., and Ramaswamy, S. (2019). A Computational Model of Loss of Dopaminergic Cells in Parkinson's Disease Due to Glutamate-Induced Excitotoxicity. *Front. Neural Circuits* 13, 11. doi:10.3389/fncir.2019.00011.

Pinsky, P. F., and Rinzel, J. (1995). Synchrony measures for biological neural networks. *Biol. Cybern.* 73, 129–137. doi:10.1007/BF00204051.

Reed, M. C., Nijhout, H. F., and Best, J. A. (2012). Mathematical Insights into the Effects of Levodopa. *Front. Integr. Neurosci.* 6, 1–24. doi:10.3389/fnint.2012.00021.

Schultz, W. (1998). Predictive Reward Signal of Dopamine Neurons. *J. Neurophysiol.* 80, 1–27. doi:10.1152/jn.1998.80.1.1.

Venda, L. L., Cragg, S. J., Buchman, V. L., and Wade-Martins, R. (2010). α-Synuclein and dopamine at the crossroads of Parkinson's disease. *Trends Neurosci.* 33, 559–568. doi:10.1016/j.tins.2010.09.004.

Véronneau-Veilleux, F., Ursino, M., Robaey, P., Lévesque, D., and Nekka, F. (2020). Nonlinear pharmacodynamics of levodopa through Parkinson's disease progression. *Chaos* 30, 093146. doi:10.1063/5.0014800.

Zadravec, M., and Matjačić, Z. (2013). Planar arm movement trajectory formation: An optimization based simulation study. *Biocybern. Biomed. Eng.* 33, 106–117. doi:10.1016/j.bbe.2013.03.006.
